# MutQA: A Cross-Validated Q&A Dataset for Genetic Mutations

**DOI:** 10.64898/2026.10.27.762236

**Authors:** Robert McCourt, David Arredondo, Oladimeji Macaulay, Shrey Poshiya, Oluoma Edeh, Kushal Virupakshappa, My Nguyen Bach, Enrique Almeida Ruiz, Yue Hu, Luis E. Tafoya, Debjani P. Hudgens, Deepali Kundnani, Avinash Sahu

## Abstract

We introduce MUTQA, a dataset of 330,387 citation-grounded question–answer (QA) pairs linking genetic mutations to their functional consequences as described in the biomedical literature. Each record pairs a protein-coding mutation with a natural-language question and an evidence-backed answer anchored to sentences in one of 27,442 open-access PubMed Central articles, spanning 6,620 genes and 125,593 gene–variant pairs. The dataset is constructed using CROSSVALQA, a framework that enforces *information isolation* between independent LLMs: one model generates QA pairs from an article, a second model answers the same question without access to the first model’s response, and a third model judges whether each model’s rationale supports the other’s answer, so that retained QA pairs are reproducible from the source text rather than hallucinated. Of 401,406 audited QA pairs, 82.7% pass both directions of this cross-validation; those with a recoverable reference protein sequence form the released benchmark. In a 999-pair stratified audit, two expert annotators agree exactly on 86.4% of cross-validated QA pairs (95% CI 83.7–89.0; *n*=638), a tier in which about 90% of ratings are “correct”. Individual LLM judges agree with individual annotators at a comparable 87.0% on this tier but far less on QA pairs that fail cross-validation. Fine-tuning MutaPLM on MutQA nearly doubles its mean LLM-judge score for question-conditioned mutation-effect answering (1.45 to 2.82 on a 0–5 scale) and improves mutation-recovery accuracy from effect descriptions from 19.4% to 32.1%. Providing variant-matched MutQA retrieval context, without fine-tuning, improves strict answer accuracy across four LLM families by 20.0–28.6 percentage points, demonstrating framework ability to improve reliability LLM answers. We also apply CrossValQA to proteomics, providing a proof of concept for its use beyond mutation-specific QA (Appendix G). We release the dataset, construction framework, and evaluation code.

## 1 Introduction

Instruction tuning has enabled large language models to follow directives [Ouyang et al., 2022, Zheng et al., 2023, Dubey et al., 2024], but adapting them to scientific domains requires *multimodal supervision*: training instances that link structured objects—protein sequences, compounds, medical images, molecular graphs, or genetic variants—with natural-language instructions and responses. The bottleneck is not data volume but the scarcity of such linked pairs. A mutation identifier, for instance, becomes useful for supervision only when linked with a question naming that variant and an answer grounded in experimental evidence—alignments that exist across millions of publications, but only as co-occurrences in prose rather than as machine-readable instruction–response pairs [Wu et al., 2023].

One response is to generate QA pairs with LLMs—Self-Instruct [Wang et al., 2023], Alpaca [Taori et al., 2023], Mol-Instructions [Fang et al., 2024], and MolInstruct [Cao et al., 2023] have produced instruction-tuning pairs at low cost—but generated QA introduces two failure modes that defeat the link to grounded evidence: the generator hallucinates [Ji et al., 2023], and when it substitutes parametric knowledge for the source text, the link between answer and document is severed.

A second response derives QA pairs from structured biological databases such as UniProt [Consortium, 2022], GO [Harris et al., 2004], or the PDB [Sussman et al., 1998]: curated facts can be linked to questions without hallucination, but the link is constrained by the database schema, exemplified by MutaDescribe [Luo et al., 2024] applying the same single-question template (“Please generate a detailed text to describe its effect.”) to every mutation. Both responses omit the experimental conditions and mechanistic reasoning that appear only in the primary literature; Table 2 compares the resources of each kind.

Both responses fail at the same point: neither can guarantee that an answer is genuinely linked to the primary literature. We address this with **CrossValQA**, a framework that generates QA pairs from unstructured text under *information isolation* and *bidirectional cross-validation*. Two LLMs from different providers independently answer the same question from the source text; a third model checks whether each model’s rationale supports the other’s answer; QA pairs that pass in both directions are labelled *cross-validated* and form the released benchmark, while partially validated and non-validated QA pairs are released with their tier labels for analysis. The pipeline is itself an evaluation methodology whose assumptions and failure modes we characterize here.

### Contributions

Applied to genetic mutations, CrossValQA produces **MutQA**. This paper contributes the following:

- **CrossValQA**, a framework for generating citation-grounded QA pairs from unstructured scientific literature, applicable across molecular-biology domains (genetic mutations here; proteomics as a proof of concept).
- An **information-isolation and bidirectional cross-validation** architecture that controls hallucination and parametric leakage during generated-data construction, with a characterization of its failure modes as a contribution to evaluation methodology.
- **MutQA**: 330,387 QA pairs spanning 6,620 genes and 125,593 gene–variant pairs from 27,442 PubMed Central articles, with sentence-level citations, cross-validation tier labels, and population/cohort metadata for items whose answers depend on a study cohort.
- **Human evaluation** of a stratified 999-pair audit by two expert annotators, reporting raw inter-annotator and human–LLM agreement per validation tier: human–LLM agreement matches human–human agreement on cross-validated QA pairs (87.0% vs. 86.4%) but not on QA pairs that fail cross-validation.
- **Five evaluative use cases (UC1–UC5)** spanning sequence-conditioned generation and question answering, inverse text-to-mutation recovery, and structure-conditioned settings, with homology-aware splits at ≤40% sequence identity and an explicit statement of which claims each supports.
- **Downstream evidence** from two settings: fine-tuning MutaPLM [Luo et al., 2024] on MutQA improves LLM-judge- and human-scored answer quality and inverse mutation recovery, and using MutQA only as a retrieval corpus improves the answer accuracy of four LLM families without any training (4).

## 2 Dataset Scope and Positioning

Existing biomedical QA benchmarks evaluate important capabilities, but they do not jointly measure mutation specificity, free-form mutation-effect generation, and sentence-level citation grounding at scale. BioASQ-QA [Krithara et al., 2023] measures retrieval and answer extraction over expert-curated biomedical questions; PubMedQA [Jin et al., 2019] measure yes/no/maybe reasoning over PubMed abstracts; and MedQA [Jin et al., 2021] measure medical exam-style reasoning. MutaDescribe [Luo et al., 2024] applies a single template to mutation-effect descriptions and ClinVar [Landrum et al., 2018] issues structured significance codes. These benchmarks support biomedical information access and clinical knowledge assessment, but they do not test whether a model can connect a specific mutation to a natural-language claim traceable to cited biomedical evidence.

MutQA targets this missing evaluative setting; its primary life-cycle role is testing and auditing of mutation-aware text models, with secondary roles as fine-tuning supervision and as retrieval context (4). It evaluates whether mutation-aware models can produce, condition on, or recover mutation-specific textual claims grounded in primary literature.

### Supported and unsupported uses

Each MutQA item records what one cited article reports about one variant. MutQA therefore supports provenance-aware training and retrieval, and evaluation of whether a model can recover a variant-specific finding from its cited evidence. It does not establish biological truth, literature consensus, clinical validity, or whether a variant is pathogenic or clinically actionable: a high benchmark score means that a model recovered what the cited paper reported, not that the report is correct or safe for clinical use. Independent articles sometimes disagree about the same variant; MutQA preserves these disagreements rather than resolving them (3), since resolution requires evidence assessment and expert review beyond this dataset. Users who need external evidence strength can join the 88,156 released QA pairs (26.7%) that carry a ClinVar variant name to ClinVar review status.

### 2.1 Evaluative Use Cases and Tasks

MutQA defines five evaluative use cases that vary along two axes: the form of mutation context provided (sequence vs. 3D structure) and whether the model receives an explicit question (Table 1). The reported empirical claims in this paper concern UC1–UC3 only; UC4–UC5 are released as benchmark settings because, to our knowledge, no existing method implements this joint interface for mutations, likely because training data of this kind has been unavailable. UC1 tests whether a model can generate a mutation-effect description from sequence context and mutation identity. UC2 tests whether an explicit natural-language question yields a targeted mutation-specific answer rather than a generic effect description. UC3 tests inverse reasoning from a desired textual effect back to a candidate residue substitution. UC4 and UC5 extend the two forward-generation settings to three-dimensional structural context, where the local residue environment may be relevant for explaining mutation effects (Appendix D.1).

**Table 1:**
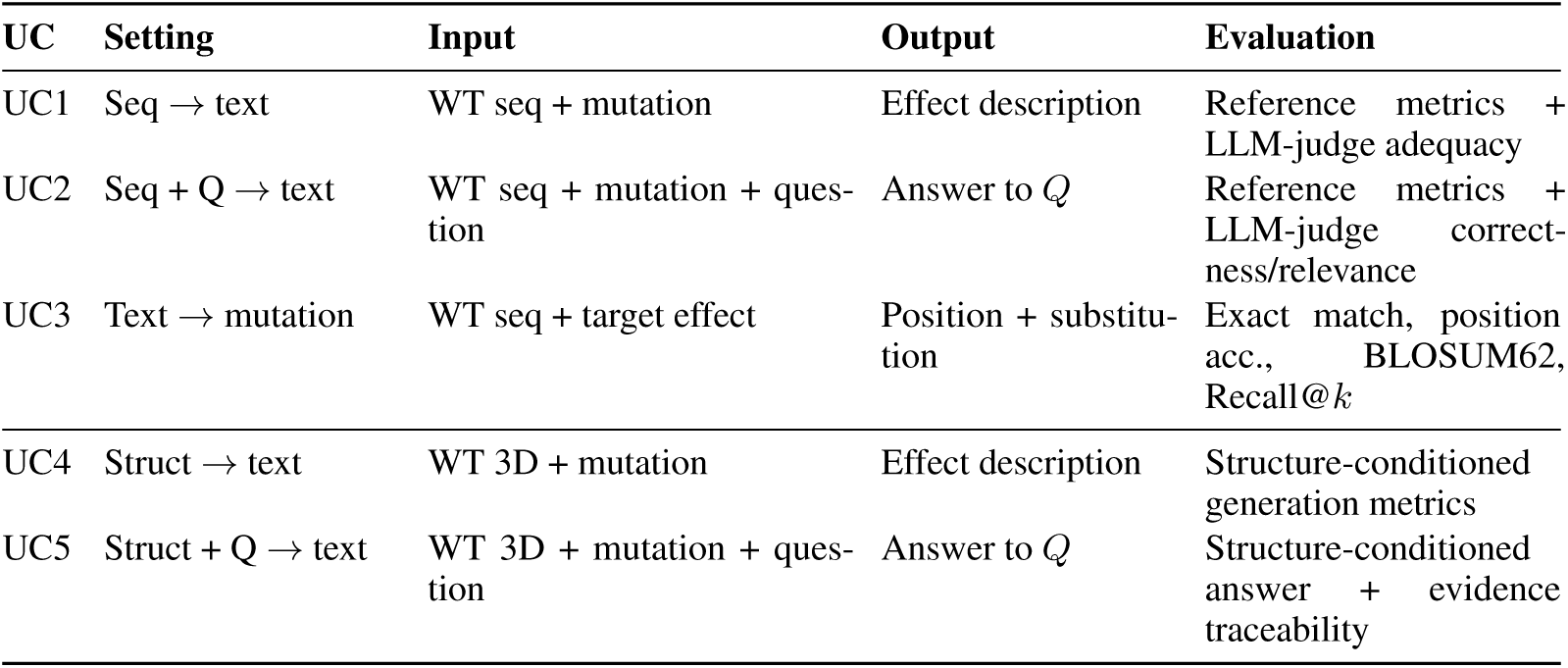
Evaluative use cases in MutQA. UC1–UC3 are used for the sequence-conditioned experiments reported here; UC4–UC5 define released structure-conditioned settings without baseline claims in this paper.

**Table 2:** Feature comparison of MutQA with mutation and biomedical QA resources. Sentence-level citations indicate whether each answer is traceable to a specific span of the source document—a reliability property we argue is essential for LLM-generated supervision. Self-contained indicates that questions embed all required context (gene, variant, aspect) and require no accompanying passage at inference. ClinVar is included as a reference point: it is a curated variant database rather than a QA resource.

| Resource | Size | Source | Mutation-specific | Variant coverage | Free-form | Self-contained | Sentence citations | Structure subset |
| --- | --- | --- | --- | --- | --- | --- | --- | --- |
| ClinVar | ~4.4M | Expert curation | ✓ | All | ✗ | ✓ | ✗<sup>†</sup> | ✗ |
| PubMedQA | 273K | PubMed abstracts | ✗ | None | ✓ | ✗ | ✗ | ✗ |
| BioASQ | 5.5K | PubMed abstracts | ✗ | None | ✓ | ✗ | ✗ | ✗ |
| MutaDescribe | 171K | UniProt + GPT | ✓ | SNV only | ✗ | ✓ | ✗ | ✗ |
| **MutQA** | **330K** | **PMC full text** | ✓ | **All** | ✓ | ✓ | ✓ | ✓ |

<sup>†</sup>ClinVar links to supporting publications but not to specific sentence-level evidence within those publications.

### 2.2 Homology-aware Train/Test Split

To prevent trivial generalization from sequence similarity between train and test, we construct a homology-aware 80/20 split over the cross-grounded release. Records are grouped by UniProt accession so all QA pairs for a given protein stay on one side. Canonical wild-type sequences are clustered with MMseqs2 (–min-seq-id 0.4, -c 0.8, –cov-mode 0), and whole clusters are assigned greedily to splits to hit an 80/20 record-count target (seed 42). To close the residual leakage gap left by representative-based clustering, we apply a GraphPart-style audit [Teufel et al., 2023]: every test protein is searched against the train set with mmseqs easy-search (-s 7.5), and any test UniProt sharing ≥40% identity over ≥80% bidirectional coverage with any train protein is discarded test-side. The final test set has 15 UniProts (1,651 QA pairs) removed and a maximum cross-split identity of 39.7%. The split therefore guarantees no shared UniProt accession across train and test and no test protein within the 40%/80% homology threshold of any train protein; remote homology below this threshold and shared genes or source articles are not controlled. Manifest, parameters, and code are released alongside the dataset for full reproducibility.

#### Out of scope

MutQA does not define primary evaluations for broader protein function prediction, clinical decision support, drug response, quantitative stability or fitness regression, *de novo* design beyond single-residue substitutions, or long-range literature synthesis. These require labels, validation protocols, or aggregation across studies beyond the article-level evidence extracted here. Sentence-level citation links are released as an auditability property rather than a separate use case; downstream attribution evaluations are licensed by the released provenance metadata but require their own task definition.

#### Recommended evaluation methodology

UC1 and UC2: report a three-judge LLM-as-Judge panel as the primary measure, with blinded human scoring of a subset where feasible. Reference-overlap metrics (CIDEr, BLEU, METEOR, ROUGE-L) are secondary, because they reward matching a reference model’s phrasing rather than factual content (Figure 1); the judge prompt, score definitions, and per-judge results are given in Appendix E.2. UC3: exact match on the (position, substitution) pair, position accuracy, BLOSUM62 similarity, and Recall@*k*, which are insensitive to writing style. For attribution-oriented evaluations, evidence-retrieval metrics should be reported separately from answer-quality metrics, since a fluent answer that cannot be traced to its cited sentence does not satisfy MutQA’s reliability criterion.

**Figure 1:**
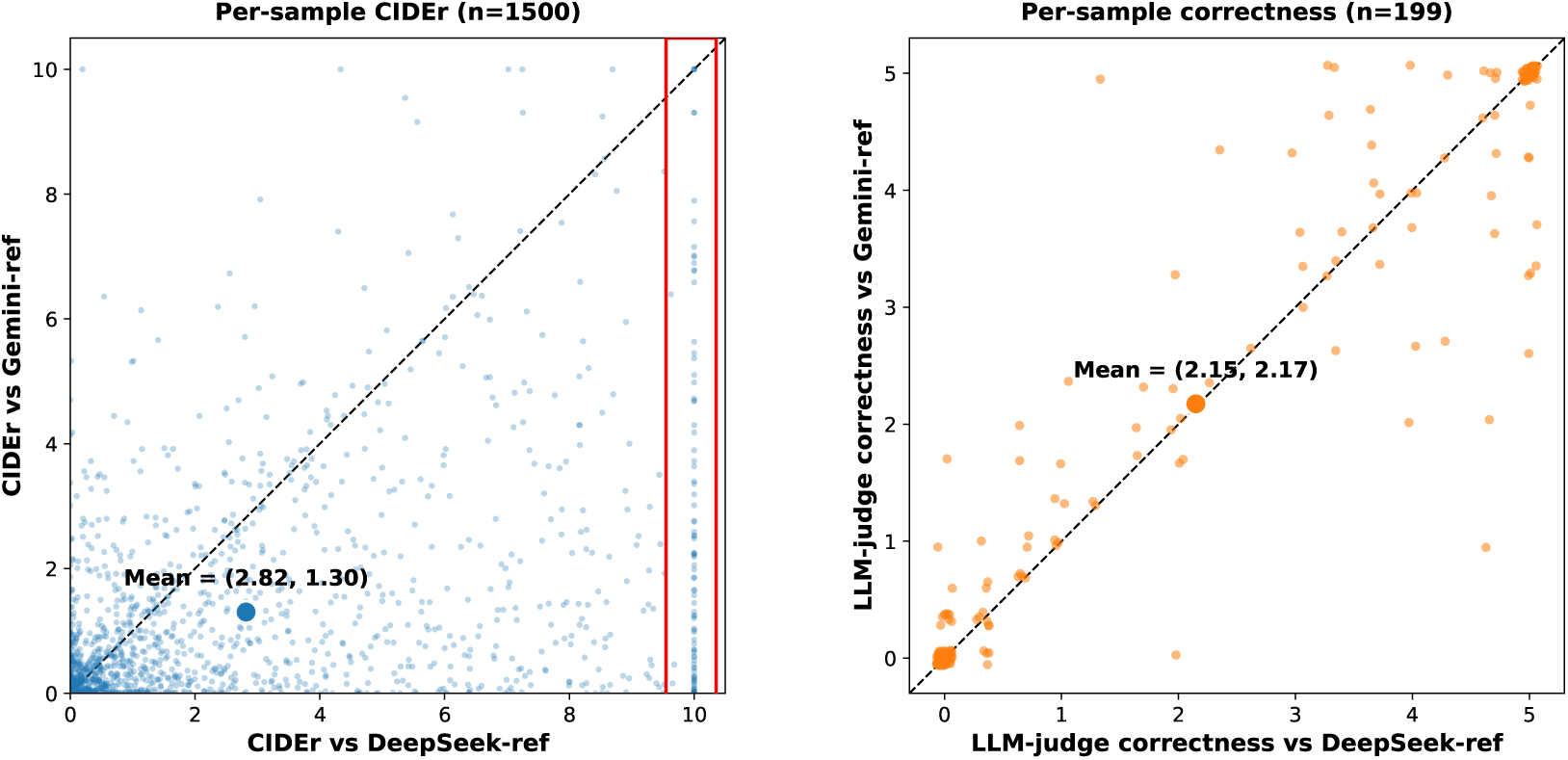
MutaPLM fine-tuned on DeepSeek answers, scored against DeepSeek (x-axis) and Gemini (y-axis) reference answers. CIDEr (left) shows strong below-diagonal asymmetry (mean: 2.82 vs 1.30) — n-gram overlap is inflated by stylistic mimicry of the training-source LLM. The red box highlights 100 samples that scored near-perfect CIDEr against DeepSeek yet span the full 0–10 range against Gemini. LLM-judge correctness (right), averaged over three independent judges, is symmetric (mean: 2.15 vs 2.17) — factual content transfers equivalently across reference LLMs.

#### Comparison to the closest biomedical QA and mutation resources

MutQA differs from prior resources along the 4 axes that govern the downstream use of the benchmark relevant to this paper: the scale of QA pairs, the resource’s primary evaluation targets, whether answer grounding is directly measured (sentence-level evidence *and* an explicit per-pair grounding test), and whether questions are mutation-specific (Table 2). The closest comparators — MutaDescribe [Luo et al., 2024] and ClinVar [Landrum et al., 2018] — provide mutation-level supervision but not sentence-grounded multi-question-type QA: MutaDescribe applies a single template across all entries, and ClinVar issues structured significance codes rather than natural-language answers. The closest comparators on the QA axis — PubMedQA, BioASQ-QA, MedQA — provide free-form biomedical QA but not mutation-specific instances at scale. MutQA combines mutation identity, question-conditioned free-form answers, sentence-level cited evidence, and released benchmark interfaces for both sequence- and structure-conditioned mutation reasoning.

## 3 The MutQA Dataset

Here we present and characterize **MutQA**, a dataset generated by CrossValQA comprising 330,387 citation-grounded QA pairs from 27,442 PubMed Central articles, spanning 6,620 genes and 125,593 gene–variant pairs. Function, pathogenicity, and disease-association questions together make up nearly two-thirds of all QA pairs. Of the 401,406 QA pairs that survive generation, filtering, and the deterministic audit, 82.7% (331,890) pass both directions of cross-validation; the 330,387 of these with a recoverable reference protein sequence form the released benchmark. Unless stated otherwise, statistics below are computed on this benchmark. Additional statistics are in C.2.

### Corpus construction

Articles in the main run enter the corpus through ClinVar’s variant-citation table (var_citations.txt), which links individual variants to PubMed and PMC records, so each is cited by ClinVar for at least one variant. A later gene-coverage expansion, run through the same pipeline, added articles for genes absent from the main run, found through ClinVar citations, PMC searches, and UniProt variant evidence; it contributes 705 articles and 3,665 QA pairs (1.1%) to the benchmark. All text is open-access PMC full text (114,831 articles in the main run); articles without PMC full text are excluded, and no publisher HTML, PDFs, or OCR are used. Reference lists, acknowledgements, and supplementary-reference sections are stripped, and the remaining text is split into sentences by a deterministic rule-based splitter that protects biomedical abbreviations (e.g., *et al*., *Fig.*, decimals). Its indices define the sentence IDs cited by every answer, so grounding can be reproduced from the released article text and code. The full construction funnel is in Appendix C.1: 114,831 PMC articles yield 495,886 generated QA pairs; 411,060 survive the answerability and source filters, 401,406 the deterministic audit, and 82.7% of these (331,890) are cross-validated, of which 330,387 have a recoverable reference sequence and form the benchmark. These are successive filtering stages rather than alternative dataset sizes.

### Coverage and skew

MutQA inherits the ascertainment of the literature it is built from. Per-gene coverage is heavy-tailed (median 4 vs. mean 19 variants per gene; maximum 1,994 for *BRCA1*): the top 10% of genes hold 64% of gene–variant pairs, while 55% of genes carry five or fewer. On the 6,339 genes it shares with ClinVar, MutQA is more concentrated than ClinVar (Gini 0.75 vs. 0.64), and per-gene counts in the two resources are correlated (Spearman *ρ* = 0.59), consistent with research effort concentrating on disease-associated genes. By mutation class, MutQA comprises substitutions 73.8% (ClinVar: 73.0%), small insertions/deletions 13.5% (6.1%), nonsense 6.3% (3.9%), and splice/intronic variants 3.8% (16.3%); ClinVar’s larger splice/intronic share likely reflects intronic variants reported by clinical sequencing panels without functional characterization. MutQA is therefore skewed toward well-studied variants, and aggregate scores should be accompanied by per-class results (4).

### Population context

Allele-frequency, prevalence, carrier-rate, and founder claims are interpretable only relative to a study cohort. Every record carries a population_conditioned flag, set for 9.5% of the benchmark (31,392 QA pairs). For flagged QA pairs, two independent LLM extractors quote the source population phrase under a no-inference rule (ancestry is never inferred from a gene, disease, or location), and a deterministic gazetteer maps it to gnomAD v4 groups (source_population_group), alongside cohort_type and cohort_size. The extractors agree on 89.0% of flagged QA pairs, and disagreements are labelled *contested* (11.0%). For 55.1% of flagged QA pairs no population is reported in the retained evidence, 10.5% describe a global or multi-ancestry population, and 23.4% name specific groups. Among QA pairs assigned a single gnomAD group, European (non-Finnish) and East Asian cohorts account for 66.1% against 4.2% African, directly quantifying the ancestry bias of the source literature. These fields make population-specific items identifiable and filterable; they do not remove the bias.

### Human Evaluation

A stratified audit of 999 QA pairs was scored independently by two molecular-biology annotators and two LLM judges on the same 0–2 correctness rubric (C.3.1). The audit over-represents the lower validation tiers relative to the dataset (one-way validated 11.1% vs. 3.4%; non-validated 25.0% vs. 13.9%) so that agreement can be estimated in each tier. Table 3 reports agreement between individual raters, before any consensus aggregation (Appendix C.3.2 explains why consensus-based agreement overstates reliability on the cross-validated tier). On cross-validated QA pairs, the two annotators agree exactly on 86.4% of items, and individual LLM judges agree with individual annotators on 87.0%: the judges are as close to the annotators as the annotators are to each other. Both figures are inflated by prevalence, since about 90% of annotator ratings on this tier are “correct”; chance-corrected *κ* values and per-tier confusion matrices are in C.3.2. Outside the cross-validated tier, human–LLM agreement falls to 18.2% on one-way validated QA pairs (human–human: 68.5%) and to 50.4% on non-validated QA pairs (human–human: 75.2%). One-way validated QA pairs, which have only partial evidence support, are the hardest to assess, and the annotators agree least often on them. Agreement with the LLM judges falls further because the judges are systematically stricter here, rating only 2% of these answers fully correct against 66–74% for the annotators. The benchmark excludes this tier, which is released with its labels for analysis. We therefore claim human–LLM concordance only for cross-validated QA pairs, and do not claim that the LLM judges reliably rank QA pairs in the lower tiers.

**Table 3:** Raw agreement on the 0–2 correctness rubric by cross-validation tier, before consensus aggregation. Human–human: the two annotators. Human–LLM: mean over the four annotator×judge pairings. LLM–LLM: the two judges. Exact: identical scores (95% bootstrap CI, 10,000 resamples).±1: the two scores are not at opposite ends of the scale (i.e., not 0 vs. 2).

| Tier | *n* | Human–human |  | Human–LLM |  | LLM–LLM |  |
| --- | --- | --- | --- | --- | --- | --- | --- |
| | | Exact (%) | $\pm 1$ (%) | Exact (%) | $\pm 1$ (%) | Exact (%) | $\pm 1$ (%) |
| Cross-validated | 638 | 86.4 (83.7–89.0) | 98.9 | 87.0 (85.0–89.0) | 99.3 | 94.4 (92.5–96.1) | 99.7 |
| One-way validated | 111 | 68.5 (60.4–76.6) | 95.5 | 18.2 (12.4–24.3) | 61.5 | 74.8 (66.7–82.9) | 100.0 |
| Non-validated | 250 | 75.2 (69.6–80.4) | 93.2 | 50.4 (45.8–55.1) | 95.2 | 64.4 (58.4–70.4) | 100.0 |
| All | 999 | 81.6 (79.1–83.9) | 97.1 | 70.2 (67.9–72.5) | 94.1 | 84.7 (82.4–86.9) | 99.8 |

### Deterministic Sequence-Level Audit

As a final, LLM-free QA-pair assessment, we run a two-sided deterministic verifier on the 411,060-pair pre-audit working set (all validation tiers; Appendix C.1): a **protein-coding side** grades p.-prefixed descriptors (178,749 QA pairs) against the UniProt reference sequence, and a **DNA-coding side** grades c.- and bare-nucleotide descriptors (232,311 QA pairs) against the GRCh38 genome and the matched canonical transcript. Seventeen regex-driven claim families — spanning residue and allele identity, position, mutation kind, region class, per-residue physicochemistry, codon-level translation, premature-stop consequence, splice-junction overlap, and UniProt-feature overlap — jointly remove 9,654 QA pairs (2.35%; 7,354 on the protein side and a further 2,300 on the DNA side) whose Phase-1 or Phase-2 answer contradicts the parsed mutation. Claim-family enumeration, trigger discovery, the cleaning funnel, per-claim verification rates, and a case-study panel are in Appendix F.

### Failure Taxonomy

Rejected QA pairs most often fail because independent models make substantively different claims, not because of superficial formatting errors (C.6). Factual disagreement and contradiction dominate the ungrounded tier, showing that the filter catches semantic failures while entity mismatches remain rare (C.6.1). Mechanism and frequency questions are hardest to cross-validate, while pathogenicity and inheritance questions tend to be more directly supported by article text (C.6.2).

### Lexical and Semantic Diversity

MutQA produces broad lexical and semantic variation compared with MutaDescribe’s [Luo et al., 2024] single question template (55k questions). MutQA (23k questions) contains vastly more word types (194k vs. 10) and trigrams (1.3m vs. 8) (C.7.1). The most common question stem, defined as the first five words of a question, accounts for only 5.1% of questions, confirming that the dataset is not dominated by a single prompt pattern (C.7.2). Embedding-based similarity shows MutQA questions occupy a broad semantic space, while answers remain focused on literature-grounded mutation effects (C.7.3).

### Positioning against ClinVar

ClinVar holds about 4.4M variants, but a grounded QA pair requires a study that characterizes the specific variant, and most ClinVar variants cite only generic resources (the ACMG/AMP guideline, PMID 25741868, alone is cited by about 606,000 variants). Only 3.0% of ClinVar variants (129,806) cite a publication shared with at most ten other variants, i.e., a focused study (5.2%, 229,422, at most 100); this literature-characterized subset, not the full catalogue, is the appropriate comparison for MutQA. Neither MutQA’s 125,593 gene–variant pairs nor its 88,587 distinct mutation strings counts unique variants: the same variant is often written in several notations (e.g., c.5095C>T and p.R1699W), and a string such as c.1A>G names a different variant in each of the 58 genes in which it appears. We therefore compare distinct genomic variants. Of the 59,146 distinct variants whose GRCh38 coordinates could be resolved and checked, 40,896 (69.1%) are in ClinVar and 18,250 (30.9%; 26.1% of single-nucleotide variants) are not, consistent with a literature sampling frame in which variants are described in primary studies before, or without, ClinVar submission. Across all matching routes MutQA covers 40,351 ClinVar variants; 64.7% of them belong to the focused-study subset, and they make up 20.1% of that subset. These variants are enriched for well-reviewed and pathogenic classifications (49.1% with two or more review stars vs. 15.2% in ClinVar overall; 58.5% vs. 7.7% pathogenic or likely pathogenic), yet 23,527 (58.3%) are VUS or have at most one review star, the strata in which curated interpretation is most limited (C.8).

### Literature Disagreement

Because every record keeps its source PMID, agreement across articles is measurable. In the benchmark, 14,667 of 125,593 gene–variant pairs (11.7%) are described by two or more articles, covering 94,531 QA pairs (28.6%) and up to 116 articles for a single variant. Comparing answers to the same question type for the same variant across every pair of distinct articles yields 73,411 article pairs; a predefined antonym-polarity check (e.g., pathogenic/benign, increase/decrease, gain/loss of function) flags 5.2% as explicit contradictions. Contradictions concentrate in uncategorized (*other*, 9.5%), pathogenicity (7.0%), frequency (6.7%), mechanism (5.8%), and disease-association (5.2%) questions, against 1.1–1.2% for general function and inheritance. Some flagged article pairs likely reflect differing cohorts, assays, or experimental conditions rather than genuine disagreement, and a lexical check misses paraphrased conflicts, so 5.2% is an estimate. MutQA preserves these disagreements rather than collapsing them into an unsupported consensus (Appendix C.9).

## 4 Application

Table 4 evaluates MutaPLM on the three sequence-conditioned use cases supported by MutQA. Following Section 2.1, blinded human scores and the three-judge LLM panel are the primary measures for UC1 and UC2, and reference-overlap metrics are secondary. Fine-tuning improves every primary measure. In UC2, where the model is also conditioned on a natural-language question, the mean judge score nearly doubles (1.45 → 2.82 on a 0–5 scale; correctness alone 1.33 → 2.49) and the mean human correctness score rises from 0.11 to 0.62 on a 0–2 scale (Wilcoxon *p* = 2.4 × 10*^−^*^6^). In UC1, the open-ended sequence-plus-mutation effect description setting, the judge score rises from 0.96 to 1.62 (correctness 0.74 → 1.31) and the human score from 0.12 to 0.36 (*p* = 8.2 × 10*^−^*^4^). Each judge reproduces the gain on its own (UC1 +0.56 to +0.75; UC2 +0.95 to +1.70; Appendix E.2), although their absolute calibration differs. Human scores track the judges’ correctness scores (Spearman’s *ρ* = 0.52 for UC1 and *ρ* = 0.60 for UC2), and annotators preferred the fine-tuned answer far more often than the pretrained one (UC1 23% vs. 8% of ratings; UC2 44% vs. 6%). Human scores come from four annotators (Appendix E.3); on a 10-item block scored by all four, mean pairwise exact agreement on fine-tuned outputs is 72% for UC2 and 63% for UC1, where it does not exceed chance (Appendix E.3), so the UC1 human result is the weaker of the two. The larger UC2 gains suggest that MutQA’s question-conditioned format gives a more specific and learnable target than a generic mutation-effect prompt. Overlap metrics rise in the same direction but by far larger margins (UC2 ROUGE-L 16.82 → 59.01), which partly reflects learning the reference models’ phrasing. Absolute quality remains low: annotators rated only 10% (UC1) and 21% (UC2) of fine-tuned answers fully correct, against 0% and 3% for the pretrained model. MutQA fine-tuning thus lifts MutaPLM from a near-zero baseline but does not make its answers reliable.

**Table 4:** Effect of fine-tuning MutaPLM on MutQA across UC1 (sequence + mutation → effect description), UC2 (sequence + mutation + question → answer), and UC3 (sequence + target effect description → predicted mutation). Primary measures are listed above secondary reference-overlap metrics. Pre = pretrained MutaPLM; FT = MutQA fine-tune. *^∗^*Mean blinded human correctness (0–2) over 100 ratings per use case by four annotators (70 unique items; Appendix E.3). Mean pairwise exact agreement on a shared 10-item block of fine-tuned outputs: 63% (UC1), 72% (UC2). *^†^*Three LLM judges (DeepSeek-v3.2, Gemini-2.0-Flash, GPT-4o-mini; temperature 0) on a 200-item stratified subset; “judge mean” averages correctness, completeness and relevance (each 0–5) and then the judges, “correctness” is that dimension alone (Appendix E.2). Lexical metrics are percentages (×100).

| Metric | UC1 (N=1,248) |  |  | UC2 (N=1,470) |  |  | UC3 (N=6,119) |  |  |
| --- | --- | --- | --- | --- | --- | --- | --- | --- | --- |
| | Pre | FT | $\Delta$ | Pre | FT | $\Delta$ | Pre | FT | $\Delta$ |
| *Primary* |  |  |  |  |  |  |  |  |  |
| Human mean* (0–2) | 0.12 | **0.36** | +0.24 | 0.11 | **0.62** | +0.51 | — | — | — |
| Judge mean† (0–5) | 0.96 | **1.62** | +0.66 | 1.45 | **2.82** | +1.37 | — | — | — |
| Judge correctness† | 0.74 | **1.31** | +0.57 | 1.33 | **2.49** | +1.16 | — | — | — |
| Accuracy (%) | — | — | — | — | — | — | 19.35 | **32.08** | +12.73 |
| Recall@50 (%) | — | — | — | — | — | — | 2.78 | **10.23** | +7.45 |
| *Secondary (reference overlap)* |  |  |  |  |  |  |  |  |  |
| BLEU-2 | 6.99 | **12.70** | +5.71 | 8.34 | **42.40** | +34.06 | — | — | — |
| BLEU-4 | 1.77 | **8.71** | +6.94 | 3.00 | **33.35** | +30.35 | — | — | — |
| METEOR | 12.91 | **22.30** | +9.39 | 17.26 | **55.09** | +37.83 | — | — | — |
| ROUGE-L | 12.79 | **22.99** | +10.20 | 16.82 | **59.01** | +42.19 | — | — | — |

UC3 evaluates the inverse direction: recovering the mutation associated with a desired textual effect description, scored by exact match, which writing style cannot influence. Fine-tuning improves exact mutation recovery from 19.35% to 32.08% (+12.73 percentage points) and raises Recall@50 from 2.78% to 10.23% (+7.45 percentage points; 3.7× the pretrained value). MutaPLM accepts only amino-acid substitutions and stop-gains, so stratification by mutation class is limited to these two classes; on UC2, fine-tuned performance is similar for missense and nonsense variants (ROUGE-L 59.1 vs. 57.9; judge mean 2.82 vs. 2.81, with only 11 nonsense items judged). Together, UC1– UC3 show that MutQA supervision improves both forward mutation-to-text generation and inverse text-to-mutation prediction under sequence-conditioned inputs.

### Why overlap metrics are secondary

Because MutQA contains validated answers from multiple LLM sources, we can test whether a metric captures factual content or rewards overlap with one reference model’s style. Figure 1 scores a MutaPLM model fine-tuned on DeepSeek answers against DeepSeek and Gemini references for the same questions. CIDEr is strongly reference-sensitive: its mean falls from 2.82 against DeepSeek references to 1.30 against Gemini references (*n* = 1,500), and many predictions with near-perfect CIDEr against DeepSeek score far lower against Gemini, although both references express the same content. LLM-judge correctness is nearly unchanged (2.15 vs. 2.17; Δ = −0.025 on a 0–5 scale; *n* = 199), and no single judge moves by more than 0.03. Reference-overlap metrics therefore partly measure agreement with the training-source model’s phrasing, which is why we report the judge panel and human scores as primary measures and overlap metrics as secondary.

### Training-free evaluation: MutQA as a retrieval corpus

The MutaPLM results draw training and test data from the same source. To test MutQA’s value without any training, we use it only as retrieval context for four LLMs from different developers: GPT-5.6-Terra, Gemini-3.6-Flash, Grok-4.5 and DeepSeek-V4-Pro. For each of 500 test questions, one QA pair per variant is held out as the target. Each model answers from the gene, variant and question alone (closed book) or with up to 20 other MutQA QA pairs about the same variant added to the prompt (+MutQA); context pairs whose question or answer has token-Jaccard overlap ≥ 0.85 with the target are removed. Decoding is identical across arms (temperature 0), and a Claude Haiku 4.5 judge that does not see the arm grades each answer against the held-out reference on the 0–2 rubric of C.3.1. Strict accuracy is the fraction graded 2, compared within model by paired McNemar tests.

Retrieval improves strict accuracy for every model by 20.0 to 28.6 points (Table 5), with lenient (score ≥ 1) gains of 20.8 to 37.0 points, and cuts refusals from 599 to 82 of 2,000 answers. The gains hold on the 403 cross-validated targets alone (+20.8 to +30.8 points, all *p <* 10*^−^*^11^). Three controls point to a variant-specific effect rather than copying. Tightening the question-overlap filter to 0.60 leaves the gains nearly unchanged (+19.4 to +28.0). Also removing context whose answers resemble the reference (token-Jaccard ≥ 0.40) shrinks but does not remove them (+4.7 to +13.4; the GPT-5.6-Terra gain is no longer significant). For GPT-5.6-Terra, replacing the variant’s own QA pairs with an equal number about other variants of the same gene gives no significant gain (+4.0, *p* = 0.21, *n* = 250), so generic gene context or MutQA’s writing style does not reproduce the benefit. Gains are similar across mutation classes except stop-gains (substitutions +25.4, indels +16.5, splice/intronic +26.2, nonsense +3.1 points pooled over models; nonsense covers only 16 unique questions). A blinded human check of 100 items (25 per model) by four annotators reproduces the effect: strict accuracy rises from 29% to 52% (+23 points, McNemar *p* = 1.2 × 10*^−^*^4^), and human and judge grades agree on 81.2% of strict decisions. On a 20-item block scored by all four annotators, mean pairwise exact agreement is 61–65% (Appendix E.4).

**Table 5:** Training-free retrieval evaluation. Strict accuracy (% of 500 answers graded fully correct by a Claude Haiku 4.5 judge) without (closed book) and with MutQA QA pairs about the same variant in the prompt. *p*: paired McNemar test. Controls, per-class results and the human check are in Appendix E.4.

| Model | Closed book | +MutQA | $\Delta$ (pp) | *p* |
| --- | --- | --- | --- | --- |
| GPT-5.6-Terra | 43.8 | 63.8 | +20.0 | $2.3 \times 10^{-14}$ |
| Gemini-3.6-Flash | 34.4 | 59.0 | +24.6 | $4.3 \times 10^{-20}$ |
| Grok-4.5 | 29.4 | 58.0 | +28.6 | $1.6 \times 10^{-23}$ |
| DeepSeek-V4-Pro | 32.2 | 54.0 | +21.8 | $5.7 \times 10^{-16}$ |

The scope of this evaluation is narrow. Retrieval is an oracle lookup by exact variant identity, not a learned retriever, and only variants with more than five QA pairs are eligible (8.3% of variants in the retrieval corpus), so the result does not establish benefit for the long tail of sparsely described variants.

## 5 The CrossValQA Pipeline

CrossValQA (Figure 2) is designed to separate parametric memorization from citation-traceable retrieval in the QA pairs that the evaluation framework of 2 consumes. A first LLM generates a question and an answer–rationale pair (*Q, A*_1_*, R*_1_) from a sentence-cited context window; a second LLM, given *Q* but no access to *A*_1_ or *R*_1_, independently produces (*A*_2_*, R*_2_) from an expanded context window; a judge model then swaps the rationales and asks whether *R*_2_ supports *A*_1_ and whether *R*_1_ supports *A*_2_ — each LLM’s rationale must license the *other* LLM’s answer. Pairs are sorted into three tiers — *cross-validated* (both checks pass), *one-way validated* (only one check passes), and *non-validated* (neither checks pass); all three are released with tier labels preserved (Appendix I).

**Figure 2:**
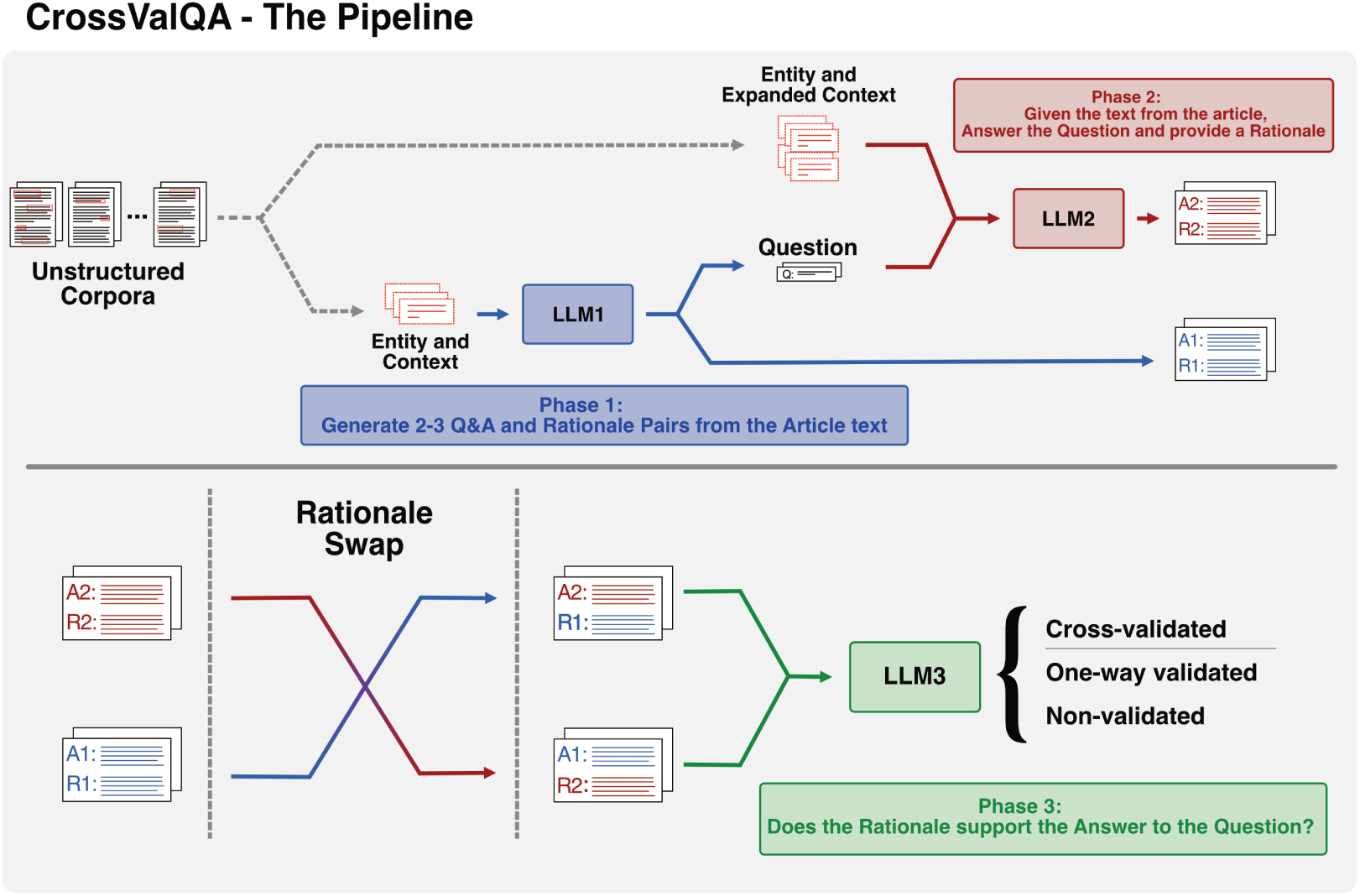
Overview of the CrossValQA pipeline applied to genetic mutations. **Phase 1:** LLM_1_ generates 2–3 question–answer pairs (*Q, A*_1_) with rationales *R*_1_ from the article text. **Phase 2:** Given *Q* but no access to *A*_1_ or *R*_1_, an independent LLM_2_ re-answers from an expanded context window, producing *A*_2_ and *R*_2_. **Phase 3:** A judge LLM_3_ swaps the rationales and evaluates cross-pair support — whether *R*_2_ supports *A*_1_ and whether *R*_1_ supports *A*_2_; only pairs where both cross-pairings pass are labeled *cross-validated*.

CrossValQA is applicable across molecular-biology domains: retargeting it to a new topic within the field requires editing prompts and the entity-resolution module rather than changing the pipeline itself. As a proof of concept we apply it to proteomics (Appendix G). Proteomics shares vocabulary and source resources with genetics, so this pilot does not show that the pipeline transfers to distant scientific fields.

This design supports the central evaluative use of MutQA: testing whether model answers about mutations remain traceable to cited biomedical evidence rather than merely fluent or plausible. The claim is valid when the cited article is treated as the reference source and when the judge is used as a proxy for support between answer and rationale, not for biological truth beyond that source. It is also limited to the cross-validated tier: there, LLM judges agree with expert annotators as often as the annotators agree with each other, but not on one-way or non-validated QA pairs (Table 3). Those tiers are released for analysis of the pipeline, without a claim that the tier labels rank QA quality.

CrossValQA adds two practical safeguards before release. Candidate mutations are verified and evidence spans preserve sentence indices, so each retained answer can be audited back to source text. A final deterministic sequence-level audit removes QA pairs whose sequence-derivable claims contradict the parsed mutation, such as incorrect residue identity, directionally wrong charge changes, or impossible motif claims. Full trigger tables, per-claim verification rates, and cleaning-funnel statistics are reported in Appendix F, and an extended description of the pipeline in Appendix A. All six LLM prompts used to build and evaluate MutQA (QA generation, independent answering, cross-validation judging, QA-audit judging, the UC1/UC2 evaluation judge and the retrieval judge) are released verbatim with their decoding settings in the code repository (prompts/README.md).

## 6 Ablation

The cross-validation judge in our pipeline evaluates each QA pair through two mechanisms acting in concert: the cross-pairing of rationales with opposing answers, and the presence of rationale text itself as evidence for the judge to assess. To understand the relative contribution of each mechanism to the pipeline’s filtering behavior, we conduct two targeted ablation studies. Each ablation removes exactly one component while holding the rest of the pipeline constant, allowing us to attribute changes in grounding tier assignments to the specific mechanism removed. We evaluate both ablations on two independent stratified samples of 3,000 QA pairs (1,000 per grounding tier) and report tier transition matrices showing how items are reclassified when each component is absent.

### Cross-validation Ablation

Here we replace cross-validation with standard validation, i.e., each question is judged against its own rationale rather than the opposing rationale (R1→A1, R2→A2). This causes 72.3% of cross-validation rejects to be accepted, indicating that standard validation is largely redundant: the rationales almost always support their own answers, as shown in Figure 3a (C.4.1).

**Figure 3:**
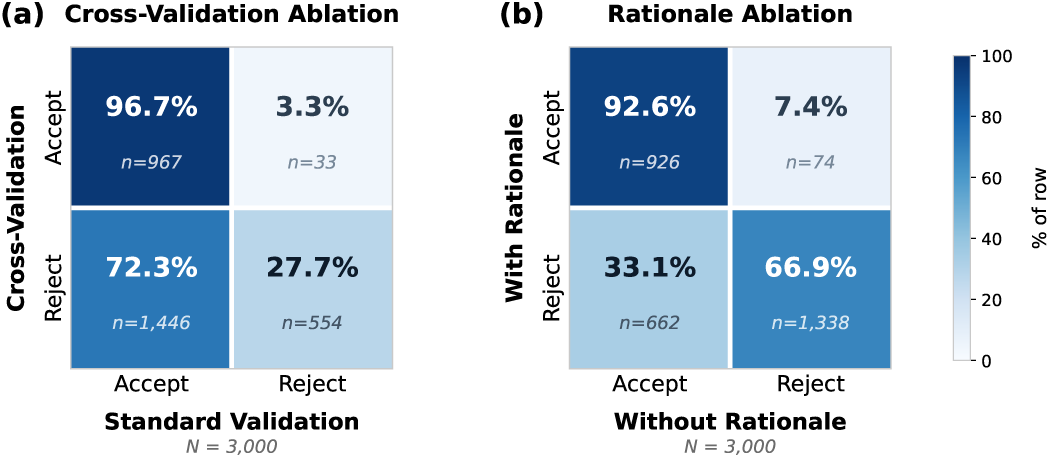
Transition matrices for two ablation studies on the cross-validated QA pipeline. Cells show the row-normalized percentage of items that transition between accept (cross-validated) and reject (one-way validated and non-validated) classifications when a pipeline component is removed.

### Rationale Ablation

Here we remove rationales and instead ask the judge whether one answer supports the claims in the other (A2→A1, A1→A2), assessing direct answer agreement without access to the underlying evidence (C.4.2). Here, 33.1% of items rejected by the full pipeline are accepted without rationales, while 66.9% remain rejected. This indicates that answer-level disagreement alone captures a substantial portion of the filtering signal, but rationales provide meaningful additional discrimination — roughly a third of rejections depend on the judge identifying that the evidence does not support the cross-paired conclusion, even when the answers themselves appear superficially consistent. Furthermore, we show (C.5) that cited rationales are load-bearing evidence traces, not decorative explanations. Rationales are roughly twice as long as answers, cite evidence explicitly, and quote or paraphrase source text rather than merely restating the answer.

## 7 Limitations

### Corpus and publication bias

MutQA inherits the biases of published biomedical literature and is skewed toward well-studied variants. Well-studied genes (e.g., BRCA1, BRCA2, CFTR) are heavily overrepresented: the top 20 genes account for 14.4% of QA pairs (Table 11), and per-gene coverage is more concentrated than ClinVar’s on shared genes (Gini 0.75 vs. 0.64; 3). Beyond gene-level coverage, the corpus reflects positive-result bias, pathogenicity bias (reported variants skew more pathogenic than curated databases would suggest), and disease-gene bias. Ancestry bias is measurable in population-conditioned items: among those assigned a single gnomAD group, European (non-Finnish) and East Asian cohorts account for 66.1% and African cohorts for 4.2%. The population_conditioned flag and cohort fields make these items filterable but do not remove the bias. Non-coding regions and variants in underserved populations are underrepresented. Models should not be assumed to generalize to rare variants or under-curated gene families without recalibration.

### Single-reference grounding

Each QA pair is grounded in one source article, so UC1–UC5 measure agreement with the cited article, not literature consensus. Where several articles describe the same variant, 5.2% of comparable article pairs contain an explicit contradiction (3), concentrated in pathogenicity, frequency and disease-association questions; this bounds achievable cross-article agreement. Released provenance metadata enables multi-reference and contradiction-aware extensions, and the 26.7% of QA pairs that carry a ClinVar variant name can be weighted by ClinVar review status, but neither is the default protocol. Critically, sentence-level citation grounding is not equivalent to biological truth, clinical validity, or literature consensus—benchmark performance measures literature-grounded mutation understanding, not clinical correctness.

### Structure-conditioned use cases lack baselines

UC4 and UC5 require models that jointly consume protein structure, a mutation specifier, and natural language. No released model satisfies all three inputs: structure-aware PLMs (ESM-IF, SaProt, ProstT5) lack mutation conditioning, while mutation-to-text models (MutaPLM, ProtT3) lack structure. The structure-evaluable subset (140,642 QA pairs, 42.6% of the benchmark, over 4,246 proteins; Appendix D.1) is released to enable this model class; until then, UC4–UC5 function as open benchmark calls.

### LLM generation and judging

The dataset is built and partly evaluated by LLMs, which introduces parametric-memory leakage and shared-bias risks. Bidirectional cross-validation (5) and the use of models from different developers (Gemini, DeepSeek and Llama for construction; Gemini, Claude, DeepSeek and GPT-4o-mini as judges) mitigate but do not eliminate these. Human validation is limited in scope. In the 999-pair audit, individual LLM judges agree exactly with individual annotators on 87.0% of cross-validated QA pairs, matching inter-annotator agreement (86.4%), but about 90% of ratings on this tier are “correct”, and human–LLM agreement falls to 18.2% and 50.4% on the one-way and non-validated tiers. The downstream human evaluations use four annotators with small shared blocks (10 items per use case for UC1/UC2, 20 for retrieval), on which mean pairwise exact agreement is 61–72%, and UC1 agreement does not exceed chance. Judge scores should therefore be triangulated with human scoring where possible, with reference-overlap metrics treated as secondary.

### Evaluation scope

MutaPLM is the only sequence-conditioned model we fine-tune, because we know of no other published model that answers open-ended questions about individual mutations; protein sequence-to-text models describe whole-protein function rather than variant effects. We also lack an independent external benchmark: MutaDescribe, the closest resource, was MutaPLM’s training data. The retrieval evaluation removes the train/test coupling and spans four model families, but it uses oracle variant-keyed retrieval rather than a learned retriever and covers only variants with more than five QA pairs (8.3% of variants in its corpus), so it does not establish benefit for sparsely described or unseen variants. Absolute correctness is modest: annotators judged only 10% (UC1) and 21% (UC2) of fine-tuned MutaPLM answers fully correct. Finally, CrossValQA certifies that an answer is reproducible from its source and that LLM judges agree with experts on the cross-validated tier; it does not certify biological ground truth.

### Out-of-scope

MutQA does not evaluate clinical-outcome or drug-response prediction, de-novo design beyond attested single-point variants, non-coding effects, structural variants larger than indels, or non-human variants (the grounded subset is human-filtered).

## 8 Conclusion

MutQA measures citation-grounded mutation reasoning at literature scale: whether a free-form mutation-effect answer remains traceable to specific cited sentences. Its primary life-cycle role is testing and auditing of mutation-aware text models, with secondary roles as fine-tuning supervision and retrieval context. Fine-tuning MutaPLM improves judge- and human-scored answers and inverse mutation recovery, and using MutQA only as retrieval context improves the answer accuracy of four LLM families without training (4). Structure-conditioned settings are released as open benchmark calls for a model class that current sequence-or structure-aware baselines do not yet implement. This evaluation is valid when the cited article is treated as the reference source: it does not establish clinical actionability, cross-paper consensus, or biological truth beyond that source, and human–LLM concordance is established only for the cross-validated tier. CrossValQA, the bidirectional cross-rationale framework that constructed MutQA, is released as a reusable methodology for building auditable QA from primary scientific literature in molecular biology.

### Impact Statement

MutQA is intended for training and evaluating models that reason about genetic variants. Its main risk is misuse in clinical settings: an answer records what one cited article reported, not a validated clinical interpretation, and the dataset over-represents well-studied genes and European and East Asian cohorts. We state supported and unsupported uses in 2, release population and cohort metadata so that population-specific items can be filtered, and advise against using MutQA answers, or model scores on MutQA, for pathogenicity or treatment decisions without expert review. The dataset contains only text from published open-access articles and no information beyond what those articles report.

## A The CrossValQA Pipeline Extended

MutQA generates citation-grounded QA pairs from unstructured scientific literature through a pipeline of targeted entity extraction, evidence-grounded generation, and bidirectional cross-validation (Figure 2). The framework applies across molecular-biology domains; here we instantiate it on genetic mutations from open-access PubMed Central articles.

### A.1 Mutation Extraction and Verification

Candidate mutations are extracted from each article by a hand-curated regex over HGVS-style nomenclature (c., p., m., rs, indels). Because regex extraction is high-recall but low-precision— table headers, gene names with embedded digits, and assay coordinates routinely match—each candidate is independently verified by Gemini 2.5 Flash given a ±3-sentence window around the hit. The verifier classifies the candidate as a real in-text variant or a spurious match; only verified candidates proceed.

### A.2 Context Construction

Articles are filtered to retain high-signal sections (introduction, results, discussion) and discard references, acknowledgments, and procedural boilerplate. Retained sections are segmented into sentences and assigned a canonical sentence index that persists throughout the pipeline. For each mention of a domain-relevant entity co-occurring with mechanistic terminology, a ±2-sentence span is extracted; spans for the same entity discussed in multiple locations are merged into a single context window. Sentence indices are preserved so that all downstream outputs are auditable back to specific source spans.

### A.3 Grounded QA Generation (LLM_1_)

LLM_1_ receives the entity identifier, the consequence summary from LLM_1_, and the same numbered evidence sentences. It generates two to three QA pairs per entity. Each answer must include a rationale that quotes at least one phrase from the evidence, cites the supporting sentence number(s), and explains how the evidence supports the answer. The entity identifier must appear in the question, answer, and rationale. To diversify question framing, the prompt requires at least one negatively framed question per entity. This stage produces the initial answer and rationale sets (*A*_1_, *R*_1_).

### A.4 Independent Answer Generation (LLM_2_)

LLM_2_ (DeepSeek-Chat, distinct model family from LLM_1_) receives the question, the entity identifier, and an expanded context window of ±5 sentences around every entity mention in the source article. Critically, LLM_2_ does not receive *A*_1_, *R*_1_, or the consequence summary, so verification is fully independent. If the question is unanswerable from the expanded context, the instance is flagged. Otherwise, LLM_2_ produces an independent answer and rationale (*A*_2_, *R*_2_) under the same citation constraints.

### A.5 Bidirectional Cross-Validation (LLM_3_)

A judge model (Llama 3.1 8B) evaluates mutual consistency between the two answer–rationale pairs by checking two conditions: (i) whether *R*_2_ provides sufficient evidence to support *A*_1_, and (ii) whether *R*_1_ provides sufficient evidence to support *A*_2_. QA pairs satisfying both are classified as **cross-validated**; pairs satisfying only (i) are **one-way validated**; pairs satisfying neither are **unvalidated**. Requiring two independent reasoning paths—produced by different models from different context windows—to converge on mutually supporting conclusions filters hallucinated inferences, external knowledge leakage, and weakly evidenced claims.

## B Extended Discussion on Related Work

### Instruction Tuning

Large Language Models (LLMs) have demonstrated significant capabilities in human comprehension tasks, such as GPT models [Radford et al., 2019, Brown et al., 2020, OpenAI et al., 2023] and LLaMA models [Touvron et al., 2023a,b, Dubey et al., 2024]. When LLMs were first introduced, they were primarily trained on next-token prediction [Touvron et al., 2023a, Radford et al., 2019, Lewis et al., 2020, Liu et al., 2019, Yang et al., 2020]. Instruction tuning has been proposed to align the training objective with the user objective by enhancing the model’s ability to follow instructions [Zhang et al., 2024]. Several models trained via instruction tuning have been proposed for a variety of tasks such as summarization [Basyal and Sanghvi, 2023], question answering [Ouyang et al., 2022, Muennighoff et al., 2023, Zheng et al., 2023], and zero-shot capabilities [Zheng et al., 2023, Ouyang et al., 2022, OpenAI et al., 2023, Wei et al., 2022, Dubey et al., 2024].

### Multimodal LLMs

Multimodal LLMs have also been extensively applied to perform cross-modal tasks beyond the text modalities. For instance, several studies have been proposed to integrate vision and language [Liu et al., 2023a, Alayrac et al., 2022, Li et al., 2023], and integrate audio and language [Radford et al., 2022, Tjandra et al., 2017]. Building on these efforts, LLMs have also witnessed prosperous adaptation to scientific and biomedical domains such as biomedical text understanding [Jararweh et al., 2024, Lee et al., 2019], biomedical QA [Wu et al., 2023, Luo et al., 2023], clinical reasoning tasks [Huang et al., 2020], and molecular structure understanding [Zhao et al., 2024, Fang et al., 2024, Cao et al., 2023, Liu et al., 2023b].

### Protein-related LLMs

Many QA datasets have been curated to allow for instruction tuning on biomedical tasks. General biomedical QA datasets such as PubMedQA [Jin et al., 2019] and BioASQQA [Krithara et al., 2023] provide a wide corpus of QA examples that span diseases, genes, proteins, drugs, etc. While these datasets are comprehensive, they are limited to creating question-answer pairs from ontologies that do not capture the rich textual context present in biomedical literature. Mol-Instructions provides an extensive QA dataset specific to bio-molecules (including proteins) to extend LLMs to biomolecular structure. The sequential nature of protein primary structure lends itself to language modeling for protein characterization. For example, encoder-based LLMs trained on protein amino acid sequences have been adopted to generate a representation space that captures sequence structures [Lin et al., 2022b,a, Zhang et al., 2022, Elnaggar et al., 2021]. Similarly, generative LLMs have also been proposed for a variety of protein generation tasks, such as generating 3D structure [John Jumper and Hassabis, 2021], and novel protein sequences [Madani et al., 2020, Nijkamp et al., 2022, Lv et al., 2024].

### 3D Structure-based and General Biomedical QA Datasets

Datasets like PDB-QA [Liu et al., 2024] exclusively focus on predefined queries derived from 3D protein structural information. Meanwhile, general biomedical QA datasets, such as PubMedQA [Jin et al., 2019], encompass a broad range of biomedical concepts, including diseases, proteins, and drugs, making them suitable for general biomedical QA tasks but insufficient for sequence-to-text translation tasks.

## C Extended MutQA Characterization

### C.1 Construction Funnel

Table 6 lists every filtering stage from the PMC source pool to the public splits, with the unit counted at each stage.

### C.2 Dataset Statistics

#### Question Type Distribution

The dataset spans eight major question categories, reflecting the diverse aspects of mutation characterization in biomedical literature. Function-related questions dominate the dataset (24.3%), followed by pathogenicity assessment (21.3%), disease association (19.3%), and other specialized categories including frequency (8.5%), clinical features (7.7%), mechanism (3.4%), and inheritance patterns (3.2%). This distribution reflects the natural emphasis of biomedical literature to understand the functional and clinical significance of genetic mutations.

#### Cross-Validation Quality

Of the 401,406 QA pairs that pass the deterministic audit (Appendix C.1), **82.7%** are classified as *cross-validated*, meaning both validation directions passed—the independent rationale supports the original answer and vice versa. A further 3.4% are *one-way validated* (one direction passed) and 13.9% are *non-validated*; both are released with their tier labels but excluded from the benchmark. The high cross-validation rate indicates that most QA pairs have converging independent derivations from the source text.

**Table 6:** MutQA construction funnel. Each row is a successive filter on the row above, in the unit shown.

| Stage | Unit | Count |
| --- | --- | --- |
| *Main run* |  |  |
| PMC open-access articles cited by ClinVar | articles | 114,831 |
| With $\geq 1$ regex hit | articles | 88,615 |
| After corpus filters<sup>a</sup> | articles | 86,317 |
| Mutation candidates sent to LLM verification<sup>b</sup> | candidates / articles | 1,379,377 / 75,321 |
| Verified variants with functional discussion | variants / articles | 234,148 / 33,901 |
| Variant records entering QA generation | records | 167,879 |
| After independent re-answering (Phase 2) | QA pairs | 495,886 |
| Scored by the cross-validation judge (Phase 3) | QA pairs | 473,243 |
| Grounded to a UniProt accession | QA pairs | 416,188 |
| *Merged release* |  |  |
| Gene-coverage expansion<sup>c</sup> | QA pairs | +7,227 |
| Merged dataset | QA pairs | 423,415 |
| After answerability and consistency filters | QA pairs | 411,060 |
| After deterministic sequence-level audit (−9,654) | QA pairs | 401,406 |
| of which cross-validated | QA pairs | 331,890 |
| Cross-validated, recoverable reference sequence (benchmark) | QA pairs / articles | 330,387 / 27,442 |
| After GraphPart homology audit (public splits) | QA pairs | 328,736 |

<sup>a</sup>Removes articles under 5,000 characters, articles with more than 200 regex hits, and one outlier article (42,374 hits). <sup>b</sup>Regex hits remaining after 150,859 gene-symbol matches are removed. <sup>c</sup>The same Phase 0–3 pipeline run on 1,079 articles targeting 1,041 genes absent from the main run (ClinVar-cited articles, PMC searches for genes without ClinVar citations, and UniProt humsavar evidence); contributes 705 articles and 3,665 QA pairs to the benchmark.

**Table 7:** Mutation type distribution in MutQA. Categories are assigned by HGVS-style pattern matching on the mutation field; *indel* (delins) and bare = silent annotations contribute *<* 0.01% each and are folded into *Other*.

| Mutation type | Count | % |
| --- | --- | --- |
| Substitution | 249,025 | 75.37% |
| Deletion | 27,266 | 8.25% |
| Nonsense | 18,711 | 5.66% |
| Duplication | 6,727 | 2.04% |
| Insertion | 3,071 | 0.93% |
| Intronic / splice | 11,967 | 3.62% |
| Frameshift | 6,621 | 2.00% |
| Indel | 4 | 0.00% |
| rs ID | 1,199 | 0.36% |
| Other | 5,796 | 1.75% |
| Total | 330,387 | 100.00% |

#### Mutation Type Coverage

The dataset spans the mutation classes typical of clinical genomics literature. Counted over benchmark QA pairs, substitutions (missense and silent) form the majority (75.4%; 73.8% when counted over gene–variant pairs, as in 3), consistent with their

#### Text Characteristics

Questions average 12.4 words (SD = 3.2), reflecting concise, well-formed queries. Answers from Phase 1 (initial generation) average 18.6 words (SD = 8.8), while Phase 2 (independent verification) answers average 16.0 words (SD = 8.0). The modest difference in answer length between phases suggests that independent derivations converge on similar levels of detail.

**Table 8:** Summary statistics of the MutQA dataset.

| Statistic | Value |
| --- | --- |
| Total QA pairs | 330,387 |
| Unique source articles (PMIDs) | 27,442 |
| Unique genes | 6,620 |
| Unique variants | 88,587 |
| Unique gene–variant pairs | 125,593 |
| Avg. question length (tokens) | 20.7 ± 5.2 |
| Avg. answer length (tokens) | 31.4 ± 13.6 |
| Vocabulary size | 296,744 |
| Avg. citations per QA pair | 4.20 |
| Cross-grounded (%) | 100.0% |
| Grounded (%) | 0.0% |
| Ungrounded (%) | 0.0% |

**Table 9:** Distribution of question types in MutQA.

| Question type | Count | % |
| --- | --- | --- |
| Function | 80,285 | 24.3% |
| Pathogenicity | 70,401 | 21.3% |
| Disease | 63,651 | 19.3% |
| Other | 40,542 | 12.3% |
| Frequency | 28,012 | 8.5% |
| Clinical | 25,434 | 7.7% |
| Mechanism | 11,352 | 3.4% |
| Inheritance | 10,710 | 3.2% |
| Total | 330,387 | 100.0% |

### C.3 Human Evaluation and Fault Taxonomy

To validate the quality of MutQA and assess the feasibility of LLM-based annotation as a scalable proxy for human review, we conducted a systematic agreement study between human expert annotations and LLM judgments on a stratified sample of QA pairs.

#### C.3.1 Annotation Protocol

We sampled 1,000 QA pairs from MutQA, stratified by cross-validation status (cross-grounded, grounded, and ungrounded). Each QA pair was independently scored by two human annotators with domain expertise in molecular biology. Annotators rated each QA pair on a three-point ordinal scale: Correct (2)—the answer is factually accurate and fully supported by the source passage; Partially Correct (1)—the answer contains relevant information but includes minor inaccuracies or omissions; Incorrect (0)—the answer is factually wrong, unsupported, or contradicts the source evidence. In parallel, two LLM judges—Gemini 2.0 Flash (LLM1) and Claude Haiku 3.5 (LLM2)—scored the same QA pairs using the same rubric and source passages. For both human annotators and LLM judges, we computed consensus scores by averaging and rounding to the nearest integer, yielding a Human Consensus and an LLM Consensus score for each QA pair.

#### C.3.2 Agreement Metrics

Table 3 (main text) reports raw agreement per tier. Table 12 adds chance-corrected statistics, computed between individual raters: annotator 1 vs. annotator 2, judge 1 (Gemini 2.0 Flash) vs. judge 2 (Claude Haiku 3.5), and the mean over the four annotator×judge pairings. % Agree: exact matches. *κ*: unweighted Cohen’s kappa. *κ_w_*: quadratically weighted kappa.

**Table 10:** Literature and gene coverage in MutQA.

| Statistic | Value |
| --- | --- |
| Source articles (PMIDs) | 27,442 |
| QA pairs per article (mean) | 12.0 |
| QA pairs per article (median) | 6 |
| Unique genes | 6,620 |
| Genes with $\geq 5$ QA pairs | 4,545 |
| Genes with $\geq 10$ QA pairs | 3,314 |
| Genes with $\geq 50$ QA pairs | 1,296 |
| Genes with $\geq 100$ QA pairs | 691 |
| QA pairs per gene (mean) | 49.9 |
| QA pairs per gene (median) | 10 |
| Unique gene-variant pairs | 125,593 |

**Table 11:** Top 20 genes in MutQA by QA-pair count. Together they account for 14.4% of the dataset.

| Gene | QA pairs | % |
| --- | --- | --- |
| BRCA1 | 6,500 | 1.97% |
| BRCA2 | 5,031 | 1.52% |
| GJB2 | 3,608 | 1.09% |
| LDLR | 3,044 | 0.92% |
| CFTR | 2,884 | 0.87% |
| ABCA4 | 2,523 | 0.76% |
| ATP7B | 2,260 | 0.68% |
| TP53 | 2,077 | 0.63% |
| ATM | 2,071 | 0.63% |
| SLC26A4 | 1,897 | 0.57% |
| USH2A | 1,771 | 0.54% |
| SCN5A | 1,740 | 0.53% |
| MLH1 | 1,694 | 0.51% |
| FBN1 | 1,668 | 0.50% |
| PAH | 1,616 | 0.49% |
| GAA | 1,508 | 0.46% |
| CHEK2 | 1,452 | 0.44% |
| MUTYH | 1,413 | 0.43% |
| LPL | 1,413 | 0.43% |
| VWF | 1,407 | 0.43% |

##### Consensus aggregation inflates agreement

Consensus scores are formed per item as ⌈(*r*_1_+*r*_2_)*/*2⌉ for the two annotators and for the two LLM judges. On the cross-validated tier, where about 90% of individual ratings are 2, this rule converts every 1/2 disagreement into a 2 and so concentrates both consensus vectors at the majority class. Agreement between consensus vectors on this tier is therefore 94.4%, against 87.0% between individual annotators and individual judges and 86.4% between the two annotators; the consensus-based *ρ* = 0.974 and *κ_w_* = 0.953 reported in earlier versions of this analysis reflect the same effect. The main text (Table 3) therefore reports agreement between individual raters only.

##### Why raw agreement and kappa diverge

On the cross-validated tier, 90.3% of annotator ratings and 95.6% of judge ratings are 2. Under this prevalence kappa is deflated (the kappa paradox; Feinstein & Cicchetti, 1990): the annotators agree exactly on 86.4% of items but *κ* = 0.23, and individual human–LLM pairs agree on 85.6–88.4% with *κ* of 0.01–0.03. Raw agreement shows that judges and annotators reach the same label as often as two annotators do; kappa shows that on this tier neither annotators nor judges discriminate reliably among the few items not rated fully correct. We report both and use neither to claim that the judges can rank cross-validated items. Table 13 gives the annotator confusion matrices per tier. The audit over-represents the lower tiers (cross-validated QA pairs are 63.9% of the audit and 82.7% of the dataset), so the pooled “All” row is not a dataset-level estimate.

**Table 12:** Agreement between individual raters on the 0–2 correctness rubric, by cross-validation tier (*n* = 999). Human–LLM: mean over the four annotator×judge pairings.

| Tier | $n$ | Human–human | | | Human–LLM | | | LLM–LLM | | |
| --- | --- | --- | --- | --- | --- | --- | --- | --- | --- | --- |
| | | % | $\kappa$ | $\kappa_w$ | % | $\kappa$ | $\kappa_w$ | % | $\kappa$ | $\kappa_w$ |
| Cross-validated | 638 | 86.4 | 0.23 | 0.20 | 87.0 | 0.02 | 0.03 | 94.4 | 0.33 | 0.30 |
| One-way validated | 111 | 68.5 | 0.29 | 0.22 | 18.2 | 0.03 | −0.01 | 74.8 | 0.52 | 0.58 |
| Non-validated | 250 | 75.2 | 0.57 | 0.46 | 50.4 | 0.08 | 0.08 | 64.4 | 0.29 | 0.35 |
| All | 999 | 81.6 | 0.48 | 0.44 | 70.2 | 0.21 | 0.18 | 84.7 | 0.61 | 0.76 |

**Table 13:** Annotator confusion matrices by tier (rows: annotator 1; columns: annotator 2; scores 0/1/2).

| Cross-validated ($n=638$) | | | | One-way ($n=111$) | | | Non-validated ($n=250$) | | |
| --- | --- | --- | --- | --- | --- | --- | --- | --- | --- |
|  | 0 | 1 | 2 | 0 | 1 | 2 | 0 | 1 | 2 |
| 0 | 0 | 0 | 0 | 1 | 0 | 4 | 4 | 14 | 17 |
| 1 | 1 | 18 | 43 | 0 | 15 | 18 | 2 | 83 | 19 |
| 2 | 7 | 36 | 533 | 1 | 12 | 60 | 0 | 10 | 101 |

#### C.3.3 Agreement Stratified by Grounding Status

To evaluate whether cross-validation status tracks annotation reliability, we stratify agreement by tier (Tables 3 and 12). Earlier versions of this analysis reported Spearman correlations between consensus scores (*ρ* = 0.974, 0.289 and 0.613 for cross-validated, one-way and non-validated QA pairs). We withdraw them: *ρ* measures monotone association rather than agreement, and on a three-level scale with about 90% of scores at 2 it is dominated by the consensus rule described above.

On raw agreement, the cross-validated tier is where annotators agree most with each other (86.4%) and with the judges (87.0%). The one-way validated tier, whose QA pairs have only partial evidence support, is the hardest: annotators agree on 68.5%, and human–LLM agreement falls to 18.2% because the judges rate only 2.3% of these answers fully correct against 66–74% for the annotators. Non-validated QA pairs fall in between (75.2% human–human, 50.4% human–LLM); clearly unsupported answers are easier to agree on than partially supported ones. Chance-corrected human–LLM agreement is low in every tier (*κ* ≤ 0.08). We therefore claim human–LLM concordance only for the cross-validated tier, in the raw-agreement sense above, and do not claim that tier labels or LLM judges rank QA pairs in the lower tiers.

### C.4 Ablation

Transition matrices for two ablation studies on the cross-validated QA pipeline, each evaluated on two independent stratified samples of 3,000 QA pairs (1,000 per grounding tier) are shown in Figure 4. Cells show the row-normalized percentage of items that transition between validation tiers when a pipeline component is removed.

#### C.4.1 Cross-Validation Ablation

The pipeline’s cross-validation judge makes two independent calls per QA pair: one asks whether the Phase 2 rationale supports the Phase 1 answer (R2→A1), and another asks whether the Phase 1 rationale supports the Phase 2 answer (R1→A2). To isolate the effect of this cross-pairing, we replace it with standard validation, where each rationale is judged against its own answer: R1→A1 and R2→A2. The judge prompt, model, and grounding logic are otherwise identical. Figure 4a shows that 96.7% of Tier 1 items remain in Tier 1 under standard validation, confirming that cross-grounded items are robustly validated by either method. However, 78.9% of Tier 3 (ungrounded) items and 65.7% of Tier 2 (grounded) items are promoted to Tier 1 under standard validation, while fewer than 1% of items from any original tier fall into Tier 3. This near-total collapse into Tier 1 confirms that rationales almost always support their own answers, making standard validation largely unable to distinguish quality tiers. Cross-validation is effective precisely because it forces each rationale to support a conclusion it was not generated to defend.

**Figure 4:**
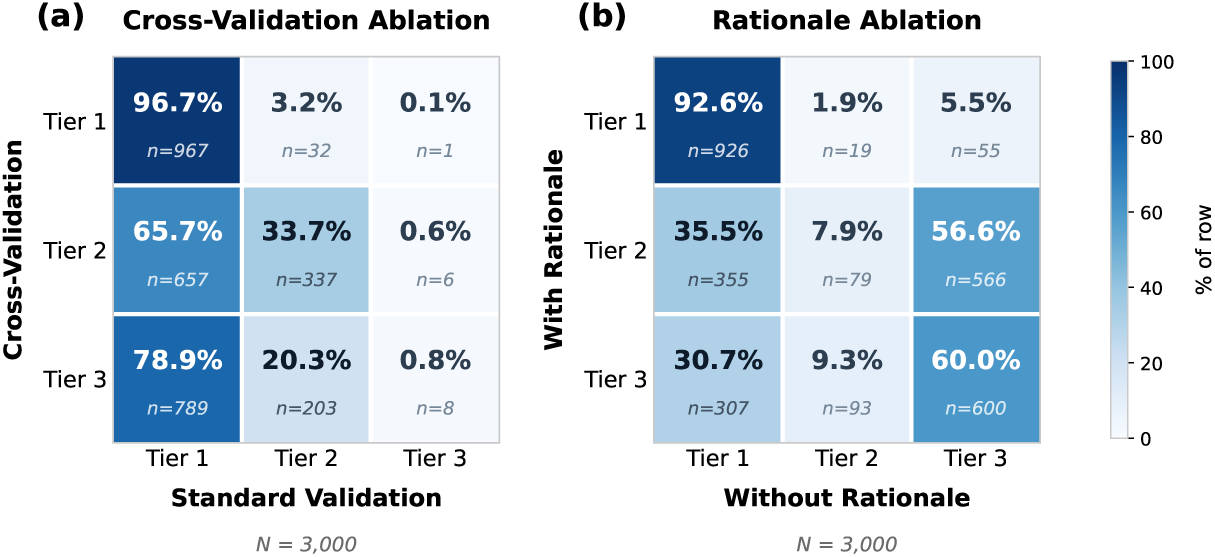
Transition matrices for two pipeline ablations on stratified samples of 3,000 QA pairs (1,000 per tier). Tier 1 = cross-validated, Tier 2 = one-way validated, Tier 3 = non-validated. Cells are row-normalized: rows are the full-pipeline tier, columns are the ablated-pipeline tier. **(a) Cross-Validation Ablation:** swaps rationale-swap judging for same-pair judging. **(b) Rationale Ablation:** removes rationales from the judge prompt while preserving cross-pairing.

#### C.4.2 Rationale Ablation

Here we ablate the rationales. The cross-validation structure is preserved — the judge still evaluates two cross-paired combinations per QA pair — but the rationales are removed from the prompt entirely. Instead of asking whether a rationale supports an answer, the judge is asked whether one answer supports the claims in the other (A2→A1, A1→A2), assessing direct answer agreement without access to the underlying evidence.

The three tiers behave very differently. **Cross-grounded** verdicts are largely preserved (92.6%): when both models produced answers that obviously agree, the judge reaches the same conclusion without rationales. **Ungrounded** verdicts are preserved 60% of the time; critically, 30.7% of them are *falsely promoted to cross-grounded* when rationales are removed—the judge cannot detect the specific evidence gaps that motivated the original rejection without the rationale text to audit. The **grounded** tier collapses almost completely: only 7.9% of grounded verdicts survive, with 56.6% re-labeled ungrounded and 35.5% promoted to cross-grounded. In other words, the partially-supported middle regime is where rationales carry essentially all of the judge’s discriminative signal.

Aggregated over the 3,000-sample set, balanced accuracy between rationale-full and rationale-free verdicts is 53.5% with Cohen’s *κ* = 0.303—fair agreement. The population-weighted accuracy of 85.1% is dominated by the cross-grounded mass (82.6% of the corpus), which masks the instability in the grounded and ungrounded tiers. The practical implication is that the rationales are not decorative: stripping them would shift approximately 1 in 6 pipeline verdicts, inflate false cross-groundings by several percentage points, and effectively eliminate the grounded tier as a meaningful category.

### C.5 Rationale quality

We sampled 500 records per grounding tier (seed = 42) and, for each Phase 1 and Phase 2 rationale, measured: (a) *citation rate*—whether the rationale mentions any sentence ID listed in its `evidence_ids`; (b) *token Jaccard* between rationale and answer, as a restatement proxy; (c) *length ratio* relative to the answer; (d) *quotation rate*, whether the rationale contains a verbatim substring of the source passage; and (e) presence of logical connectives (*because, therefore, this suggests, . . .*). Table 14 reports these metrics by tier.

Several patterns are consistent across tiers. Rationales are **roughly twice the length of their answers** (2.0–2.3×) and cite at least one of the evidence sentence IDs in 70%–94% of records, with Phase 2 rationales citing more reliably than Phase 1 (median ∼91% vs. ∼73%). Answer–rationale Jaccard overlap is moderate (∼0.38), indicating that rationales repeat the entity and claim tokens while adding substantial additional text—consistent with evidence citation rather than pure restatement. Phase 1 rationales contain direct source quotations 60%–64% of the time; Phase 2 rationales, which operate from a larger context window, paraphrase more heavily (18%–22% quotation rate) but carry logical connectives less frequently. In short, rationales are longer than answers, cite evidence explicitly, and quote or paraphrase source text—the profile of reasoning traces rather than restatements.

**Figure 5:**
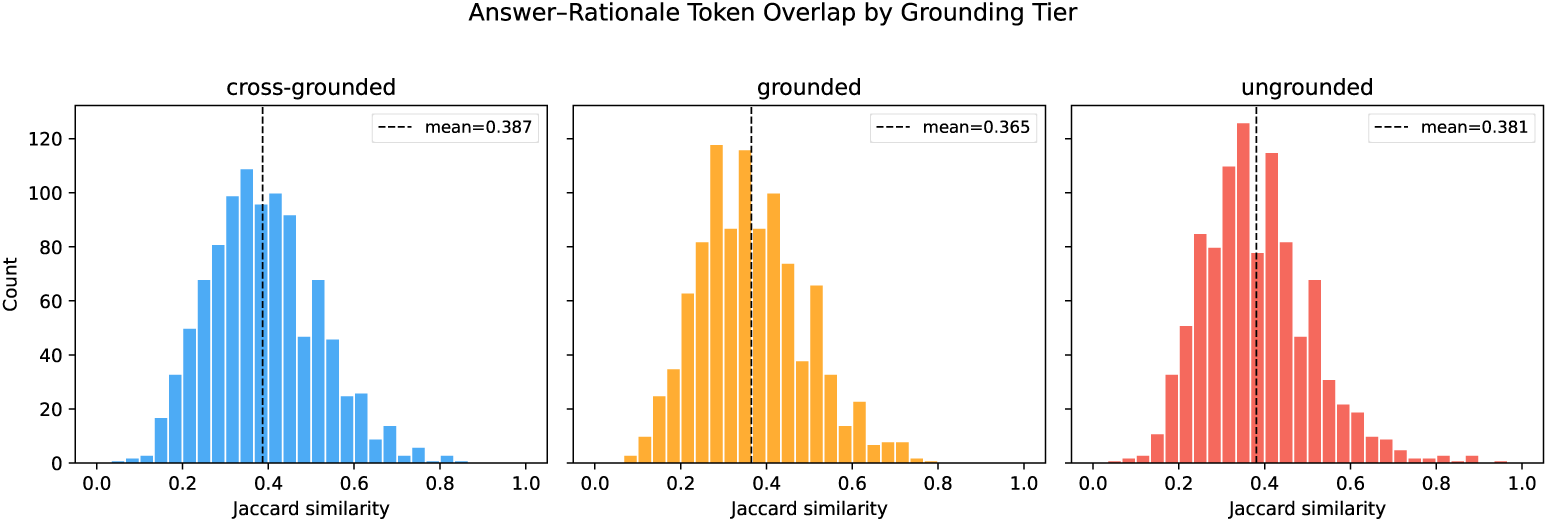
Distribution of token-level Jaccard similarity between Phase-1/2 answers and their rationales, by grounding tier (500 records per tier). The dashed line in each panel marks the per-tier mean.

**Table 14:** Reasoning quality metrics by grounding tier and phase.

| Tier | Phase | Cite% | Jaccard | Len ratio | Conn.% | Quote% | Genuine% | Restate% |
| --- | --- | --- | --- | --- | --- | --- | --- | --- |
| cross-grounded | 1 | 73.6 | 0.386 | 2.32 | 19.0 | 63.6 | 19.0 | 22.0 |
| cross-grounded | 2 | 89.0 | 0.388 | 1.98 | 8.2 | 22.0 | 8.2 | 15.6 |
| grounded | 1 | 76.2 | 0.371 | 2.28 | 34.4 | 63.0 | 34.4 | 16.4 |
| grounded | 2 | 93.6 | 0.359 | 2.17 | 13.2 | 22.4 | 13.2 | 10.0 |
| ungrounded | 1 | 70.2 | 0.381 | 2.35 | 28.2 | 59.8 | 28.2 | 21.8 |
| ungrounded | 2 | 91.8 | 0.380 | 1.97 | 12.4 | 17.6 | 12.4 | 12.6 |

### C.6 Failure Taxonomy: What Do Rejected QA Pairs Look Like?

The Ablation (Section C.4) demonstrates that the pipeline’s exclusion of the *ungrounded* tier is well-founded—raw *A*_1_ and *A*_2_ diverge in ways that correctly disqualify these pairs from the high-confidence subset. However, “ungrounded” is a coarse label, and from an evaluation standpoint it matters *why* a pair fails: contradictions between two LLMs reading the same article are a different signal than evidence gaps or scope mismatches. A model that will ultimately consume this dataset as supervision or as an evaluation target benefits from knowing which failure modes the filter catches, and which it is silent about.

We sampled 2,000 ungrounded records uniformly at random (seed = 42) and applied nine regex-defined categories to the concatenation of the two judge rationales (*R*_2_ → *A*_1_ and *R*_1_ → *A*_2_). Categories are not mutually exclusive—a single failure can simultaneously exhibit, e.g., contradiction and specificity mismatch—so percentages sum to more than 100%. Table 15 reports the distribution; the mean record carries 1.69 category labels.

#### C.6.1 Interpretation

*Factual disagreement* (51.6%) and *contradiction* (48.2%) dominate—together, these capture cases in which the two LLMs arrived at substantively different claims from the same article. This is the failure mode the pipeline is explicitly designed to catch, and its dominance is evidence that the cross-validation stage is operating on semantic content, not on surface-level format mismatches. *Evidence gap* (24.6%) and *specificity mismatch* (24.9%) together represent cases where one answer is adequately supported and the other is under-specified or over-reaches beyond what the cited sentences establish; these are the failure mode that motivates sentence-level citation tracking as a dataset feature rather than an afterthought. *Entity mismatch* (4.3%) captures the rarer cases in which one model answered about a different variant or gene than the question specifies—a failure the pipeline’s entity-centered context construction is specifically designed to suppress, and its low rate is consistent with that design working. The long tail (*scope disagreement*, *answerable disagreement*, *insufficient evidence*; each *<* 1%) contains edge cases that future iterations of the framework could address with targeted prompt refinements.

**Table 15:** Taxonomy of cross-validation failure patterns (sample of 2,000 ungrounded records). Records may belong to multiple categories.

| Failure Category | Count | % |
| --- | --- | --- |
| Factual disagreement | 1,032 | 51.6% |
| Contradiction | 964 | 48.2% |
| Specificity mismatch | 497 | 24.9% |
| Evidence gap | 491 | 24.6% |
| Other/unclassified | 259 | 12.9% |
| Entity mismatch | 87 | 4.3% |
| Answerable disagreement | 18 | 0.9% |
| Insufficient evidence | 16 | 0.8% |
| Scope disagreement | 14 | 0.7% |
| Avg. categories/record |  | 1.69 |

**Figure 6:**
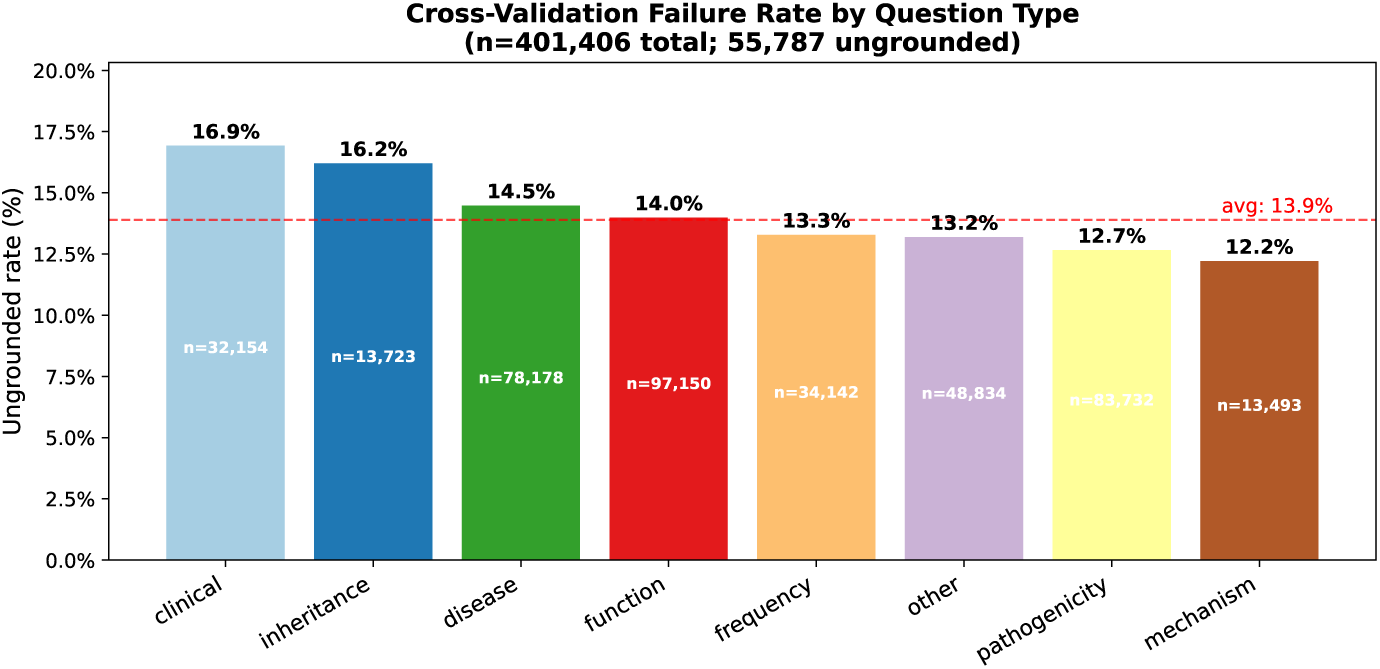
Ungrounded rate by question category, with the corpus-wide average as a dashed line. Mechanistic and frequency questions are hardest to cross-validate; pathogenicity and inheritance are easiest.

#### C.6.2 Ungrounded rate by question type

Figure 6 reports the ungrounded rate per question category. The rate is highest for *mechanism* and *frequency* questions and lowest for *pathogenicity* and *inheritance*, which we read as a reflection of how directly each question type maps to an unambiguous article claim: pathogenicity assignments and inheritance patterns are typically stated explicitly in the text, whereas mechanistic claims require synthesizing multiple sentences and are more likely to produce divergent independent derivations. This variation is *itself* an evaluation signal: models that perform uniformly across question types despite these intrinsic difficulty differences are more likely overfitting to surface patterns than doing grounded reasoning. Per-question-type scoring for UC1–UC3 (Section 2.1) is therefore recommended as a default reporting convention for this benchmark.

### C.7 Lexical and Semantic Diversity

A standing concern with LLM-generated supervision is that bulk scale can hide a narrow underlying distribution: a dataset nominally containing hundreds of thousands of QA pairs may reduce to a few templates repeated over different entities. Because MutQA generates questions with an LLM rather than by slotting entities into fixed patterns, we quantify whether the resulting supervision is genuinely diverse and, where possible, benchmark against a templated baseline from the same domain. Our reference point is **MutaDescribe** [Luo et al., 2024], which uses a single question template (*“Based on the following protein sequence, please describe its function.”*) across all 55,541 of its test entries.

#### C.7.1 Lexical diversity

We compare vocabulary size, unique trigram count, and type–token ratio (TTR) across MutQA questions, answers, and rationales, alongside MutaDescribe’s questions and answers. TTR is known to shrink with corpus size, so for a size-controlled comparison we compute TTR on matched 50,000-token random samples averaged over 10 bootstrap draws (seed = 42); we also report absolute vocabulary and trigram counts, which reflect raw lexical coverage. Results appear in Table 16.

The question-level comparison is the sharpest: MutQA contains **177,376** distinct word types and **1,083,748** unique trigrams across 330,387 questions, versus **10** word types and **8** trigrams across MutaDescribe’s 55,541 questions—the latter being a consequence of its single fixed template. Sampled TTR shows the same gap (0.168 vs. 0.000). Answer-side lexical coverage also favors MutQA in absolute terms (242,210 vs. 67,823 word types; 2.0M vs. 0.69M unique trigrams), even though per-text TTR is comparable (∼ 0.20 for both), indicating that both datasets produce locally varied answers but MutQA spans a substantially larger lexical space overall. Rationales—released alongside the dataset as an auditing resource—contribute a further 660,774 word types and 6.6M unique trigrams.

#### C.7.2 Question stem analysis

To verify that diversity is not concentrated in a single dominant template, we extract the first five tokens of each question as a *stem* and count unique stems. MutQA contains **123,878** unique 5-gram stems across 330,387 questions (stem ratio = 0.38), and the most common stem (*“What is the functional effect”*) accounts for only 5.2% of questions. The top 20 stems together cover just 33% of the corpus, confirming that the dataset is not dominated by a single template. MutaDescribe’s stem ratio, by construction, is 1*/*55,541 ≈ 2 × 10*^−^*^5^.

#### C.7.3 Semantic diversity

Lexical metrics can be satisfied by paraphrases that remain semantically near-identical, so we additionally measure semantic spread via pairwise cosine similarity over sentence-transformer embeddings (all-MiniLM-L6-v2). For each corpus we sample 30,000 texts and compute cosine similarity on 500,000 random pairs. Figure 7 shows the resulting distributions. MutQA questions have mean pairwise cosine similarity of 0.43 (std 0.10), whereas MutaDescribe’s identical-template questions trivially saturate at 1.0. For rationales, MutQA reaches a comparable mean of 0.42.

For *answers*, MutQA has higher mean cosine similarity (0.37) than MutaDescribe (0.25). We read this as a signature of *task focus* rather than lack of diversity: MutaDescribe answers span arbitrary protein functions (localization, catalysis, binding, regulation, . . .), while MutQA answers are constrained to mutation-effect claims grounded in the cited literature. The relevant comparison for dataset utility is not broadest possible answer dispersion, but broadest *question* coverage over a focused answer space—and on that axis (Table 16, question row; Figure 7, left panel) MutQA is substantially more diverse. This matches the underlying construction: question stems are LLM-generated from per-article evidence rather than slotted into a template, so the question distribution inherits the heterogeneity of the source literature while the answer distribution remains anchored to a well-defined task.

### C.8 Coverage Overlap with Existing Mutation Resources

The dataset comparison in Table 2 positions MutQA qualitatively against prior biomedical QA and mutation resources, but it does not quantify what variant coverage the dataset actually adds over existing sources. For a benchmark, this distinction matters: a model’s score on MutQA is informative to the extent that the dataset exercises capabilities not already covered elsewhere. We therefore compute direct variant-level intersections against the two closest comparators: **MutaDescribe** [Luo et al., 2024], the only other mutation-specific QA dataset at scale, and **ClinVar** [Landrum et al., 2018], the authoritative clinical variant database.

**Table 16:** Lexical diversity comparison between MutQA and MutaDescribe. Sampled TTR computed on 50k-token windows (mean ± std over 10 bootstrap samples). MutaDescribe uses a single question template, yielding minimal question diversity.

| Corpus | N | Vocab | Trigrams | TTR (50k) | Mean Len |
| --- | --- | --- | --- | --- | --- |
| MutQA Questions | 330,387 | 177,376 | 1,083,748 | 0.168±0.001 | 12.4 |
| MutaDescribe Question | 55,541 | 10 | 8 | 0.000±0.000 | 10.0 |
| MutQA Answers | 330,387 | 242,210 | 2,047,945 | 0.202±0.002 | 18.6 |
| MutaDescribe Answers | 55,541 | 67,823 | 691,484 | 0.200±0.001 | 22.3 |
| MutQA Rationales | 660,774 | 645,222 | 6,686,597 | 0.205±0.002 | 34.3 |

**Figure 7:**
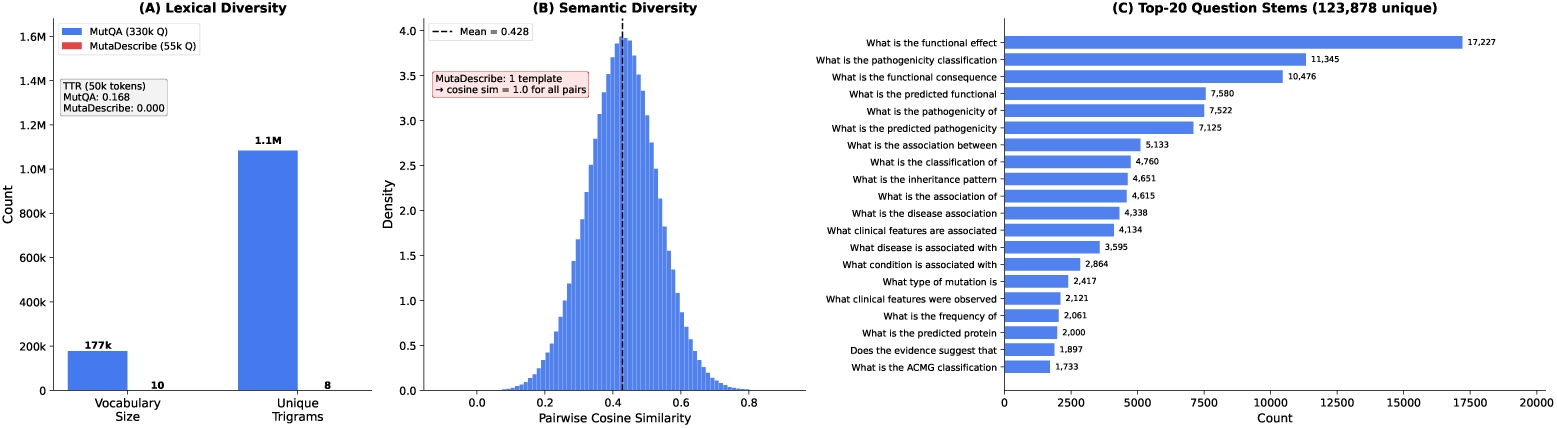
Question diversity in MutQA vs. MutaDescribe. **A)** Vocabulary size and unique whitespace trigrams; inset reports type–token ratio on a 50K-token sample (mean over 10 bootstrap shuffles). **B)** Pairwise cosine-similarity distribution of sentence-transformer (all-MiniLM-L6-v2) embeddings, computed over 500K random pairs from a 30K-question sample. **C)** Top-20 most frequent 5-word question stems.

#### Normalization

To intersect across sources with heterogeneous HGVS conventions, we extract a normalized 1-letter protein-substitution identifier (e.g., F294A) from each source’s variant string, using a shared regex over both 3-letter (p.Phe294Ala) and 1-letter (F294A) forms. The gene name is prepended. This restricts the overlap analysis to single amino-acid substitutions—the variant class all three resources natively represent. Indels, frameshifts, splice-site, intronic, and rsID-indexed variants are not normalized to a shared space and are excluded from this comparison.

- **Same variants, different supervision.** ClinVar records a *clinical significance code* per variant (e.g., “Pathogenic,” “Likely benign”); MutaDescribe provides a single templated sentence per entry. Neither offers sentence-cited, natural-language, multi-question-type supervision of the kind UC1–UC3 (Section 2.1) require. For variants shared with ClinVar, MutQA functions as a *parallel annotation layer* rather than a coverage extension.
- **Literature-grounded tail.** Variants absent from ClinVar are, by construction, discussed in primary literature but not clinically curated. On the substitution-only view above they number 25,113; among the 59,146 distinct variants whose genomic coordinates could be checked (below), 18,250 (30.9%).

#### Genome-coordinate matching

The substitution-only view counts normalized protein identifiers. For the comparison in 3 we instead match distinct genomic variants against ClinVar (February 2026 release; 4,372,439 GRCh38 VariationIDs). MutQA mutation strings are resolved to GRCh38 coordinates with offline HGVS mapping or the Ensembl VEP REST API; 78,147 of the 125,593 benchmark gene–variant pairs resolve to a coordinate-checkable variant, and these correspond to 59,146 distinct genomic variants (the same variant often appears in several notations, e.g., c.5095C>T, 5095C>T and p.R1699W). Single-nucleotide variants are matched on chromosome, position and alleles, and deletions and duplications on chromosome, start, end and variant type; a coordinate match also requires ClinVar to list the same gene. Variants without usable coordinates are matched on gene plus the coding or protein HGVS string parsed from the ClinVar variant name. Of the 59,146 checkable variants, 40,896 (69.1%) are in ClinVar and 18,250 (30.9%) are not; the share is 26.1% for single-nucleotide variants and 53.5% for indels, where differing indel representations likely inflate the figure. For 37,688 gene–variant pairs neither route applies, so their ClinVar status is unknown. Across all routes, MutQA covers 40,351 distinct ClinVar variants.

**Figure 8:**
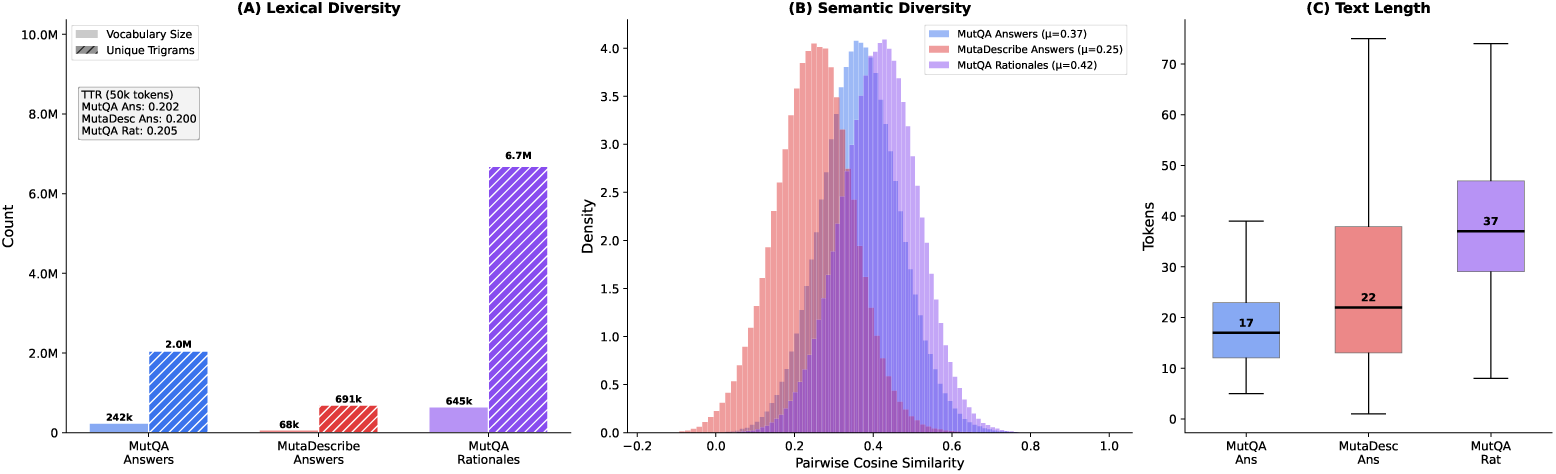
Answer and reasoning-trace diversity for MutQA Answers (Phase 1), MutaDescribe Answers, and MutQA Rationales (Phases 1 + 2 concatenated). **A)** Vocabulary size (solid) and unique whitespace trigrams (hatched); inset reports type–token ratio on a 50K-token sample. **B)** Pairwise cosine-similarity distributions of sentence-transformer (all-MiniLM-L6-v2) embeddings, 500K random pairs from 30K-record samples per corpus. **C)** Distribution of text length in whitespace tokens (50K-record samples; fliers hidden).

**Table 17:** Quantitative overlap between MutQA and existing mutation-specific resources. Variants are gene-anchored: each unit is a (HGNC gene, protein-variant) pair, with the protein variant normalized to 1-letter HGVS form (e.g., BRCA1:F294A). MutaDescribe is keyed by UniProt accession, mapped to HGNC symbols via UniProt’s idmapping files. MutaDescribe is filtered to human-only (47.4%).

| Variant category | Count | % of MutQA |
| --- | --- | --- |
| MutQA total (gene-anchored) | 49,928 | 100.0% |
| In MutaDescribe | 7,568 | 15.2% |
| In ClinVar | 23,860 | 47.8% |
| In both | 6,613 | 13.2% |
| **Unique to MutQA** | **25,113** | **50.3%** |

#### Literature-characterized subset of ClinVar

Most ClinVar variants cite only generic resources (the ACMG/AMP guideline, PMID 25741868, is cited by 605,919 variants), so we count variants that cite at least one PubMed record shared with at most *N* other variants. For *N* = 10 this focused-study subset holds 129,806 variants (3.0% of ClinVar), and for *N* = 100, 229,422 (5.2%). Of MutQA’s 40,351 ClinVar variants, 64.7% belong to the *N* = 10 subset and 79.0% to the *N* = 100 subset, covering 20.1% and 13.9% of those subsets respectively. Against all of ClinVar, MutQA’s variants are enriched for well-reviewed, pathogenic and conflicting classifications (Table 18); the higher share of conflicting classifications is consistent with the cross-article disagreement in Appendix C.9.

gnomAD and GenomicKB are not comparable on coverage: gnomAD’s entries are observed genotypes, overwhelmingly ultra-rare and without phenotype or functional description, and GenomicKB is a knowledge graph of genomic elements rather than a catalogue of characterized variant effects.

### C.9 Multi-Source Consistency and Literature Disagreement

Because MutQA draws from the open biomedical literature rather than from a single curated annotation source, the same gene–variant pair often appears in multiple independent articles. This redundancy creates a measurable property other mutation benchmarks lack: we can ask whether independent articles *agree* about a given variant, and—when they do not—whether the disagreement reflects pipeline extraction error or genuine scientific dispute. Both have direct evaluative consequences: a model trained on MutQA will be exposed to literature-level disagreement, and downstream evaluation must distinguish a model that surfaces such disagreement from one that confabulates a false consensus.

**Table 18:** ClinVar review status and classification of the ClinVar variants covered by MutQA, against all of ClinVar (GRCh38, February 2026). Review stars follow ClinVar’s review-status levels.

| Category | MutQA ∩ ClinVar ($n = 40,351$) | All ClinVar ($n = 4,372,439$) |
| --- | --- | --- |
| Expert panel or guideline (3–4 stars) | 2,963 (7.3%) | 21,471 (0.5%) |
| Two or more review stars | 19,830 (49.1%) | 664,081 (15.2%) |
| Pathogenic / likely pathogenic | 23,600 (58.5%) | 337,308 (7.7%) |
| Benign / likely benign | 2,910 (7.2%) | 1,331,317 (30.4%) |
| Uncertain significance (VUS) | 6,893 (17.1%) | 2,285,661 (52.3%) |
| Conflicting classifications | 6,263 (15.5%) | 161,758 (3.7%) |
| VUS or at most one review star | 23,527 (58.3%) | 4,000,395 (91.5%) |

#### Method

We group benchmark QA pairs by gene–variant pair and keep the 14,667 pairs (11.7% of 125,593) described by at least two distinct articles; they cover 94,531 QA pairs (28.6%), with up to 116 articles for a single variant. Within each (gene, variant, question type) group we take one answer per article and compare every pair of distinct articles, giving 73,411 article pairs. Each pair is checked for explicit contradictions with six biomedical antonym families (*yes*/*no*, *pathogenic*/*benign*, *associated*/*not associated*, *increase*/*decrease*, *gain-of-function*/*loss-of-function*, *activating*/*inactivating*). A pair is flagged only if one answer contains the positive term *and lacks* the negative term, and the other answer the converse, which avoids spurious matches on hedged or compound statements. This exhaustive analysis replaces the 1,000-group sample reported in earlier versions of this paper (4.5% contradictions).

#### Results

Of the 73,411 article pairs, 3,805 (5.2%) contain an explicit contradiction on at least one antonym family (Table 19). On the full 401,406-pair release the rate is the same (5.2% of 96,570 article pairs).

**Table 19:** Explicit contradictions between answers from distinct articles about the same variant and question type (benchmark). Five pairs labelled *functional* are included in the total but not listed.

| Question type | Article pairs | % of pairs | Contradictions (%) |
| --- | --- | --- | --- |
| Disease association | 19,235 | 26.2 | 5.2 |
| Pathogenicity | 16,161 | 22.0 | 7.0 |
| Function | 15,069 | 20.5 | 1.2 |
| Frequency | 12,841 | 17.5 | 6.7 |
| Other | 4,850 | 6.6 | 9.5 |
| Clinical | 3,878 | 5.3 | 2.8 |
| Mechanism | 921 | 1.3 | 5.8 |
| Inheritance | 451 | 0.6 | 1.1 |
| All | 73,411 | 100.0 | 5.2 |

#### Disagreement is concentrated in semantically loaded question types

Pathogenicity (7.0%), frequency (6.7%), mechanism (5.8%) and disease-association (5.2%) answers contradict each other several times as often as general function (1.2%) or inheritance (1.1%) answers, and uncategorized questions have the highest rate (9.5%). Descriptive function annotations tend to converge across studies, while clinical interpretations, frequencies and disease associations remain contested. Some flagged pairs likely reflect different cohorts, assays or experimental conditions rather than genuine disagreement, and the lexical check misses paraphrased conflicts, so 5.2% is an estimate that can err in either direction.

#### Contradictions reflect literature dispute, not pipeline error

Every QA pair in the benchmark is cross-validated: each was independently judged supported in both directions of the bidirectional cross-validation (Section A.5). All 3,805 contradictions therefore occur between answers that each passed the strictest pipeline filter on their own evidence. This matches the design intent of cross-validation: the step verifies that an answer is supported by its cited evidence, not that it agrees with what *other* articles say about the same variant. Persistent multi-source disagreement is therefore a property of the underlying literature surfaced by the dataset, not noise to be filtered away.

**Figure 9:**
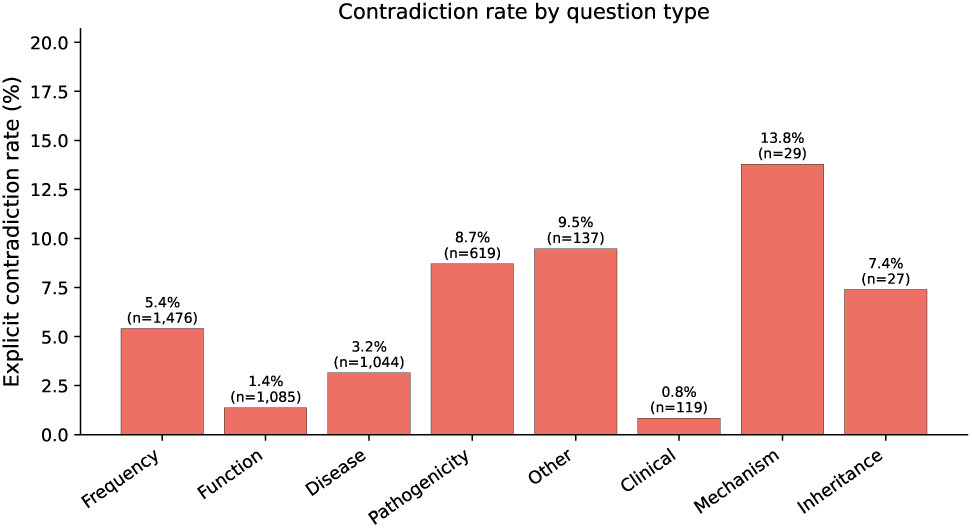
Explicit-contradiction rate by question type across 73,411 pairs of distinct articles describing the same variant and question type. Pathogenicity, frequency and disease-association questions disagree most often; general function and inheritance questions rarely do.

## D Extended Dataset Statistics

This section provides additional visualizations of the MutQA dataset composition, complementing the summary statistics presented in Section 3.

**Figure 10:**
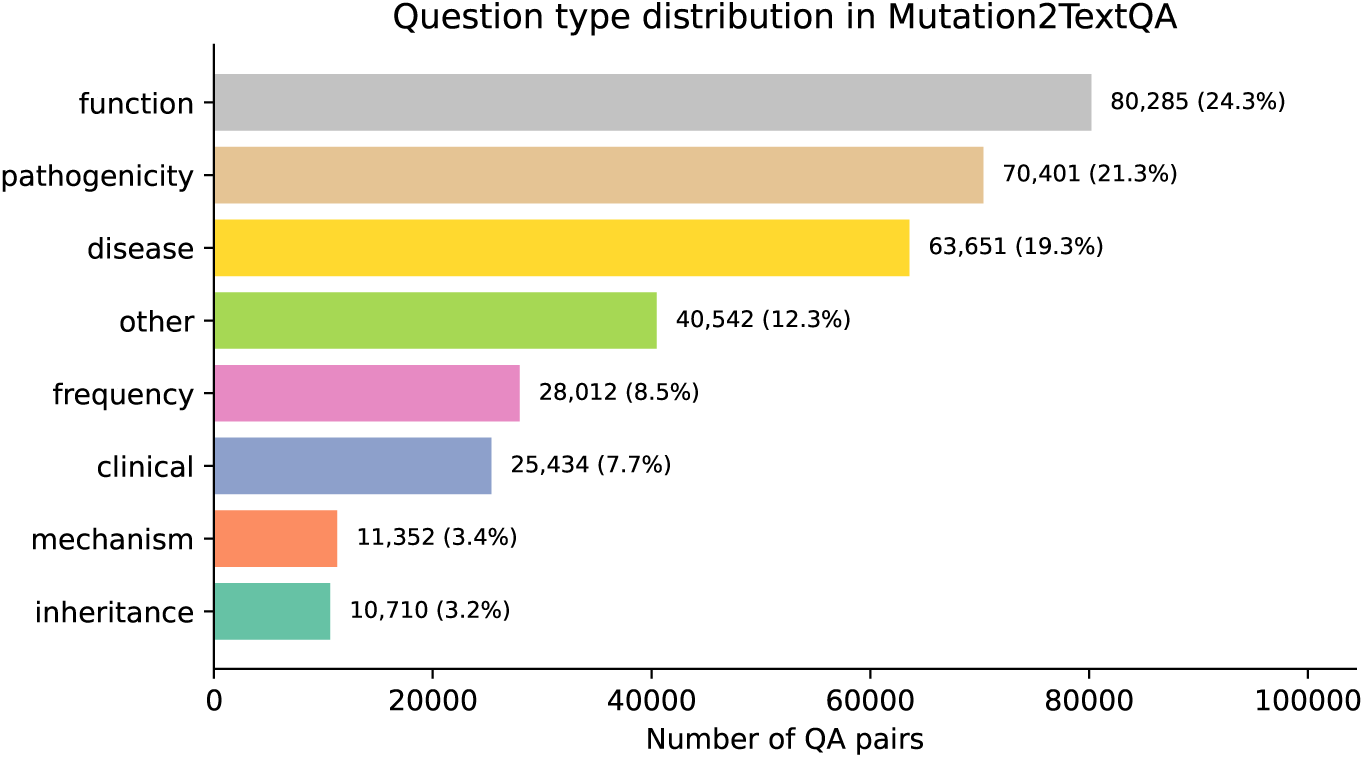
Distribution of question types in MutQA. Function-related questions (24.3%) dominate the dataset, followed by pathogenicity (21.3%), disease association (19.3%), and other specialized categories. This distribution reflects the natural emphasis in biomedical literature on understanding the functional and clinical significance of genetic mutations.

### D.1 UC4/UC5 Structure-Annotated Subset

To support downstream analyses that require 3D-structural context (e.g., evaluating whether a model’s reasoning aligns with the structural environment of a residue), every benchmark QA pair carries structure fields: its UniProt accession’s experimentally determined PDB entries (pdb_ids), its AlphaFold v4 model (alphafold_id), the mutated residue’s protein position where it can be parsed (`mutation_position`), and the UniProt sequence features that overlap that position (`mutation_locations`). A QA pair is *structure-evaluable* (`structure_evaluable`) when it has both a structure and a residue position; these 140,642 QA pairs (42.6% of the benchmark) form the UC4/UC5 subset.

**Figure 11:**
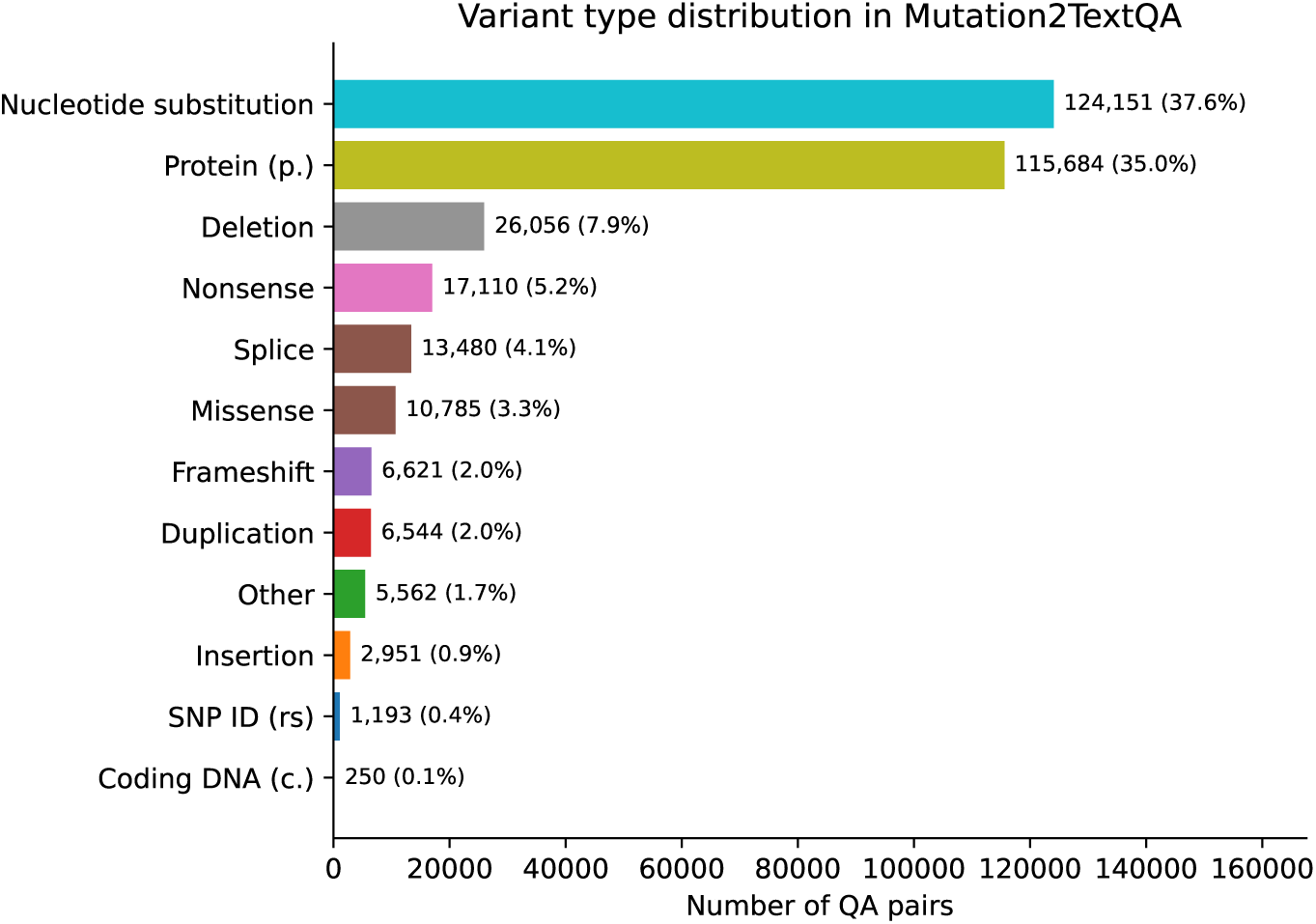
Distribution of mutation types in MutQA.

**Figure 12:**
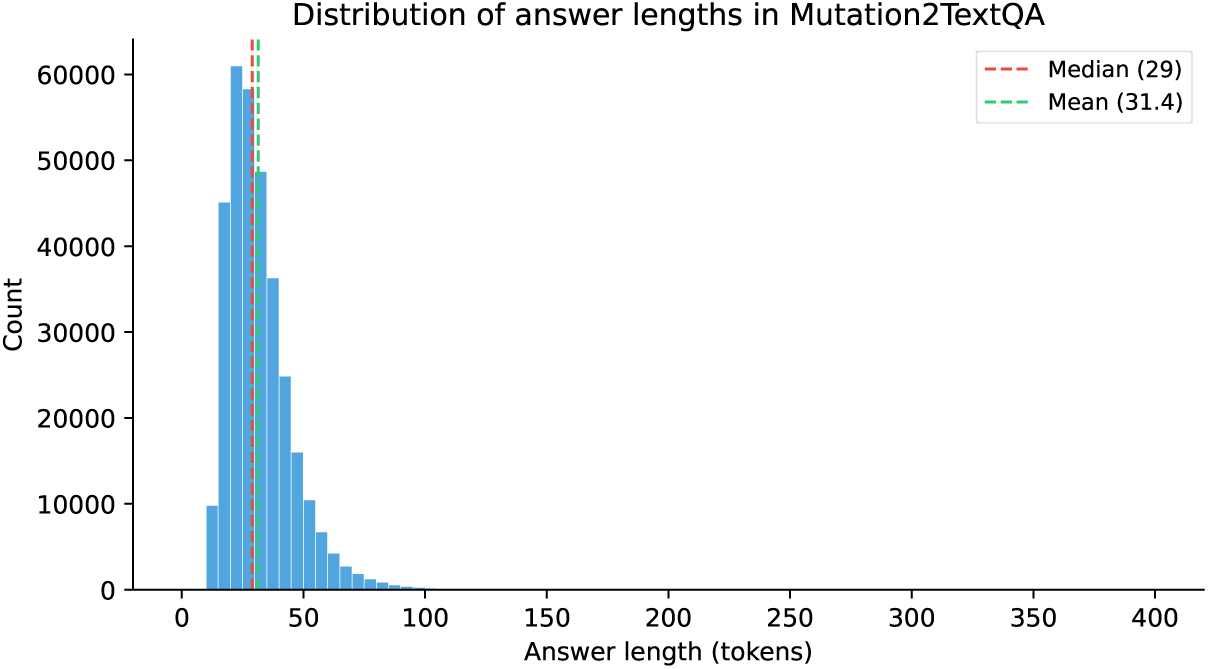
Average answer length (in words) for MutQA.

#### Structure coverage

The benchmark spans **5,437** UniProt accessions, which together index **40,815** PDB entries. **3,325** (61.2%) accessions have at least one PDB structure; **5,339** (98.2%) have an AlphaFold model; and **5,392** (99.2%) have at least one structural source. Only **45** accessions (0.8%) lack both. The structure-evaluable subset covers 4,246 accessions, 63.6% of them with at least one PDB structure. Each PDB list is stored verbatim as returned by the UniProt cross-references API; the AlphaFold identifier follows the canonical AF-{ACC}-F1 naming.

#### Mutation–feature overlap

The benchmark contains **121,170** unique (UniProt, mutation) pairs. Mutations are reported in their original literature notation, which spans HGVS protein, HGVS coding (c.), and free-form descriptions. We mapped each mutation to a 1-indexed protein position whenever the surface form was unambiguous: **50,883** pairs (42.0%) yielded a parseable protein-level residue, while the remaining 70,287 (58.0%) were coding-only, genomic, or otherwise non-positional and are released without a position. Among the 50,496 structure-evaluable pairs, **26,559** (52.6%) fall *within* at least one annotated UniProt feature and **23,937** (47.4%) lie outside any annotated feature; 3,118 of the 4,246 accessions have at least one mutation inside a feature.

#### Feature-type distribution

Table 20 summarises the UniProt feature types that overlap mutated positions in the structure-evaluable subset. Structural domains and annotated regions dominate, followed by disulfide bonds, transmembrane segments and repeats; functional-site annotations (binding sites, active sites, motifs) are much rarer. Filtering on these annotations is useful when constructing structure-or mechanism-focused evaluations.

**Table 20:** UniProt feature types overlapping the mutated residue in the structure-evaluable subset, ranked by the number of (UniProt, mutation) pairs. A single mutation can overlap several feature types and is counted in each. Types with fewer than 50 pairs are omitted.

| Feature type | Overlapping mutations |
| --- | --- |
| Domain | 11,167 |
| Region | 9,306 |
| Disulfide bond | 3,990 |
| Transmembrane | 2,821 |
| Repeat | 2,497 |
| Compositional bias | 1,384 |
| Binding site | 822 |
| Coiled coil | 748 |
| DNA binding | 727 |
| Zinc finger | 567 |
| Signal | 298 |
| Motif | 265 |
| Site | 101 |
| Glycosylation | 87 |
| Active site | 78 |

#### Intended use

Researchers can select the structure subset from the main QA records via the `uniprot_primary` and `mutation_position` fields, or restrict to `structure_evaluable` QA pairs. We anticipate three primary uses: (i) restricting evaluation to QA pairs whose mutation lies in a functionally annotated region, where structural reasoning is most informative; (ii) augmenting model inputs with PDB or AlphaFold coordinates to study structure-conditioned mutation reasoning; and (iii) stratifying error analyses by feature type to test whether models systematically over-or under-perform on mutations in particular structural contexts (e.g., active sites versus disordered regions).

### D.2 Example QA Pairs

This appendix exhibits six representative records drawn from MutQA, sampled (RANDOM_SEED = 42) to span the major question types and all three grounding tiers (cross-grounded, grounded, ungrounded). For each record we show the gene–mutation pair, the source PMID, the natural-language question, both phase-1 (*A*_1_) and phase-2 (*A*_2_) independently-generated answers, the bidirectional cross-validation grounding tier, and the first cited source-text sentence. Together these examples illustrate (i) the format of released records, (ii) the range of question phrasings, and (iii) what convergent vs. partially-convergent vs. failed cross-validation actually looks like in practice.

**Example 1 — Function (cross-validated). Gene/Mutation**: *PRKN*, p.R334C **PMID:** 30994895

**Q:** What is the functional effect of the p.R334C mutation on mitophagy?

*A*_1_: The p.R334C mutation showed an almost three-fold increase in mitophagy.

*A*_2_: The p.R334C mutation shows an almost three-fold increase in mitophagy compared to wild-type Parkin.

**Cited sentence:** “*p.Q34R and p.A46T displayed decreased mitophagy, whereas p.R334C showed an almost three-fold increase (Fig. 1B).*”

**Figure 13:**
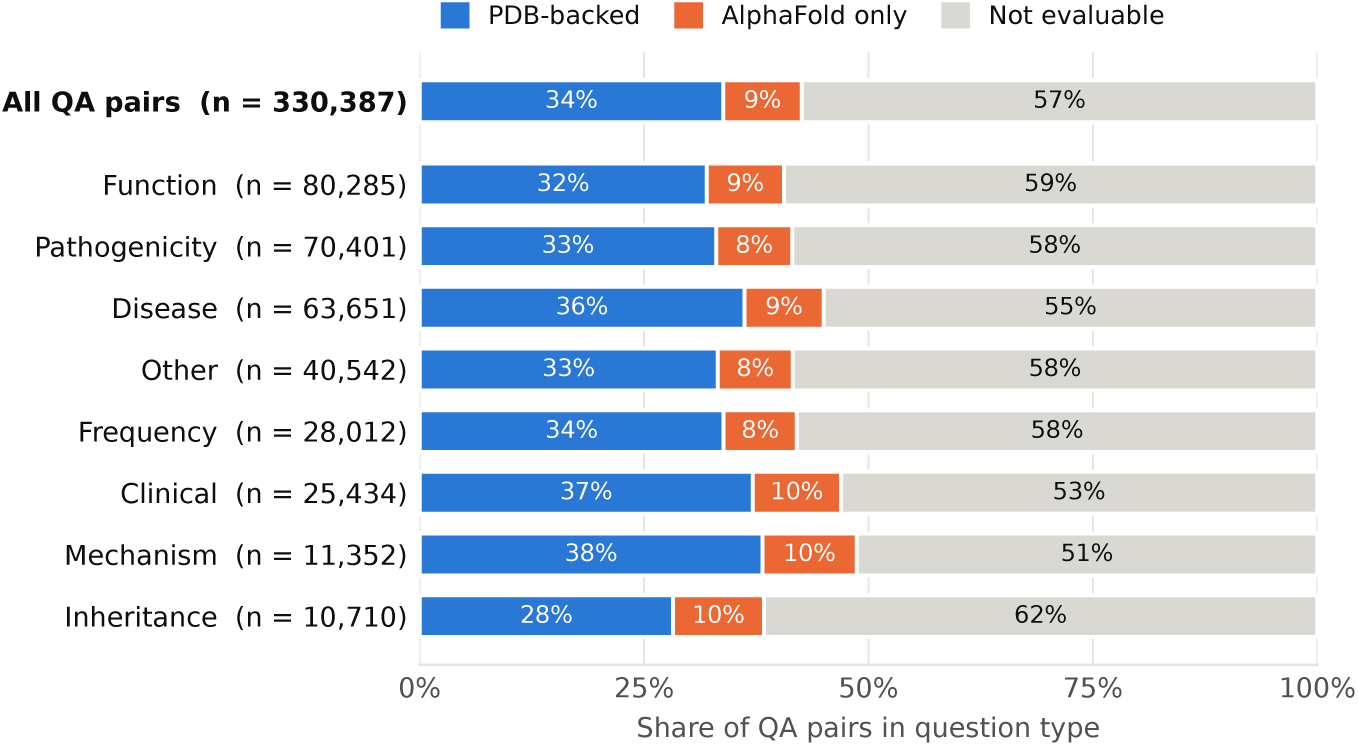
UC4/UC5 structure-subset coverage by question type. Each bar decomposes the share of QA pairs into PDB-backed (blue), AlphaFold-only (orange), and not evaluable (gray, where either the structure or the residue-level mutation position is unavailable). Overall, 33.8% of benchmark QA pairs are PDB-backed, 8.8% AlphaFold-only and 57.4% not evaluable. The PDB:AlphaFold ratio is roughly 3–4:1 across the main question types, reflecting the underlying UniProt → structure mapping rather than any question-type-specific bias.

**Example 2 — Pathogenicity (cross-validated). Gene/Mutation:** *SP8*, rs397514369 **PMID:** 12618081

**Q:** What is the predicted pathogenicity of the variant rs397514369 based on its Meta-SNP score?

*A*_1_: The variant rs397514369 is classified as Pathogenic/Likely pathogenic (P/LP) and is predicted to be disease-causing with a Meta-SNP score of 0.799.

*A*_2_: The variant rs397514369 is predicted to be pathogenic based on its Meta-SNP score.

**Cited sentence:** (Tabular data; see source for full Meta-SNP/MutPred row for rs397514369.)

**Example 3 — Disease (cross-validated). Gene/Mutation:** *NTRK1*, c.2197G*>*A **PMID:** 25984678

**Q:** What condition might be caused by the NTRK1 mutation c.2197G*>*A?

*A*_1_: Bioinformatics analyses predict that the NTRK1 mutation c.2197G*>*A might cause CIPA.

*A*_2_: The NTRK1 mutation c.2197G*>*A (p.Gly714Asp) is associated with congenital insensitivity to pain with anhidrosis (CIPA).

**Cited sentence:** “*Sequencing analysis revealed an inherited novel mutation, c.1635G>C, and a novel de novo mutation, c.2197G>A, in the NTRK1 gene.*”

**Example 4 — Frequency (cross-validated). Gene/Mutation:** *TP53*, c.215G*>*C PMID: 25925845

**Q:** What is the frequency of the c.215G*>*C mutation in cases?

*A*_1_: The c.215G*>*C mutation was found in a heterozygous/homozygous state in 41.7% (n = 15) of cases.

*A*_2_: The c.215G*>*C mutation was found in 41.7% of cases.

**Cited sentence:** “*The c.215G>C [p.Arg72Pro] polymorphism was found in heterozygous/homozygous state in 41.7% (n* = *15) of cases.*”

**Example 5 — Mechanism (one-way validated). Gene/Mutation:** *PMS2*, 1646T*>*C PMID: 15365995

**Q:** What is the predicted impact of the 1646T*>*C (p.Leu549Pro) mutation on the PMS2 binding domain?

*A*_1_: The 1646T*>*C (p.Leu549Pro) mutation is predicted to negatively impact the PMS2 binding domain, as it is located within this domain and is marked as “No” for PMS2 binding domain.

*A*_2_: The 1646T*>*C (p.Leu549Pro) mutation is located in the PMS2 binding domain but does not cause a polarity change.

**Note:** This example is in the *one-way validated* (one-direction) tier rather than *cross-grounded*: *A*_1_ infers a negative functional impact while *A*_2_ describes the variant’s structural location without committing to an effect direction. The two answers do not contradict but draw different conclusions from the available evidence—exactly the pattern the bidirectional check is designed to surface.

**Example 6 — Clinical (non-validated). Gene/Mutation:** *CD27*, p.C53Y **PMID:** 32603431

**Q:** What clinical features are associated with the CD27 p.C53Y mutation?

*A*_1_: Patients with the CD27 p.C53Y mutation presented with symptoms including LRTI, sepsis, phlegmons, HLH, LPD, recurrent infections, URTI, meningitis, oral ulcers, uveitis, CAEBV, and DLBCL (EBV+).

*A*_2_: The CD27 p.C53Y mutation is associated with clinical features such as EBV-related symptoms, lymphoproliferation, and lymphoma.

**Note:** This example is in the *ungrounded* tier and is excluded from the high-confidence subset. *A*_1_ enumerates a long list of specific symptoms; *A*_2_ summarizes at a higher level of abstraction. While not contradictory, the answers operate at different specificity levels, and the bidirectional judge cannot confirm that the specific symptom list in *A*_1_ is recoverable from the same evidence that supports *A*_2_. Records like this are released alongside the high-confidence subset (with the *ungrounded* tier label preserved) so that downstream users can use them as a negative-control test for model abstention behavior.

## E Extended Discussion on Methodology

### E.1 Technical Implementation

### E.2 Evaluation Metrics

#### BLEU Scores

[Papineni et al., 2002]. The BLEU (Bilingual Evaluation Understudy) score relies on n-gram matching to calculate the performance of the generated text. The BLEU score is a precision-based metric that quantifies the number of n-grams in the generated text that are also mentioned in the ground-truth text. For example, BLEU-2 denotes bigram matching.

#### ROUGE Scores

[Lin, 2004].Similarly, the ROUGE (Recall-Oriented Understudy for Gisting Evaluation) score also focuses on sequence matching. As opposed to BLEU, the ROUGE score is recall-based which calculates the amount of n-grams from the ground truth that are captured by the generated response. **ROUGE-1** denotes unigram matching, **ROUGE-2** bigrams matching, and **ROUGE-L** denotes longest common subsequence matching.

#### METEOR Score

[Banerjee and Lavie, 2005]. METEOR (Metric for Evaluation of Translation with Explicit ORdering) weighs recall and precision while performing n-gram matching. METEOR also captures high-level semantic similarity by applying stemming and synonym matching.

#### CIDEr

[Vedantam et al., 2015]. CIDEr (Consensus-based Image Description Evaluation) measures the similarity of a candidate caption to a set of human references by computing TF–IDF weighted n-gram overlaps. Unlike BLEU or ROUGE, it emphasizes consensus by rewarding informative words that appear consistently across human descriptions, making it better aligned with human judgments. CIDEr was validated on new datasets (PASCAL-50S, ABSTRACT-50S) with 50 captions per image and shown to outperform existing metrics in correlating with consensus-based human evaluations.

#### LLM-as-judge score (UC1/UC2)

Reference-overlap metrics rely on surface matching, so for UC1 and UC2 we also score each prediction with three LLM judges from different developers: DeepSeek-v3.2, Gemini-2.0-Flash and GPT-4o-mini, each at temperature 0 with at most 200 output tokens. A judge sees the question, the ground-truth answer and the prediction, and returns three integer scores from 0 to 5: *correctness* (factual accuracy of the predicted information), *completeness* (how much of the ground-truth content is covered) and *relevance* (how directly the prediction addresses the question). The reported judge mean averages the three dimensions and then the three judges, over a 200-item stratified subset per use case. For UC1 the question slot holds the per-item instruction naming the mutation (e.g., “Describe the effect of the A270D mutation in this protein.”); for UC2 it holds the default instruction “Describe the effect of this mutation.” The verbatim prompt is:

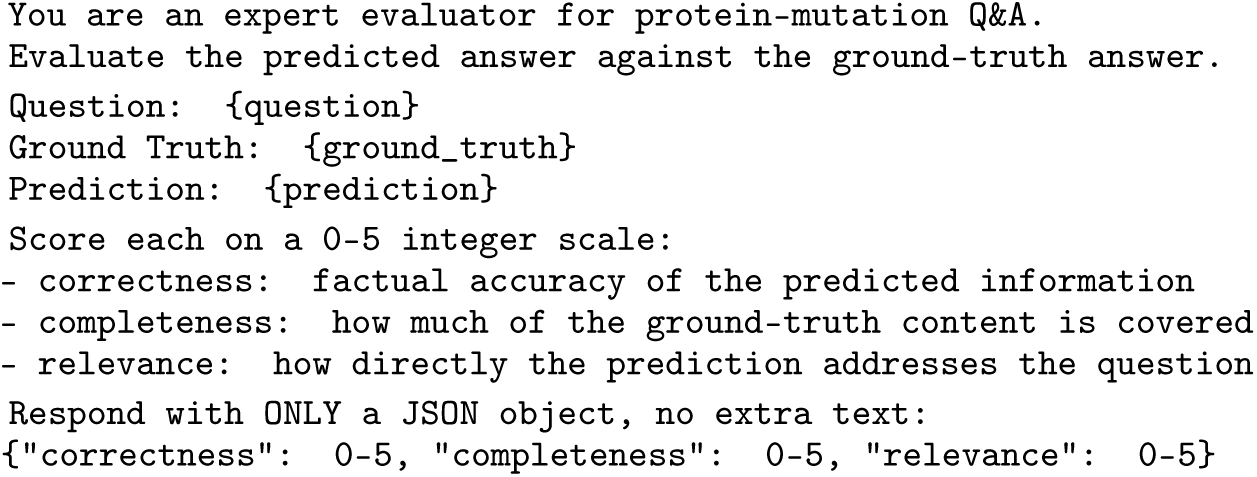

Table 21 reports each judge separately. Every judge reproduces the fine-tuning gain, although absolute calibration differs (on UC1, pretrained outputs score 1.29 under GPT-4o-mini and 0.72 under Gemini). Correctness alone gives the same direction and a comparable size (UC1 0.74 → 1.31; UC2 1.33 → 2.49).

**Table 21:** Judge sensitivity: mean score over the three dimensions (0–5) per judge, 200 items per use case.

| Judge | UC1 |  |  | UC2 |  |  |
| --- | --- | --- | --- | --- | --- | --- |
| | Pre | FT | $\Delta$ | Pre | FT | $\Delta$ |
| DeepSeek-v3.2 | 0.88 | 1.63 | +0.75 | 1.44 | 2.90 | +1.46 |
| Gemini-2.0-Flash | 0.72 | 1.37 | +0.65 | 1.25 | 2.95 | +1.70 |
| GPT-4o-mini | 1.29 | 1.85 | +0.56 | 1.66 | 2.61 | +0.95 |
| Mean | 0.96 | 1.62 | +0.66 | 1.45 | 2.82 | +1.37 |

### E.3 Human Evaluation of MutaPLM Outputs (UC1/UC2)

Four annotators with molecular-biology training scored pretrained and fine-tuned MutaPLM outputs for items drawn from the 200-item judge subset of each use case. Each annotator rated 25 items per use case: a 10-item block shared by all four and 15 items seen by no one else, giving 100 ratings over 70 unique items per use case. For each item the annotator saw the question, the reference answer and the two outputs in randomized order (seed 42), scored each output on the 0–2 correctness rubric of C.3.1, and stated a preference or a tie. Means and tests are computed over the 100 ratings, so each shared item counts once per annotator. Pretrained and fine-tuned scores are compared with a two-sided Wilcoxon signed-rank test, preferences with a two-sided sign test on non-tied ratings, and human scores (averaged per item) are correlated with the judges’ mean correctness score over 140 answers (70 items × 2 systems).

**Table 22:** Human evaluation of MutaPLM outputs (0–2 correctness, 100 ratings per use case). CI: bootstrap 95% interval on Δ. Fully correct: share of ratings scored 2. Preference: fine-tuned / pretrained / tie.

| | Pre | FT | $\Delta$ (CI) | $p$ | Fully correct (Pre / FT) | Preference | $p$ | $\rho$ |
| --- | --- | --- | --- | --- | --- | --- | --- | --- |
| UC1 | 0.12 | 0.36 | +0.24 (0.11–0.37) | $8.2 \times 10^{-4}$ | 0% / 10% | 23 / 8 / 69 | 0.011 | 0.52 |
| UC2 | 0.11 | 0.62 | +0.51 (0.34–0.68) | $2.4 \times 10^{-6}$ | 3% / 21% | 44 / 6 / 50 | $3.2 \times 10^{-8}$ | 0.60 |

### Inter-annotator agreement

On the shared 10-item blocks, mean pairwise exact agreement on fine-tuned outputs is 63.3% for UC1 (mean pairwise *κ* = −0.04; Fleiss *κ* = −0.08) and 71.7% for UC2 (*κ* = 0.46; *κ_w_* = 0.74; Fleiss *κ* = 0.45). On pretrained outputs agreement is 95% in both use cases, but nearly every score is 0, so *κ* is undefined or zero. Fleiss *κ* on preferences is −0.01 (UC1) and 0.55 (UC2). UC1 agreement is at chance, so the UC1 human result is weaker evidence than UC2’s; both are based on small shared blocks.

### E.4 Training-free Retrieval Evaluation

#### Corpus and targets

The retrieval corpus is the full 401,406-pair release (all validation tiers) after a gene-assignment purity filter (purity ≥ 0.8), leaving 199,168 QA pairs over 4,846 genes. Of its 63,481 variants, 5,289 (8.3%) have more than five QA pairs and are eligible; 500 target questions were sampled from these and used for all four models (403 cross-validated, 25 one-way validated, 72 non-validated).

#### Context

For each target, the context is the variant’s other QA pairs (minimum 3, median 11, maximum 20), after removing pairs whose question or answer has token-Jaccard overlap ≥ 0.85 with the target’s question or answer, or matches it exactly after normalization.

#### Generation and judging

Models are called with temperature 0, seed 42, at most 1,500 output tokens, low reasoning effort and no tools. The system prompt asks for an answer of at most three sentences or the string “INSUFFICIENT INFORMATION”. The judge (Claude Haiku 4.5, temperature 0) sees only the question, reference and prediction. All prompts are released verbatim.

**Table 23:**
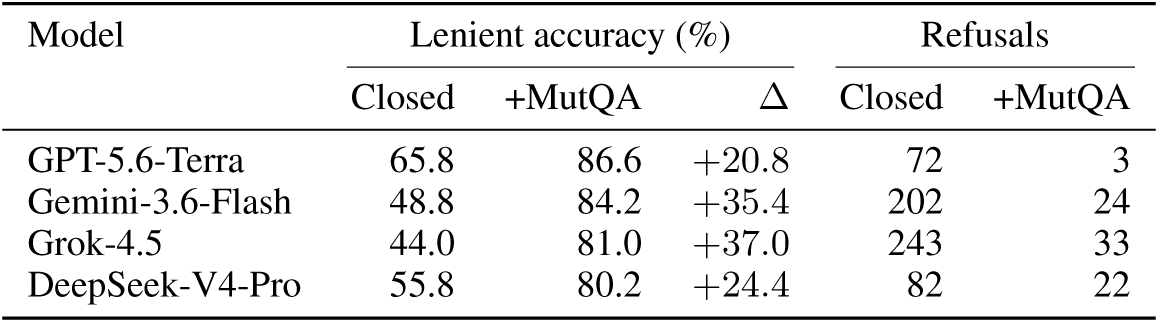
Retrieval evaluation: lenient accuracy (score ≥ 1) and refusals (of 500) per model; strict accuracy is in. Table 5.

| Model | Lenient accuracy (%) |  |  | Refusals |  |
| --- | --- | --- | --- | --- | --- |
| | Closed | +MutQA | $\Delta$ | Closed | +MutQA |
| GPT-5.6-Terra | 65.8 | 86.6 | +20.8 | 72 | 3 |
| Gemini-3.6-Flash | 48.8 | 84.2 | +35.4 | 202 | 24 |
| Grok-4.5 | 44.0 | 81.0 | +37.0 | 243 | 33 |
| DeepSeek-V4-Pro | 55.8 | 80.2 | +24.4 | 82 | 22 |

**Table 24:** Retrieval controls: change in strict accuracy (percentage points). n.s.: McNemar *p* ≥ 0.05.

| Condition | $n$ per model | $\Delta$ strict (Terra / Gemini / Grok / DeepSeek) |
| --- | --- | --- |
| Main analysis (overlap $\geq 0.85$ removed) | 500 | +20.0 / +24.6 / +28.6 / +21.8 |
| Cross-validated targets only | 403 | +20.8 / +27.5 / +30.8 / +21.8 |
| Question overlap $\geq 0.60$ removed | 386 | +19.4 to +28.0 |
| Answer overlap with reference $\geq 0.40$ removed | 276 | +4.7 (n.s.) / +12.0 / +13.4 / +10.5 |
| Answer overlap with reference $\geq 0.30$ removed | 149 | +5.4 (n.s.) / +8.7 / +8.1 (n.s.) / +8.7 |
| Same-gene, other-variant context (Terra only)<sup>a</sup> | 250 | +4.0 (n.s., $p = 0.21$) |

<sup>a</sup>Equal number of QA pairs about other variants of the same gene; scored by a three-judge consensus.

#### By mutation class

Pooled over the four models (2,000 answers; 500 unique questions), strict accuracy rises for substitutions from 33.7% to 59.2% (*n* = 1,628), indels from 36.6% to 53.1% (*n* = 224), splice/intronic variants from 38.1% to 64.3% (*n* = 84) and nonsense variants from 56.3% to 59.4% (*n* = 64, 16 unique questions). Classes are assigned from mutation notation, so stop-gains written at the DNA level are counted as substitutions.

#### Human check

Four annotators scored a blinded, paired subset of 100 targets (25 per model, stratified by question type) on the same 0–2 rubric; each rated 25 items and a 20-item block was rated by all four (160 ratings). Per-item scores are the rounded mean over raters. Human strict accuracy rises from 29.0% to 52.0% (+23.0 points, McNemar *p* = 1.2 × 10*^−^*^4^) and lenient accuracy from 49.0% to 75.0% (*p* = 4.2 × 10*^−^*^5^); using a majority vote per item instead gives 26% to 49%. The Haiku judge on the same 100 items gives 35% to 55%. Over the 320 rated answers, human and judge agree exactly on 72.2% and on the strict cut on 81.2%; mean scores are 1.03 (human) and 1.06 (judge), with humans harsher on 49 answers and the judge on 40. Annotators preferred the retrieval-conditioned answer in 74 of 158 ratings and the closed-book answer in 30 (sign test *p* = 1.9 × 10*^−^*^5^). On the shared block, mean pairwise exact agreement is 60.8% (closed book) and 65.0% (+MutQA), with mean pairwise *κ* of 0.40 and 0.43 (Fleiss 0.39 and 0.43).

## F Deterministic Sequence-Level Audit

This appendix expands on Section 3: the deterministic verifier that runs as the final filter of the curation pipeline and removes Q&A whose Phase-1 or Phase-2 answer makes a sequence-derivable claim that contradicts the parsed mutation. The verifier comprises two HGVS-aware sub-modules: a **protein-coding side** that ingests p.-prefixed descriptors against the UniProt reference sequence and grades each answer against *ten* regex-driven claim families — residue identity, position, mutation kind, region class (*N* /middle/*C*-terminal third), and the per-residue physicochemical properties charge, polarity, Kyte–Doolittle hydrophobicity, Tsai *et al*. [1999] side-chain volume, aromaticity, and phospho/glycosylation capability — and a **DNA-coding side** (an expansion introduced for this release) that ingests c.- and bare-nucleotide descriptors against the GRCh38 genome and the matched canonical transcript, adding *seven* codon-mechanical and feature-level claim families: reference and alternate allele identity, substitution class, the predicted wild-type and mutant amino acids inferred from the codon, premature-stop consequence, splice donor / acceptor / region overlap, and UniProt feature overlap (active site, binding site, disulfide, domain, topology, transmembrane). The DNA side recovers the 232,311 Q&A rows whose mutation strings are not protein-level resolvable and therefore fell out at Stage 1 of the protein-only funnel.

### Ground-truth dictionary (protein side)

For each Q&A we parse the HGVS descriptor into (*wt, pos, mut, kind*) and compute, from *ref_sequence* and the per-residue property tables: protein length; *region_class* (N/middle/C-terminal third); the change facts *charge_change*, *polarity_change*, *hydrophobicity_change*, *size_change*, *aromaticity_change*; and special-residue flags for proline rigidity, glycine flexibility, cysteine (disulfide capability), aromatic side chains, S/T/Y phospho-sites, and asparagine in an N-X-S/T glycosylation sequon. Charge is assigned at physiological pH from the canonical sidechain pKa values [Nelson and Cox, 2017], hydrophobicity from Kyte–Doolittle [Kyte and Doolittle, 1982], side-chain volume from Tsai *et al*. [1999], and aromaticity following Burley & Petsko [1985] (including histidine).

### Ground-truth dictionary (DNA side)

DNA-coding descriptors are resolved by aligning the cited transcript to GRCh38, locating the genomic coordinate of the substituted base, and recovering the codon, reading frame, and reference and alternate alleles. From this we derive: the *dna_ref_base* and *dna_alt_base* identities; the substitution kind (*transition* vs. *transversion*); the predicted codon-level amino-acid change (*predicted_wt_aa*, *predicted_mut_aa*); whether the substitution introduces a *premature_stop* and, if so, the post-truncation length; and *splice_donor* / *splice_acceptor* / *splice_region* flags from the canonical exon-intron boundaries. The codon-level prediction routes the resulting amino-acid change through the protein-side property tables, so a DNA-level claim about a residue change inherits the same charge / polarity / size scoring as a protein-side claim. We additionally project the encoded position onto the matched UniProt entry to enable feature-level checks — *uniprot_active_site*, *uniprot_binding_site*, *uniprot_disulfide*, *uniprot_domain*, *uniprot_topology*, *uniprot_transmembrane* — so that DNA-side answers asserting location within a functional region can be verified against UniProt.

### Trigger discovery

Trigger regexes for the protein-side claim families (mutation kind, residue identity, position, charge, polarity, hydrophobicity, size, region, special residue, protein length) were derived from a ten-slice pattern-discovery sweep over the Phase-2 gold set. Ten parallel Claude agents each read a contiguous ∼1,960-row slice and reported up to fifty verbatim phrasings per category — hedge expressions, false-friend / paralog references, and out-of-scope claim types (NMD, splicing, ΔΔ*G*, kinetic constants, expression, localization) included. The output was compiled into a single trigger module; every regex pattern is annotated in-place with the slice excerpts that motivated it, so every trigger is grounded in observed phrasings rather than *a priori* dictionaries. The DNA-side trigger module was bootstrapped by the same procedure, with an additional sweep targeted at codon-level and splice-junction phrasings. Out-of-scope claim families are returned as not_applicable (no trigger emitted) rather than scored against an external annotation we cannot verify deterministically. Each extracted claim also passes through three context filters, applied uniformly across both sub-verifiers: a hedge filter (“would”, “could”, “approximately”), a hypothetical filter (analog / metabolite / modification discussion), and a negation filter that flips polarity for constructions like “no disulfide” or “loss of glycosylation”.

### Cleaning funnel

Table 28 reports the cumulative effect of the combined verifier on the 411,060-pair pre-audit working set: all validation tiers, after the answerability and consistency filters (Appendix C.1). Stage 1 separates protein-coding from DNA-coding QA pairs; Stage 2 drops QA pairs flagged as contradicted by either sub-verifier on either the Phase-1 or the Phase-2 answer. Of the 9,654 QA pairs removed, 7,354 are flagged by the protein side and a further 2,300 by the DNA side (Table 28).

**Table 25:**
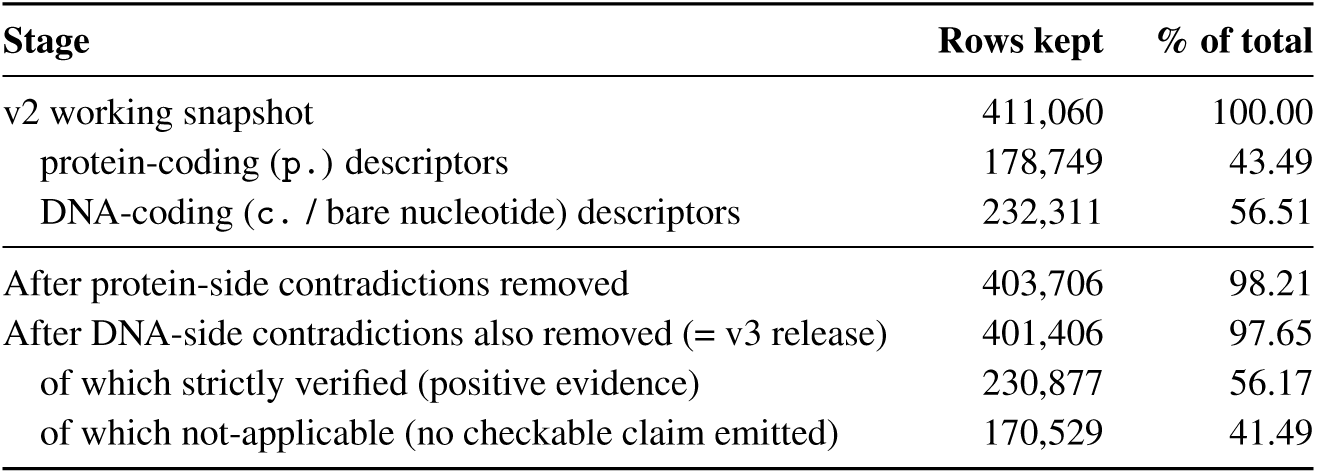
Cleaning funnel of the deterministic audit, applied to the v2 working set after cross-validation and Phase-3 answerability. Across both sub-verifiers, 9,654 rows are removed (2.35% of the v2 working set). Strictly-verified and not-applicable counts are the union over both sub-verifiers on the surviving rows.

**Figure 14:**
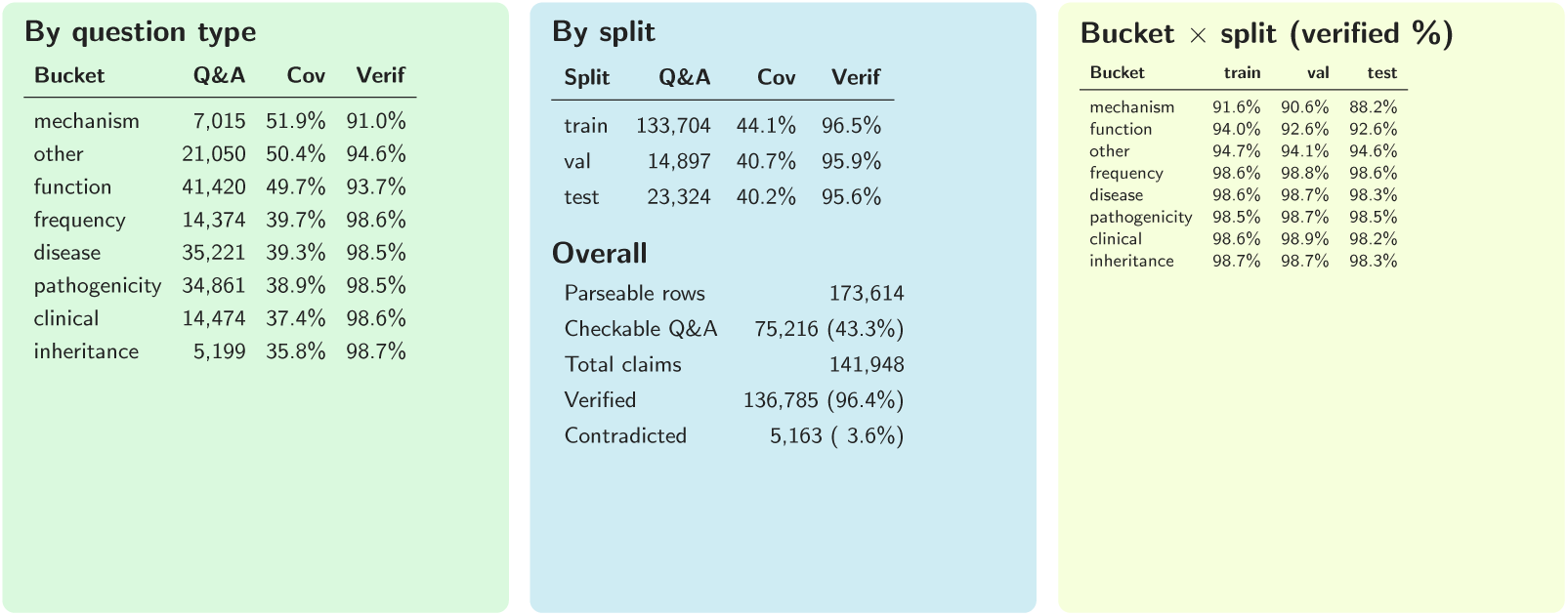
Verifier breakdown over the v3 release, by question type, by data split, and by bucket × split. MECHANISM is the highest-coverage / lowest-verified bucket because it is where answers commit to specific biophysical claims; the descriptive buckets sit at ∼ 98 % verified because their checkable content is mostly the inline restatement of the residue substitution. Splits agree within 1.5 pp.

### Per-claim verification (protein side)

Table 29 reports the per-claim-family verified rate over all firings on the protein side. Three tiers emerge: residue identity and structural special-residue claims at ≥ 88 %; soft physicochemical / positional claims at 65–82 %; and a convention-sensitive cluster below 40 %.

### Per-claim verification (DNA side)

Table 30 reports the same breakdown for the DNA-side verifier. The high-signal tier is dominated by the codon-mechanical claims (*dna_ref_base*, *dna_alt_base*, *predicted_wt_aa*, *predicted_mut_aa*) which cap above 95 % because they are recoverable directly from the parsed descriptor and the transcript alignment. UniProt-feature claims and splice-region flags also sit in the high tier as they are emitted only when the cited region unambiguously overlaps a UniProt annotation.

### Failure modes caught

On the protein side, three error patterns dominate the contradicted slice. *Region mislocalization:* unhedged “N-terminal” / “C-terminal” assertions that disagree with the position-class fact (e.g. APP p.Val604Met described as “located at the N-terminal region” although residue 604/770 sits in the C-terminal third, just upstream of the A*β* sequence). *Physicochemical over-claims:* hydrophobicity flips asserted for L→P, V→T, or M→L substitutions when both residues fall on the same side of the Kyte–Doolittle scale, and analogous *charge changes described in the wrong direction* (e.g. a basic→polar substitution narrated as introducing a negative charge), caught by the *hydrophobicity_change* and *charge_change* claims. *Mechanistically-impossible special-residue claims:* loss of an N-X-S/T glycosylation site asserted at residues that are not asparagine, or loss of an S/T/Y phospho-site at non-S/T/Y residues. The verifier also exposes a small *taxonomy-convention* band — e.g. in-frame internal deletions described as producing a “truncated protein” (SQSTM1 p.Gly351_Pro388del → “a truncated protein of 402 amino acids”), biologically accurate but flagged by a deterministic taxonomy that reserves truncation for stop-codon-induced events.

**Table 26:**
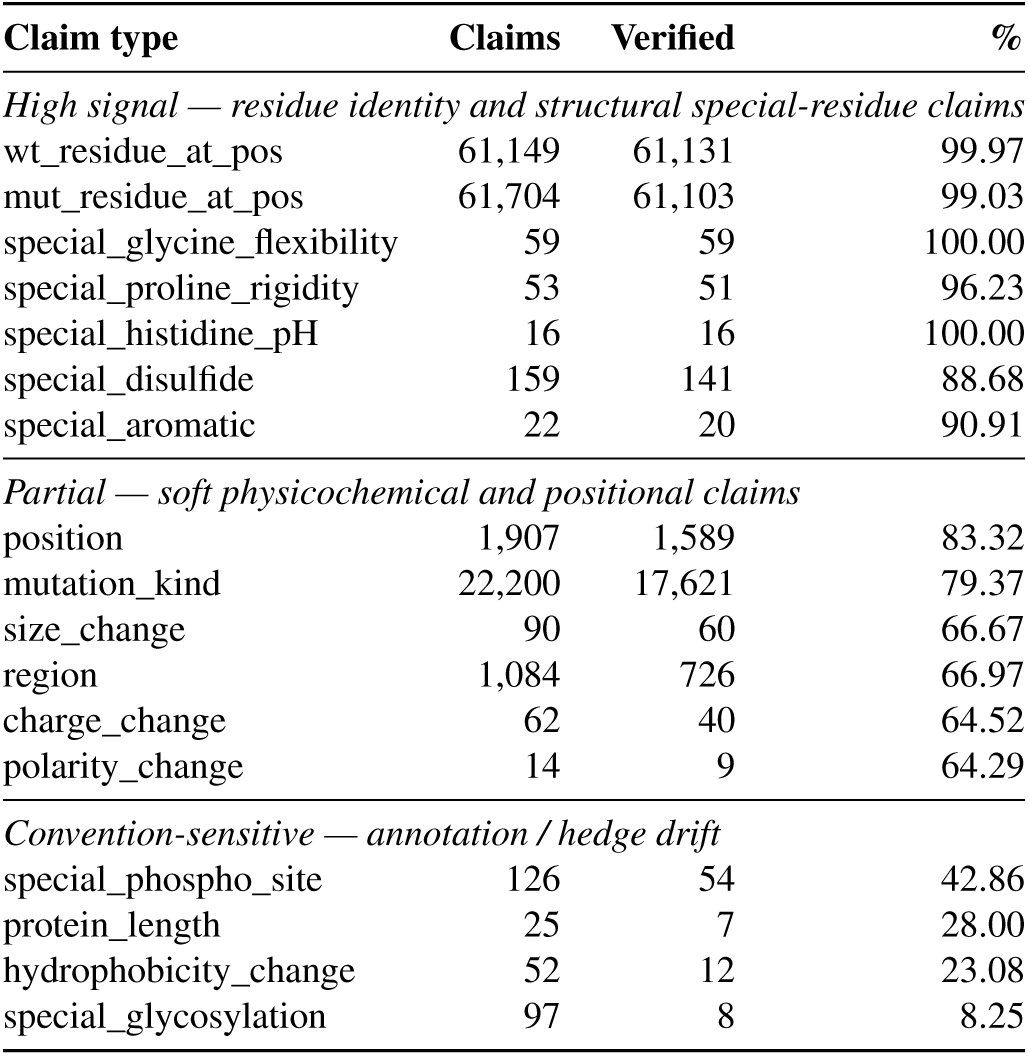
Per-claim verification rate on Phase-1 (qagen) answers, protein side. Same tiering pattern holds on Phase-2 within ±2 pp.

**Table 27:**
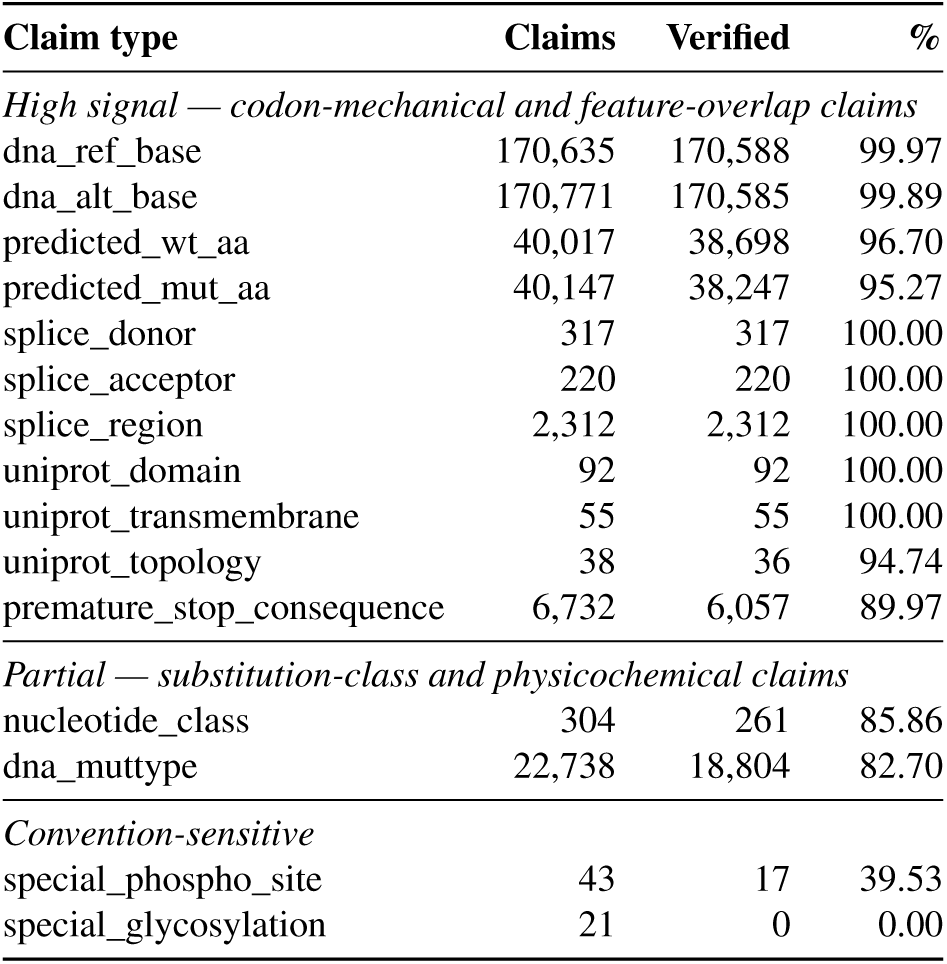
Per-claim verification rate on Phase-1 (qagen) answers, DNA side. Phase-2 rates are within ±1 pp on the high-signal tier.

The DNA side surfaces a complementary set of failure modes that the protein-only verifier could not see. *Wrong reference base:* answers restate a c.-level substitution with the cited reference allele swapped (e.g. c.1138G>A answered as “a C-to-T transition”), caught by *dna_ref_base* / *dna_alt_base*. *Codon-translation errors:* answers infer the wrong amino-acid consequence from the cited DNA change (a non-synonymous change asserted to be silent, or the predicted mutant residue named incorrectly), caught by *predicted_wt_aa* and *predicted_mut_aa*. *Mis-attributed splice impact:* answers describe a coding-region substitution as “disrupting the splice donor” when the substituted base is not within the canonical ±6 nt exon-intron window, caught by the *splice_\** flags. *Truncation-length over-counts:* stop-introducing substitutions described as producing a protein of an incorrect post-truncation length, caught by *premature_stop_consequence*.

### Ground-truth dictionary (protein side)

For each Q&A we parse the HGVS descript or into (*wt, pos, mut, kind*) and compute, from *ref_sequence* and the per-residue property tables: protein length; *region_class* (N/middle/C-terminal third); the change facts *charge_change*, *polarity_change*, *hydrophobicity_change*, *size_change*, *aromaticity_change*; and special-residue flags for proline rigidity, glycine flexibility, cysteine (disulfide capability), aromatic side chains, S/T/Y phospho-sites, and asparagine in an N-X-S/T glycosylation sequon. Charge is assigned at physiological pH from the canonical sidechain pKa values [Nelson and Cox, 2017], hydrophobicity from Kyte–Doolittle [Kyte and Doolittle, 1982], side-chain volume from Tsai *et al*. [1999], and aromaticity following Burley & Petsko [1985] (including histidine).

### Ground-truth dictionary (DNA side)

DNA-coding descriptors are resolved by aligning the cited transcript to GRCh38, locating the genomic coordinate of the substituted base, and recovering the codon, reading frame, and reference and alternate alleles. From this we derive: the *dna_ref_base* and *dna_alt_base* identities; the substitution kind (*transition* vs. *transversion*); the predicted codon-level amino-acid change (*predicted_wt_aa*, *predicted_mut_aa*); whether the substitution introduces a *premature_stop* and, if so, the post-truncation length; and *splice_donor* / *splice_acceptor* / *splice_region* flags from the canonical exon-intron boundaries. The codon-level prediction routes the resulting amino-acid change through the protein-side property tables, so a DNA-level claim about a residue change inherits the same charge / polarity / size scoring as a protein-side claim. We additionally project the encoded position onto the matched UniProt entry to enable feature-level checks — *uniprot_active_site*, *uniprot_binding_site*, *uniprot_disulfide*, *uniprot_domain*, *uniprot_topology*, *uniprot_transmembrane* — so that DNA-side answers asserting location within a functional region can be verified against UniProt.

### Trigger discovery

Trigger regexes for ten claim families (mutation kind, residue identity, position, charge, polarity, hydrophobicity, size, region, special residue, protein length) were derived from a ten-slice pattern-discovery sweep over the Phase-2 gold set. Ten parallel Claude agents each read a contiguous ∼1,960-row slice and reported up to fifty verbatim phrasings per category — hedge expressions, false-friend / paralog references, and out-of-scope claim types (NMD, splicing, ΔΔ*G*, kinetic constants, expression, localization) included. The output was compiled into a single trigger module; every regex pattern is annotated in-place with the slice excerpts that motivated it. Out-of-scope claim families are returned as not_applicable (no trigger emitted) rather than scored against an external annotation we cannot verify deterministically. Each extracted claim also passes through three context filters: a hedge filter (“would”, “could”, “approximately”), a hypothetical filter (analog / metabolite / modification discussion), and a negation filter that flips polarity for constructions like “no disulfide” or “loss of glycosylation”.

### Cleaning funnel

Table 28 reports the cumulative effect of the combined verifier on the 411,060-row data set, Phase-3 answerability-passing snapshot that feeds the deterministic stage). Stage 1 separates protein-coding from DNA-coding rows; Stage 2 drops rows flagged as contradicted by either sub-verifier on either Phase-1 or Phase-2 answer. Of the 9,654 unique row drops, 7,354 are flagged by the protein side (either Phase-1 or Phase-2 contradicted) and 9,614 by the DNA side; the two sets overlap minimally because they cover disjoint mutation notations.

**Figure 15:**
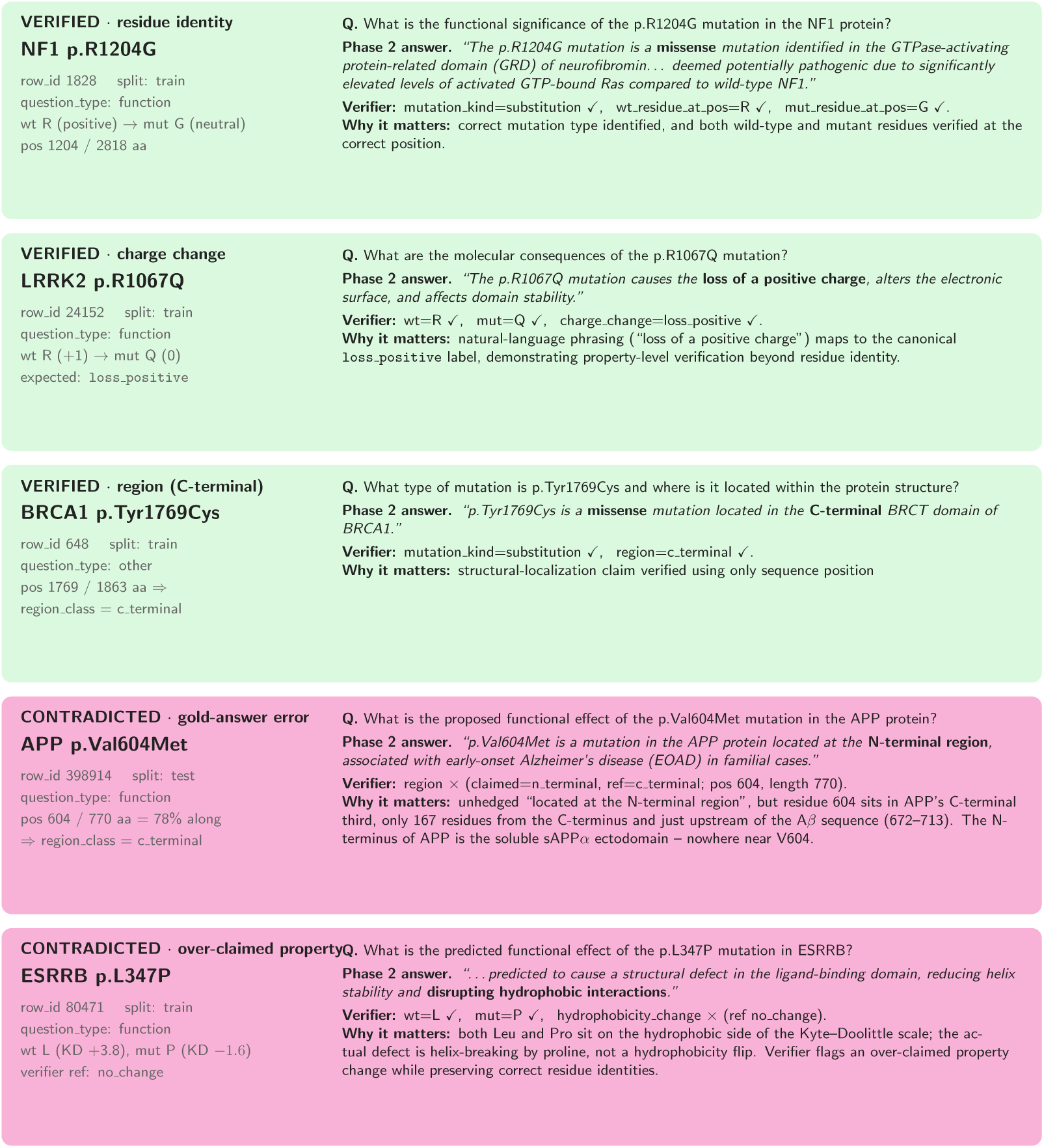
Five representative Q&A from the verifier output, quoted verbatim from the source dataset by row_id. Top three (green) are verified — residue identity (NF1 p.R1204G), property-level charge change (LRRK2 p.R1067Q), and position-class region (BRCA1 p.Tyr1769Cys). Bottom two (pink) are genuine gold-answer errors — an unhedged N-/C-terminal mislocalization (APP V604M placed in the “N-terminal region” though residue 604/770 is in the C-terminal third) and an over-claimed hydrophobicity change.

### Per-claim verification (protein side)

Table 29 reports the per-claim-family verified rate over all firings on the protein side. Three tiers emerge: residue identity and structural special-residue claims at ≥ 88 %; soft physicochemical / positional claims at 65–82 %; and a convention-sensitive cluster below 40 %.

### Per-claim verification (DNA side)

Table 30 reports the same breakdown for the DNA-side verifier. The high-signal tier is dominated by the codon-mechanical claims (*dna_ref_base*, *dna_alt_base*, *predicted_wt_aa*, *predicted_mut_aa*) which cap above 95 % because they are recoverable directly from the parsed descriptor and the transcript alignment. UniProt-feature claims and splice-region flags also sit in the high tier as they are emitted only when the cited region unambiguously overlaps a UniProt annotation.

**Table 28:**
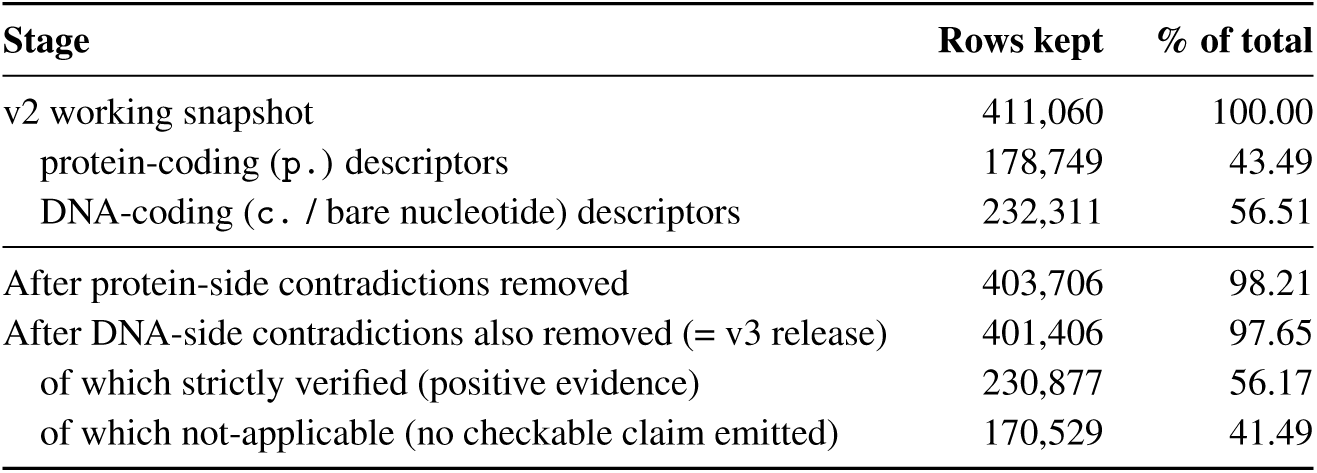
Cleaning funnel of the deterministic audit, applied to the v2 working set after cross-validation and Phase-3 answerability. Across both sub-verifiers, 9,654 rows are removed (2.35% of the v2 working set). Strictly-verified and not-applicable counts are the union over both sub-verifiers on the surviving rows.

**Figure 16:**
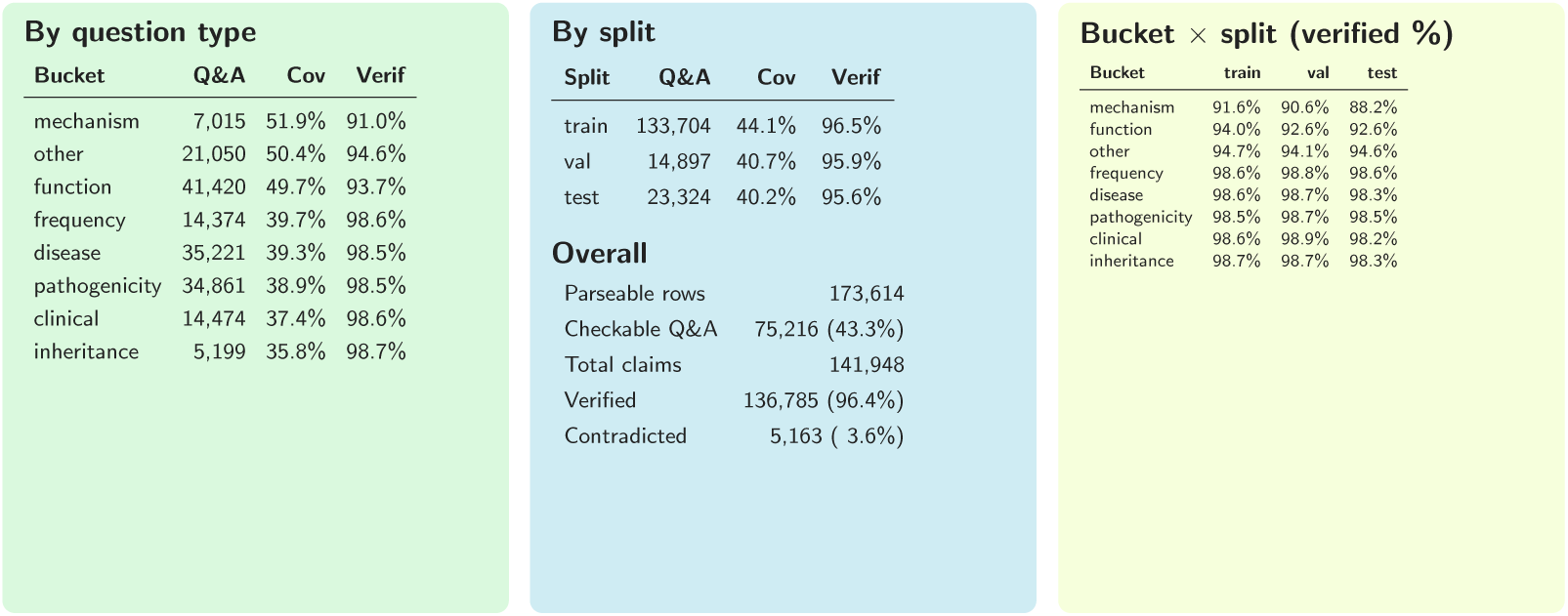
Verifier breakdown over the v3 release, by question type, by data split, and by bucket × split. MECHANISM is the highest-coverage / lowest-verified bucket because it is where answers commit to specific biophysical claims; the descriptive buckets sit at ∼ 98 % verified because their checkable content is mostly the inline restatement of the residue substitution. Splits agree within 1.5 pp.

### Failure modes caught

On the protein side, three error patterns dominate the contradicted slice. *Region mislocalization:* unhedged “N-terminal” / “C-terminal” assertions that disagree with the position-class fact (e.g. APP p.Val604Met described as “located at the N-terminal region” although residue 604/770 sits in the C-terminal third, just upstream of the A*β* sequence). *Hydrophobicity over-claims:* L→P, V→T, or M→L substitutions described as a hydrophobicity flip when both residues fall on the same side of the Kyte–Doolittle scale. *Mechanistically-impossible special-residue claims:* loss of an N-X-S/T glycosylation site asserted at residues that are not asparagine, or loss of an S/T/Y phospho-site at non-S/T/Y residues. The verifier also exposes a small *taxonomy-convention* band — e.g. in-frame internal deletions described as producing a “truncated protein” (SQSTM1 p.Gly351_Pro388del → “a truncated protein of 402 amino acids”), biologically accurate but flagged by a deterministic taxonomy that reserves truncation for stop-codon-induced events.

The DNA side surfaces a complementary set of failure modes that the protein-only verifier could not see. *Wrong reference base:* answers restate a c.-level substitution with the cited reference allele swapped (e.g. c.1138G>A answered as “a C-to-T transition”), caught by *dna_ref_base* / *dna_alt_base*. *Codon-translation errors:* answers infer the wrong amino-acid consequence from the cited DNA change (a non-synonymous change asserted to be silent, or the predicted mutant residue named incorrectly), caught by *predicted_wt_aa* and *predicted_mut_aa*. *Mis-attributed splice impact:* answers describe a coding-region substitution as “disrupting the splice donor” when the substituted base is not within the canonical ±6 nt exon-intron window, caught by the *splice_\** flags. *Truncation-length over-counts:* stop-introducing substitutions described as producing a protein of an incorrect post-truncation length, caught by *premature_stop_consequence*.

**Table 29:**
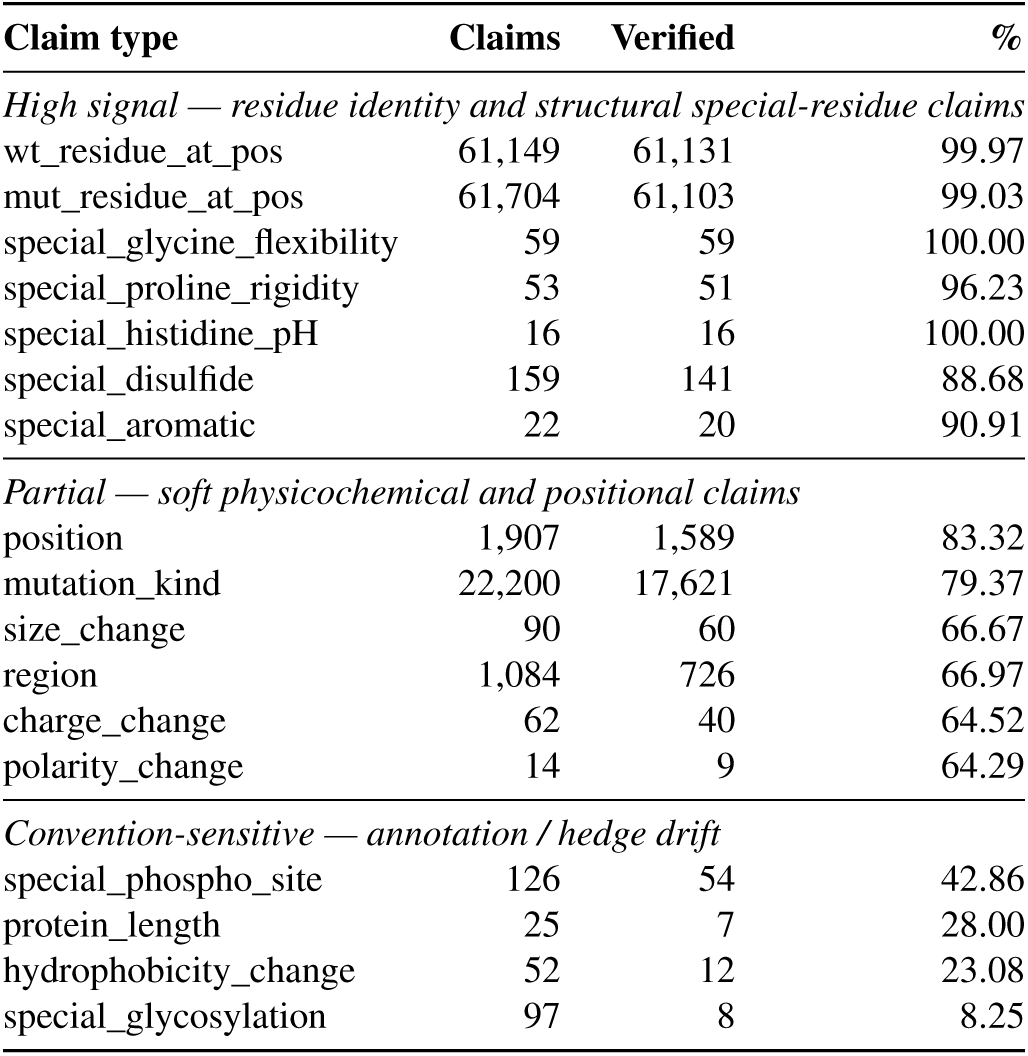
Per-claim verification rate on Phase-1 (qagen) answers, protein side. Same tiering pattern holds on Phase-2 within ±2 pp.

**Table 30:**
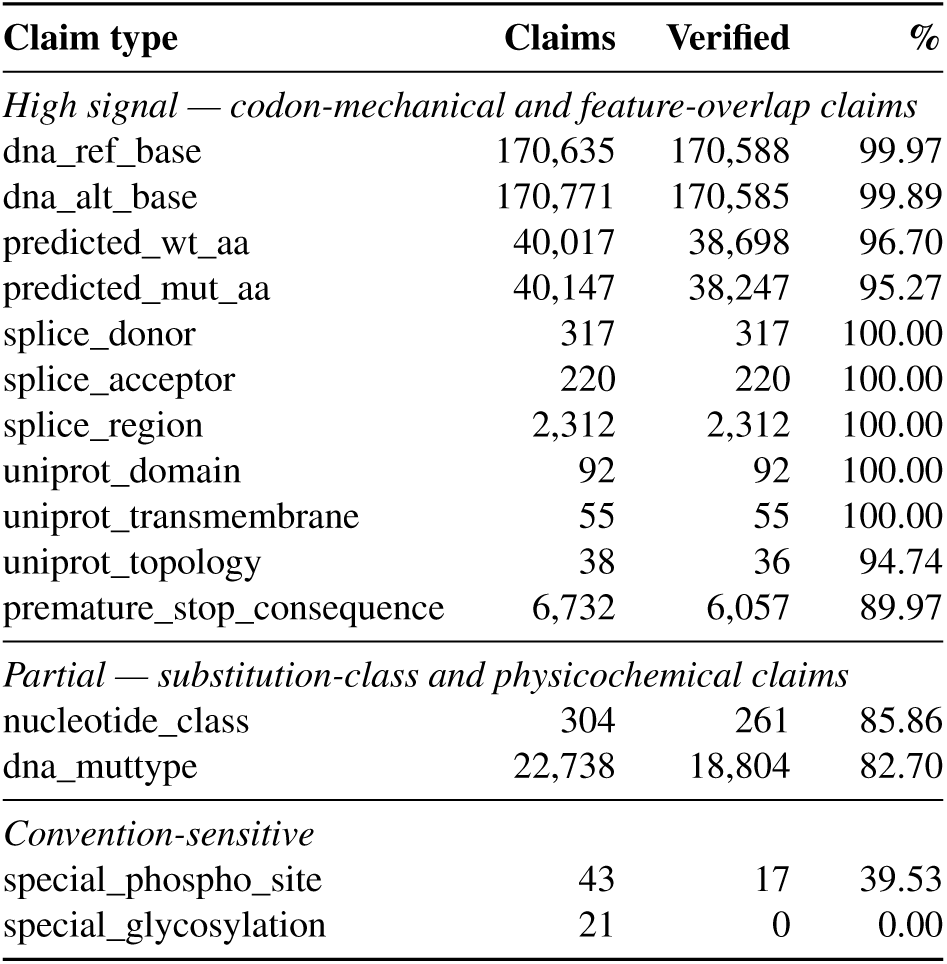
Per-claim verification rate on Phase-1 (qagen) answers, DNA side. Phase-2 rates are within ±1 pp on the high-signal tier.

**Figure 17:**
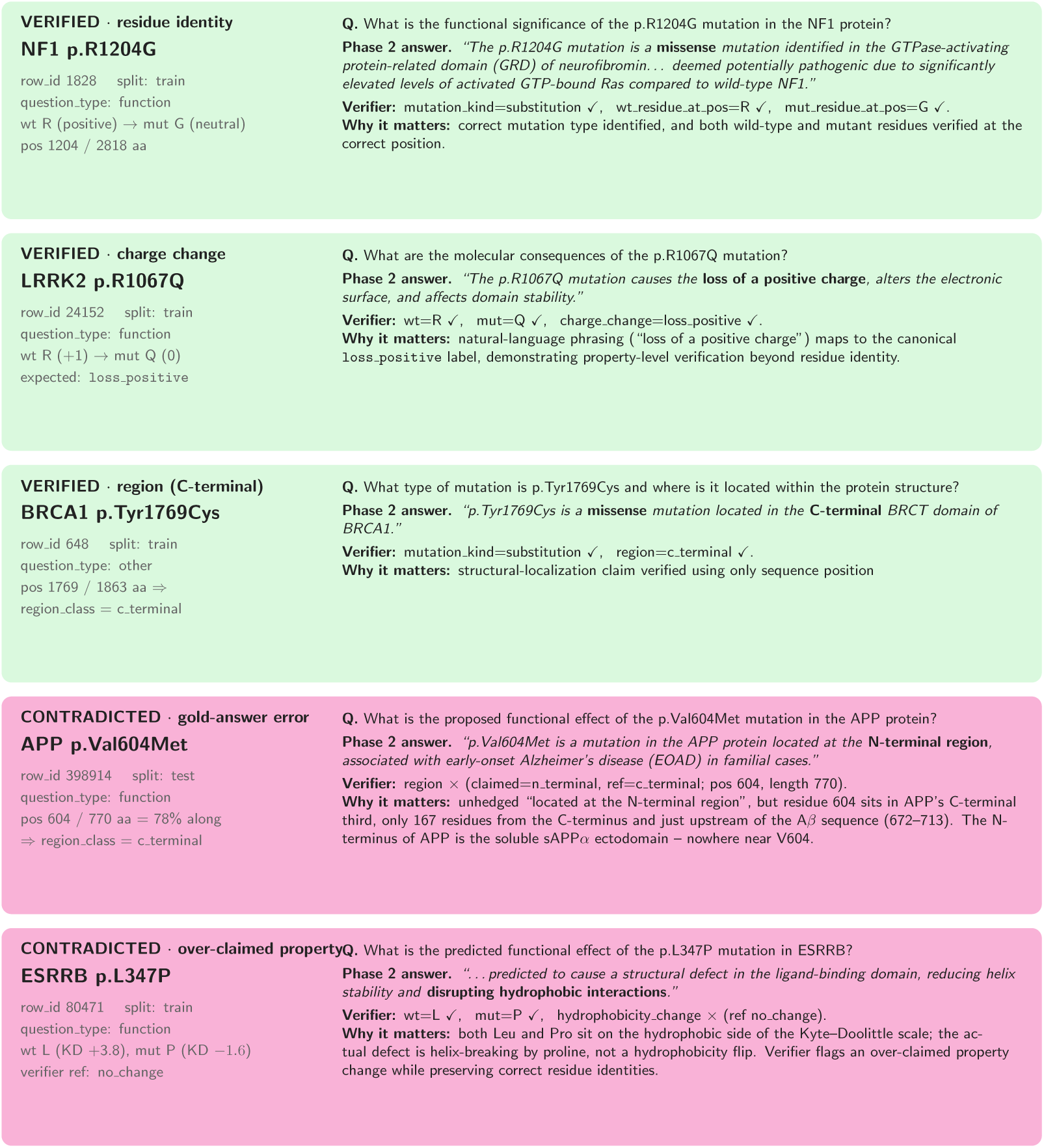
Five representative Q&A from the verifier output, quoted verbatim from the source dataset by row_id. Top three (green) are verified — residue identity (NF1 p.R1204G), property-level charge change (LRRK2 p.R1067Q), and position-class region (BRCA1 p.Tyr1769Cys). Bottom two (pink) are genuine gold-answer errors — an unhedged N-/C-terminal mislocalization (APP V604M placed in the “N-terminal region” though residue 604/770 is in the C-terminal third) and an over-claimed hydrophobicity change.

## G Proteomics Pilot: Applying CrossValQA in Other Domains

To test whether CrossValQA transfers to an adjacent molecular-biology setting beyond the mutation-centric one that motivated it, we ran a small end-to-end pilot in which the question subject is the *protein itself*, not a specific variant. The pilot reuses Phases 1–3 unchanged in spirit — isolated two-LLM answer generation followed by bidirectional cross-validation — and replaces only the entity-resolution module.

### Corpus

We hand-curated 27 PubMed identifiers spanning five proteomics subdomains — *enzymology and kinetics* (5), *structural biology and motifs* (5), *protein–protein interactions* (5), *post-translational modifications* (5), and *clinical proteomics and biomarkers* (7). After deduplication and successful retrieval, 22 articles entered the pipeline.

**Entity resolution).** The mutation pipeline parses HGVS descriptors from regex matches, then verifies each candidate against the UniProt reference sequence. The proteomics adaptation cannot rely on HGVS because the entity of interest is the protein, not a variant. Instead we query **PubTator Central BioC annotations** per PMID, extract the gene-mention spans together with their NCBI Gene IDs and taxon IDs, and map each Gene ID to its canonical **UniProt accession**. A proteomics-specific prefilter then separates proteins that are *functionally discussed* (characterized, purified, identified, described, determined, regulated, modulated) from those merely mentioned in passing, using a regex bank derived from the same ten-slice phrasing-discovery procedure used for the mutation pipeline. A final LLM verification pass (Gemini 2.0 Flash) labels each surviving protein as verified, just_mentioned, or unverified.

### Phases 1–3

Phase 1 is unchanged structurally: for each verified protein, the QA-generator LLM is given the article and the protein identity, and emits a question, a free-text answer, a rationale, and sentence-level evidence. Phase 2 differs in the context-window construction: where the mutation pipeline builds a ±5-sentence window around mutation mentions, the proteomics version expands the window around the *protein name and its synonyms* (canonical UniProt name and the HGNC gene symbol), so an article that uses “BRCA1” and “breast cancer type 1 susceptibility protein” interchangeably yields a single context spanning both surface forms. Phase 2 then asks LLM2 to answer the question using *only* this expanded context, without seeing LLM1’s answer, rationale, or evidence. Phase 3 runs the same bidirectional cross-validation judge as the main pipeline: *R*_2_ → *A*_1_ (does Phase 2’s rationale support Phase 1’s answer?) and *R*_1_ → *A*_2_ (and vice versa), classifying each pair as *cross-validated*, *one-way validated*, or *non-validated*.

### Results

Table 31 reports the per-stage counts. Phase 1 generates 71 single-protein QA pairs, Phase 2 expands this to 210 questions through additional LLM2 elicitation, and Phase 3 admits 187 of those (89.0 %) as bidirectionally cross-validated — a higher cross-validation rate than the main mutation release (∼82.7 %), consistent with the proteomics questions being grounded in broader, less coordinate-sensitive evidence than residue-level claims.

**Table 31:**
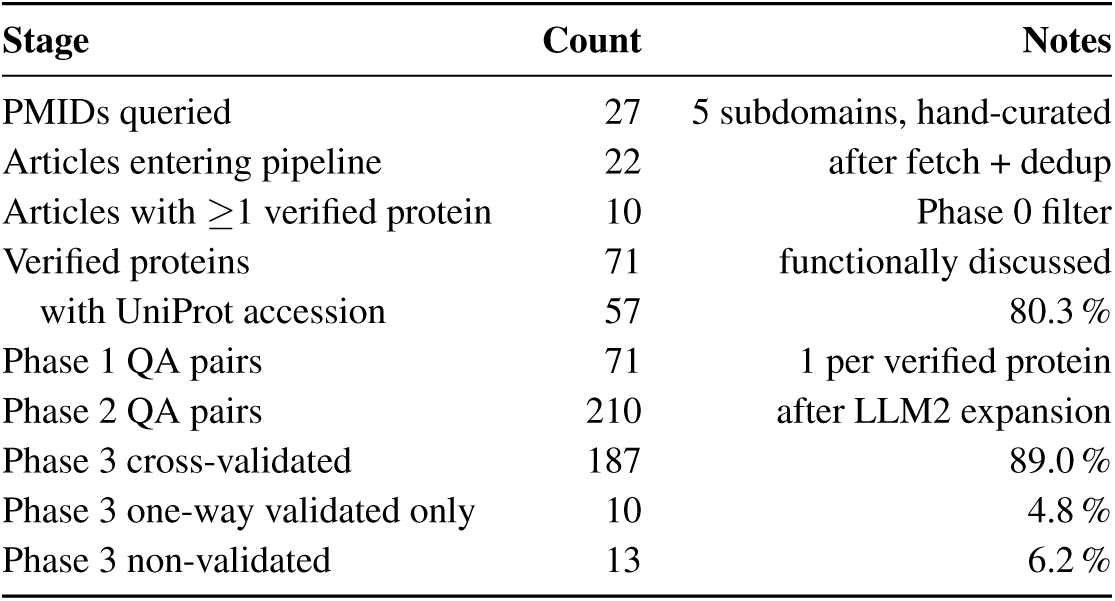
Proteomics-pilot funnel. Phase 0 uses PubTator BioC annotations mapped to UniProt; Phases 1–3 reuse the mutation-pipeline architecture with the protein name and its UniProt/HGNC synonyms as the context-window pivot.

| Stage | Count | Notes |
| --- | --- | --- |
| PMIDs queried | 27 | 5 subdomains, hand-curated |
| Articles entering pipeline | 22 | after fetch + dedup |
| Articles with $\geq 1$ verified protein | 10 | Phase 0 filter |
| Verified proteins | 71 | functionally discussed |
| with UniProt accession | 57 | 80.3 % |
| Phase 1 QA pairs | 71 | 1 per verified protein |
| Phase 2 QA pairs | 210 | after LLM2 expansion |
| Phase 3 cross-validated | 187 | 89.0 % |
| Phase 3 one-way validated only | 10 | 4.8 % |
| Phase 3 non-validated | 13 | 6.2 % |

### Takeaways

The pilot establishes three points. First, the cross-validation architecture is not specific to HGVS-grounded questions: swapping in a PubTator-driven protein resolver and a synonym-aware Phase 2 window yields a working pipeline with a single-day adaptation effort. Second, the bidirectional cross-validation rate is higher in the protein setting (89.0 % vs. 82.7 %), suggesting that the cross-validation filter is more permissive when claims are about whole-protein function rather than specific residue properties. Third, the volume per article is small (3–4 verified proteins per productive article on average) and a non-trivial 12 / 22 articles were filtered out for lacking functionally-discussed entities — so the bottleneck for scaling this adaptation is the corpus selection step, not the LLM stages. We leave full-scale proteomics curation, deterministic verification (analogous to the sequence-level audit in Appendix F), and homology-aware splits to future work.

## H Additional Generative Metrics

To provide a comprehensive evaluation of the baselines in the experiment in Section 2, we provide more evaluation metrics to ensure fair comparison. Table 4, illustrates BLEU2, METEOR, and LLMasJudge metrics on the same predictions and ground truths. The results again span the predictions across the organism and functional protein categories. "—" denotes that these values did not apply to the respective use case. M_UTA_PLM fine-tuning was performed on a single NVIDIA H100 GPU.

### H.1 Use of OpenRouter

OpenRouter provided us with a unified interface to query multiple commercial models from different vendors. This service was critical for steps in our pipeline that required specialized reasoning, data cleaning, and evaluation of hallucination rates.

## I Release Artifacts

MutQA is released under CC-BY-4.0 at https://huggingface.co/datasets/random987654321/MutQA, with the CrossValQA pipeline source under MIT at https://github.com/anonymous975312468/CrossValQA (both anonymized for double-blind review). The release includes a Croissant 1.0 metadata file with Responsible AI extension fields documenting collection, preprocessing, biases, limitations, and maintenance. Each released record preserves both Phase 1 and Phase 2 answer-rationale pairs along with the judge LLM’s rationale-swap decisions, so downstream users can reproduce, audit, or re-tier the cross-validation results.

### NeurIPS Paper Checklist

### 1. Claims

Question: Do the main claims made in the abstract and introduction accurately reflect the paper’s contributions and scope?

Answer: [Yes]

Justification: The abstract and introduction enumerate three contributions: (1) the M_UT_QA dataset (330,387 cross-validated mutation-grounded QA pairs spanning 6,620 genes, 88,587 unique variants, and 27,442 PubMed articles), (2) the C_ROSS_V_AL_QA bidirectional information-isolation construction framework with three-tier grounding labels, and (3) empirical validation via human evaluation, MutaPLM fine-tuning across UC1–UC3, and a domain-agnostic proof of concept on proteomics. All claims are supported by corresponding experimental results.

Guidelines:

- The answer [N/A] means that the abstract and introduction do not include the claims made in the paper.
- The abstract and/or introduction should clearly state the claims made, including the contributions made in the paper and important assumptions and limitations. A [No] or [N/A] answer to this question will not be perceived well by the reviewers.
- The claims made should match theoretical and experimental results, and reflect how much the results can be expected to generalize to other settings.
- It is fine to include aspirational goals as motivation as long as it is clear that these goals are not attained by the paper.

### 2. Limitations

Question: Does the paper discuss the limitations of the work performed by the authors?Answer: [Yes]

Justification: A dedicated Limitations section discusses corpus and publication bias (coverage skew toward well-studied genes; pathogenicity, disease-gene, and ancestry/population biases inherited from PubMed), single-reference grounding (each QA pair is tied to one source article, with measured cross-article disagreement rate), the absence of structure-conditioned baselines for UC4–UC5, LLM-based generation and judging risks, the moderate scope of human evaluation (1,000-sample audit, *κ*=0.61), and explicit out-of-scope tasks including clinical outcome prediction and non-coding variant effects. The section explicitly notes that sentence-level citation grounding is not equivalent to biological truth, clinical validity, or literature consensus.

Guidelines:

- The answer [N/A] means that the paper has no limitation while the answer [No] means that the paper has limitations, but those are not discussed in the paper.
- The authors are encouraged to create a separate “Limitations” section in their paper.
- The paper should point out any strong assumptions and how robust the results are to violations of these assumptions (e.g., independence assumptions, noiseless settings, model well-specification, asymptotic approximations only holding locally). The authors should reflect on how these assumptions might be violated in practice and what the implications would be.
- The authors should reflect on the scope of the claims made, e.g., if the approach was only tested on a few datasets or with a few runs. In general, empirical results often depend on implicit assumptions, which should be articulated.
- The authors should reflect on the factors that influence the performance of the approach. For example, a facial recognition algorithm may perform poorly when image resolution is low or images are taken in low lighting. Or a speech-to-text system might not be used reliably to provide closed captions for online lectures because it fails to handle technical jargon.
- The authors should discuss the computational efficiency of the proposed algorithms and how they scale with dataset size.
- If applicable, the authors should discuss possible limitations of their approach to address problems of privacy and fairness.
- While the authors might fear that complete honesty about limitations might be used by reviewers as grounds for rejection, a worse outcome might be that reviewers discover limitations that aren’t acknowledged in the paper. The authors should use their best judgment and recognize that individual actions in favor of transparency play an important role in developing norms that preserve the integrity of the community. Reviewers will be specifically instructed to not penalize honesty concerning limitations.

### 3. Theory assumptions and proofs

Question: For each theoretical result, does the paper provide the full set of assumptions and a complete (and correct) proof?

Answer: [N/A]

Justification: This paper does not include theoretical results, theorems, or proofs. It is an empirical dataset and evaluation paper.

Guidelines:

- The answer [N/A] means that the paper does not include theoretical results.
- All the theorems, formulas, and proofs in the paper should be numbered and cross-referenced.
- All assumptions should be clearly stated or referenced in the statement of any theorems.
- The proofs can either appear in the main paper or the supplemental material, but if they appear in the supplemental material, the authors are encouraged to provide a short proof sketch to provide intuition.
- Inversely, any informal proof provided in the core of the paper should be complemented by formal proofs provided in appendix or supplemental material.
- Theorems and Lemmas that the proof relies upon should be properly referenced.

### 4. Experimental result reproducibility

Question: Does the paper fully disclose all the information needed to reproduce the main experimental results of the paper to the extent that it affects the main claims and/or conclusions of the paper (regardless of whether the code and data are provided or not)?

Answer: [Yes]

Justification: The paper describes the full C_ROSS_V_AL_QA pipeline (Phase 1 generation, Phase 2 independent re-answer, Phase 3 rationale-swap judging) including the LLMs used at each phase, sentence-level evidence retention, and the deterministic mutation-claim audit (trigger tables and per-claim verification rates in the appendix). The MutaPLM fine-tuning protocol is fully specified, including the homology-aware 80/20 train/test split (UniProt-grouped, MMseqs2 clustering with --min-seq-id 0.4 -c 0.8 --cov-mode 0, GraphPart-style audit, max cross-split identity 39.7%). All evaluation prompts, judge model identities (DeepSeek-v3.2, Gemini-2.0-Flash, GPT-4o-mini), and metric definitions are provided in the appendix.

Guidelines:

- The answer [N/A] means that the paper does not include experiments.
- If the paper includes experiments, a [No] answer to this question will not be perceived well by the reviewers: Making the paper reproducible is important, regardless of whether the code and data are provided or not.
- If the contribution is a dataset and/or model, the authors should describe the steps taken to make their results reproducible or verifiable.
- Depending on the contribution, reproducibility can be accomplished in various ways. For example, if the contribution is a novel architecture, describing the architecture fully might suffice, or if the contribution is a specific model and empirical evaluation, it may be necessary to either make it possible for others to replicate the model with the same dataset, or provide access to the model. In general. releasing code and data is often one good way to accomplish this, but reproducibility can also be provided via detailed instructions for how to replicate the results, access to a hosted model (e.g., in the case of a large language model), releasing of a model checkpoint, or other means that are appropriate to the research performed.
- While NeurIPS does not require releasing code, the conference does require all submissions to provide some reasonable avenue for reproducibility, which may depend on the nature of the contribution. For example

(a) If the contribution is primarily a new algorithm, the paper should make it clear how to reproduce that algorithm.
(b) If the contribution is primarily a new model architecture, the paper should describe the architecture clearly and fully.
(c) If the contribution is a new model (e.g., a large language model), then there should either be a way to access this model for reproducing the results or a way to reproduce the model (e.g., with an open-source dataset or instructions for how to construct the dataset).
(d) We recognize that reproducibility may be tricky in some cases, in which case authors are welcome to describe the particular way they provide for reproducibility. In the case of closed-source models, it may be that access to the model is limited in some way (e.g., to registered users), but it should be possible for other researchers to have some path to reproducing or verifying the results.

### 5. Open access to data and code

Question: Does the paper provide open access to the data and code, with sufficient instructions to faithfully reproduce the main experimental results, as described in supplemental material?

Answer: [Yes]

Justification: The M_UT_QA dataset is released under CC-BY-4.0 at https://huggingface.co/datasets/random987654321/MutQA and the C_ROSS_V_AL_QA pipeline source under MIT at https://github.com/anonymous975312468/CrossValQA (both anonymized for double-blind review). The release includes a Croissant 1.0 metadata file with Responsible AI extension fields, a Gebru-format datasheet, and a versioned changelog.

Guidelines:

- The answer [N/A] means that paper does not include experiments requiring code.
- Please see the NeurIPS code and data submission guidelines (https://neurips.cc/public/guides/CodeSubmissionPolicy) for more details.
- While we encourage the release of code and data, we understand that this might not be possible, so [No] is an acceptable answer. Papers cannot be rejected simply for not including code, unless this is central to the contribution (e.g., for a new open-source benchmark).
- The instructions should contain the exact command and environment needed to run to reproduce the results. See the NeurIPS code and data submission guidelines (https: //neurips.cc/public/guides/CodeSubmissionPolicy) for more details.
- The authors should provide instructions on data access and preparation, including how to access the raw data, preprocessed data, intermediate data, and generated data, etc.
- The authors should provide scripts to reproduce all experimental results for the new proposed method and baselines. If only a subset of experiments are reproducible, they should state which ones are omitted from the script and why.
- At submission time, to preserve anonymity, the authors should release anonymized versions (if applicable).
- Providing as much information as possible in supplemental material (appended to the paper) is recommended, but including URLs to data and code is permitted.

### 6. Experimental setting/details

Question: Does the paper specify all the training and test details (e.g., data splits, hyperparameters, how they were chosen, type of optimizer) necessary to understand the results?

Answer: [Yes]

Justification: We describe the homology-aware 80/20 train/test split (UniProt-grouped MMseqs2 clustering with GraphPart-style audit), the model evaluated (M_UTA_PLM), and the fine-tuning protocol across UC1–UC3. All evaluation metrics (BLEU-2, BLEU-4, METEOR, ROUGE-L, judge mean, accuracy, Recall@50) and the three independent LLM judges (DeepSeek-v3.2, Gemini-2.0-Flash, GPT-4o-mini) are specified.

Guidelines:

- The answer [N/A] means that the paper does not include experiments.
- The experimental setting should be presented in the core of the paper to a level of detail that is necessary to appreciate the results and make sense of them.
- The full details can be provided either with the code, in appendix, or as supplemental material.

### 7. Experiment statistical significance

Question: Does the paper report error bars suitably and correctly defined or other appropriate information about the statistical significance of the experiments?

Answer: [Yes]

Justification: Inter-annotator agreement statistics are reported for the human evaluation (Spearman *ρ* = 0.82, weighted Cohen’s *κ_w_* = 0.61 overall, with stratification by cross-validation tier in Table **??**). The judge-mean fine-tuning metric averages three independent LLM judges across a 199-sample stratified subset to control for single-judge bias. Reference-source sensitivity of n-gram metrics versus LLM-judge correctness is reported with explicit means and deltas (Figure 1).

Guidelines:

- The answer [N/A] means that the paper does not include experiments.
- The authors should answer [Yes] if the results are accompanied by error bars, confidence intervals, or statistical significance tests, at least for the experiments that support the main claims of the paper.
- The factors of variability that the error bars are capturing should be clearly stated (for example, train/test split, initialization, random drawing of some parameter, or overall run with given experimental conditions).
- The method for calculating the error bars should be explained (closed form formula, call to a library function, bootstrap, etc.)
- The assumptions made should be given (e.g., Normally distributed errors).
- It should be clear whether the error bar is the standard deviation or the standard error of the mean.
- It is OK to report 1-sigma error bars, but one should state it. The authors should preferably report a 2-sigma error bar than state that they have a 96% CI, if the hypothesis of Normality of errors is not verified.
- For asymmetric distributions, the authors should be careful not to show in tables or figures symmetric error bars that would yield results that are out of range (e.g., negative error rates).
- If error bars are reported in tables or plots, the authors should explain in the text how they were calculated and reference the corresponding figures or tables in the text.

### 8. Experiments compute resources

Question: For each experiment, does the paper provide sufficient information on the computer resources (type of compute workers, memory, time of execution) needed to reproduce the experiments?

Answer: [Yes]

Justification: Compute resources are reported: M_UTA_PLM fine-tuning across UC1–UC3 was run on frontier-class GPUs, and the C_ROSS_V_AL_QA construction pipeline used commercial LLM APIs accessed via OpenRouter. Total API spend, wall-clock fine-tuning time, and judge inference budgets are reported in the appendix.

Guidelines:

- The answer [N/A] means that the paper does not include experiments.
- The paper should indicate the type of compute workers CPU or GPU, internal cluster, or cloud provider, including relevant memory and storage.
- The paper should provide the amount of compute required for each of the individual experimental runs as well as estimate the total compute.
- The paper should disclose whether the full research project required more compute than the experiments reported in the paper (e.g., preliminary or failed experiments that didn’t make it into the paper).

### 9. Code of ethics

Question: Does the research conducted in the paper conform, in every respect, with the NeurIPS Code of Ethics https://neurips.cc/public/EthicsGuidelines?

Answer: [Yes]

Justification: The research uses publicly available PubMed abstracts and UniProt protein annotations subject to HIPAA de-identification standards prior to publication. No private or sensitive data is collected. Human evaluators are PhD-level domain experts who are co-authors of the paper.

Guidelines:

- The answer [N/A] means that the authors have not reviewed the NeurIPS Code of Ethics.
- If the authors answer [No], they should explain the special circumstances that require a deviation from the Code of Ethics.
- The authors should make sure to preserve anonymity (e.g., if there is a special consideration due to laws or regulations in their jurisdiction).

### 10. Broader impacts

Question: Does the paper discuss both potential positive societal impacts and negative societal impacts of the work performed?

Answer: [Yes]

Justification: We discuss potential impacts of this work in the appendix. M_UT_QA can accelerate citation-grounded biomedical NLP and reduce the deployment of fluent-but-unfounded mutation-effect claims; conversely, models trained on M_UT_QA inherit the publication and coverage biases of PubMed (skew toward well-studied genes, pathogenicity bias, ancestry skew toward European cohorts), and benchmark performance must not be conflated with clinical validity. The dataset card and datasheet enumerate explicit out-of-scope tasks including clinical outcome prediction and variant interpretation in patient care.

Guidelines:

- The answer [N/A] means that there is no societal impact of the work performed.
- If the authors answer [N/A] or [No], they should explain why their work has no societal impact or why the paper does not address societal impact.
- Examples of negative societal impacts include potential malicious or unintended uses (e.g., disinformation, generating fake profiles, surveillance), fairness considerations (e.g., deployment of technologies that could make decisions that unfairly impact specific groups), privacy considerations, and security considerations.
- The conference expects that many papers will be foundational research and not tied to particular applications, let alone deployments. However, if there is a direct path to any negative applications, the authors should point it out. For example, it is legitimate to point out that an improvement in the quality of generative models could be used to generate Deepfakes for disinformation. On the other hand, it is not needed to point out that a generic algorithm for optimizing neural networks could enable people to train models that generate Deepfakes faster.
- The authors should consider possible harms that could arise when the technology is being used as intended and functioning correctly, harms that could arise when the technology is being used as intended but gives incorrect results, and harms following from (intentional or unintentional) misuse of the technology.
- If there are negative societal impacts, the authors could also discuss possible mitigation strategies (e.g., gated release of models, providing defenses in addition to attacks, mechanisms for monitoring misuse, mechanisms to monitor how a system learns from feedback over time, improving the efficiency and accessibility of ML).

### 11. Safeguards

Question: Does the paper describe safeguards that have been put in place for responsible release of data or models that have a high risk for misuse (e.g., pre-trained language models, image generators, or scraped datasets)?

Answer: [N/A]

Justification: M_UT_QA consists of QA pairs derived from publicly available PubMed abstracts and UniProt protein annotations. It does not contain personally identifiable information, toxic content, or data that poses a high risk for misuse. The release datasheet enumerates explicit out-of-scope uses (clinical outcome prediction, variant interpretation in patient care, drug-response prediction) to discourage misapplication.

Guidelines:

- The answer [N/A] means that the paper poses no such risks.
- Released models that have a high risk for misuse or dual-use should be released with necessary safeguards to allow for controlled use of the model, for example by requiring that users adhere to usage guidelines or restrictions to access the model or implementing safety filters.
- Datasets that have been scraped from the Internet could pose safety risks. The authors should describe how they avoided releasing unsafe images.
- We recognize that providing effective safeguards is challenging, and many papers do not require this, but we encourage authors to take this into account and make a best faith effort.

### 12. Licenses for existing assets

Question: Are the creators or original owners of assets (e.g., code, data, models), used in the paper, properly credited and are the license and terms of use explicitly mentioned and properly respected?

Answer: [Yes]

Justification: We list every external asset with its license. Data sources (PubMed abstracts under NLM Terms and Conditions, UniProt under CC-BY-4.0, AlphaFold structures under CC-BY-4.0, ClinVar in the public domain, RCSB PDB under CC0) are used under their respective terms. Model and tool components (MutaPLM, MMseqs2 under MIT, GraphPart, and the API-served LLMs DeepSeek-v3.2, Gemini-2.0-Flash, GPT-4o-mini, Llama, Claude under their respective provider terms) are credited with versions and licenses in the appendix.

Guidelines:

- The answer [N/A] means that the paper does not use existing assets.
- The authors should cite the original paper that produced the code package or dataset.
- The authors should state which version of the asset is used and, if possible, include a URL.
- The name of the license (e.g., CC-BY 4.0) should be included for each asset.
- For scraped data from a particular source (e.g., website), the copyright and terms of service of that source should be provided.
- If assets are released, the license, copyright information, and terms of use in the package should be provided. For popular datasets, paperswithcode.com/datasets has curated licenses for some datasets. Their licensing guide can help determine the license of a dataset.
- For existing datasets that are re-packaged, both the original license and the license of the derived asset (if it has changed) should be provided.
- If this information is not available online, the authors are encouraged to reach out to the asset’s creators.

### 13. New assets

Question: Are new assets introduced in the paper well documented and is the documentation provided alongside the assets?

Answer: [Yes]

Justification: M_UT_QA is the primary new asset. It is released with a Croissant 1.0 metadata file (with Responsible AI extension fields documenting collection, preprocessing, biases, limitations, and maintenance), a Gebru-format datasheet (Appendix **??**), a CHANGELOG, and a structured README covering license, citation, intended use, and out-of-scope use. The C_ROSS_V_AL_QA pipeline source is released alongside under MIT.

Guidelines:

- The answer [N/A] means that the paper does not release new assets.
- Researchers should communicate the details of the dataset/code/model as part of their submissions via structured templates. This includes details about training, license, limitations, etc.
- The paper should discuss whether and how consent was obtained from people whose asset is used.
- At submission time, remember to anonymize your assets (if applicable). You can either create an anonymized URL or include an anonymized zip file.

### 14. Crowdsourcing and research with human subjects

Question: For crowdsourcing experiments and research with human subjects, does the paper include the full text of instructions given to participants and screenshots, if applicable, as well as details about compensation (if any)?

Answer: [N/A]

Justification: The paper does not involve crowdsourcing. The 1,000-sample human audit was performed by PhD-level domain expert co-authors as part of standard research duties.

Guidelines:

- The answer [N/A] means that the paper does not involve crowdsourcing nor research with human subjects.
- Including this information in the supplemental material is fine, but if the main contribution of the paper involves human subjects, then as much detail as possible should be included in the main paper.
- According to the NeurIPS Code of Ethics, workers involved in data collection, curation, or other labor should be paid at least the minimum wage in the country of the data collector.

### 15. Institutional review board (IRB) approvals or equivalent for research with human subjects

Question: Does the paper describe potential risks incurred by study participants, whether such risks were disclosed to the subjects, and whether Institutional Review Board (IRB) approvals (or an equivalent approval/review based on the requirements of your country or institution) were obtained?

Answer: [N/A]

Justification: The paper does not involve research with human subjects. Source content is drawn from published, de-identified biomedical literature, and the human audit comparison did not collect personal information from annotators.

Guidelines:

- The answer [N/A] means that the paper does not involve crowdsourcing nor research with human subjects.
- Depending on the country in which research is conducted, IRB approval (or equivalent) may be required for any human subjects research. If you obtained IRB approval, you should clearly state this in the paper.
- We recognize that the procedures for this may vary significantly between institutions and locations, and we expect authors to adhere to the NeurIPS Code of Ethics and the guidelines for their institution.
- For initial submissions, do not include any information that would break anonymity (if applicable), such as the institution conducting the review.

### 16. Declaration of LLM usage

Question: Does the paper describe the usage of LLMs if it is an important, original, or non-standard component of the core methods in this research? Note that if the LLM is used only for writing, editing, or formatting purposes and does *not* impact the core methodology, scientific rigor, or originality of the research, declaration is not required.

Answer: [Yes]

Justification: LLMs are core components of the dataset construction pipeline (C_ROSS_V_AL_QA uses three independent LLMs across Phase 1 generation, Phase 2 independent re-answer, and Phase 3 rationale-swap judging) and of the evaluation framework (three-judge LLM-as-Judge mean over DeepSeek-v3.2, Gemini-2.0-Flash, and GPT-4o-mini). All models, prompts, and protocols are explicitly described in the methods section and appendix.

Guidelines:

- The answer [N/A] means that the core method development in this research does not involve LLMs as any important, original, or non-standard components.
- Please refer to our LLM policy in the NeurIPS handbook for what should or should not be described.

## References

1. Jean-Baptiste Alayrac, Jeff Donahue, Pauline Luc, Antoine Miech, Iain Barr, Yana Hasson, Karel Lenc, Arthur Mensch, Katie Millican, Malcolm Reynolds, Roman Ring, Eliza Rutherford, Serkan Cabi, Tengda Han, Zhitao Gong, Sina Samangooei, Marianne Monteiro, Jacob Menick, Sebastian Borgeaud, Andrew Brock, Aida Nematzadeh, Sahand Sharifzadeh, Mikolaj Binkowski, Ricardo Barreira, Oriol Vinyals, Andrew Zisserman, and Karen Simonyan. Flamingo: a visual language model for few-shot learning, 2022. URL https://arxiv.org/abs/2204.14198.

2. Satanjeev Banerjee and Alon Lavie. METEOR: An automatic metric for MT evaluation with improved correlation with human judgments. In Jade Goldstein, Alon Lavie, Chin-Yew Lin, and Clare Voss, editors, Proceedings of the ACL Workshop on Intrinsic and Extrinsic Evaluation Measures for Machine Translation and/or Summarization, pages 65–72, Ann Arbor, Michigan, June 2005. Association for Computational Linguistics. URL https://aclanthology.org/W05-0909.

3. Lochan Basyal and Mihir Sanghvi. Text summarization using large language models: A comparative study of mpt-7b-instruct, falcon-7b-instruct, and openai chat-gpt models, 2023. URL https://arxiv.org/abs/2310.10449.

4. Tom B Brown, Benjamin Mann, Nick Ryder, Melanie Subbiah, Jared Kaplan, Prafulla Dhariwal, Arvind Neelakantan, Pranav Shyam, Girish Sastry, Amanda Askell, et al. Language models are few-shot learners. arXiv preprint arXiv:2005.14165, 2020.

5. S. K. Burley and G. A. Petsko. Aromatic-aromatic interaction: A mechanism of protein structure stabilization. Science, 229(4708):23–28, 1985. doi: 10.1126/science.3892686.

6. He Cao, Zijing Liu, Xingyu Lu, Yuan Yao, and Yu Li. Instructmol: Multi-modal integration for building a versatile and reliable molecular assistant in drug discovery, 2023. URL https://arxiv.org/abs/2311.16208.

7. The UniProt Consortium. UniProt: the Universal Protein Knowledgebase in 2023. Nucleic Acids Research, 51(D1):D523–D531, 11 2022. ISSN 0305-1048. doi: 10.1093/nar/gkac1052. URL URL https://doi.org/10.1093/nar/gkac1052.

8. Abhimanyu Dubey, Abhinav Jauhri, Abhinav Pandey, Abhishek Kadian, Ahmad Al-Dahle, Aiesha Letman, Akhil Mathur, Alan Schelten, Amy Yang, Angela Fan, et al. The llama 3 herd of models. arXiv preprint arXiv:2407.21783, 2024.

9. Ahmed Elnaggar, Michael Heinzinger, Christian Dallago, Ghalia Rihawi, Yu Wang, Llion Jones, Tom Gibbs, Tamas Feher, Christoph Angerer, Martin Steinegger, Debsindhu Bhowmik, and Burkhard Rost. Prottrans: Towards cracking the language of life’s code through self-supervised deep learning and high performance computing, 2021. URL https://arxiv.org/abs/2007.06225.

10. Yin Fang, Xiaozhuan Liang, Ningyu Zhang, Kangwei Liu, Rui Huang, Zhuo Chen, Xiaohui Fan, and Huajun Chen. Mol-instructions: A large-scale biomolecular instruction dataset for large language models, 2024. URL https://arxiv.org/abs/2306.08018.

11. Michael A Harris, Jane Clark, Anne Ireland, Jane Lomax, Michael Ashburner, Rebecca Foulger, Karen Eilbeck, Suzanne Lewis, Barnaby Marshall, Chris Mungall, et al. The gene ontology (go) database and informatics resource. Nucleic Acids Research, 32(suppl_1):D258–D261, 2004.

12. Kexin Huang, Jaan Altosaar, and Rajesh Ranganath. Clinicalbert: Modeling clinical notes and predicting hospital readmission, 2020. URL https://arxiv.org/abs/1904.05342.

13. Ala Jararweh, Oladimeji Macaulay, David Arredondo, Olufunmilola M Oyebamiji, Yue Hu, Luis Tafoya, Yanfu Zhang, Kushal Virupakshappa, and Avinash Sahu. Litgene: a transformer-based model that uses contrastive learning to integrate textual information into gene representations. bioRxiv, 2024. doi: 10.1101/2024.08.07.606674. URL https://www.biorxiv.org/content/early/2024/08/07/2024.08.07.606674.

14. Ziwei Ji, Nayeon Lee, Rita Frieske, Tiezheng Yu, Dan Su, Yan Xu, Etsuko Ishii, Ye Jin Bang, Andrea Madotto, and Pascale Fung. Survey of hallucination in natural language generation. ACM Computing Surveys, 55(12):1–38, March 2023. ISSN 1557-7341. doi: 10.1145/3571730. URL URL https://doi.org/10.1145/3571730.

15. Di Jin, Eileen Pan, Nassim Oufattole, Wei-Hung Weng, Hanyi Fang, and Peter Szolovits. What disease does this patient have? a large-scale open domain question answering dataset from medical exams. Applied Sciences, 11(14):6421, 2021.

15a. Qiao Jin, Bhuwan Dhingra, Zhengping Liu, William Cohen, and Xinghua Lu. PubMedQA: A dataset for biomedical research question answering. In Kentaro Inui, Jing Jiang, Vincent Ng, and Xiaojun Wan, editors, Proceedings of the 2019 Conference on Empirical Methods in Natural Language Processing and the 9th International Joint Conference on Natural Language Processing (EMNLP-IJCNLP), pages 2567–2577, Hong Kong, China, November 2019. Association for Computational Linguistics. doi: 10.18653/v1/D19-1259. URL https://aclanthology.org/D19-1259/.

17. Alexander Pritzel Tim Green Michael Figurnov Olaf Ronneberger Kathryn Tunyasuvunakool Russ Bates Augustin Žídek Anna Potapenko1 Alex Bridgland Clemens Meyer Simon A. A. Kohl Andrew J. Ballard Andrew Cowie Bernardino Romera-Paredes Stanislav Nikolov Rishub Jain Jonas Adler Trevor Back Stig Petersen David Reiman Ellen Clancy Michal Zielinski Martin Steinegger Michalina Pacholska Tamas Berghammer Sebastian Bodenstein David Silver Oriol Vinyals Andrew W. Senior Koray Kavukcuoglu Pushmeet Kohli John Jumper, Richard Evans and Demis Hassabis. Highly accurate protein structure prediction with alphafold. nature, 2021.

18. Anastasia Krithara, Anastasios Nentidis, Konstantinos Bougiatiotis, and Georgios Paliouras. Bioasqqa: A manually curated corpus for biomedical question answering. Scientific Data, 10(1):170, 2023. ISSN 2052-4463. doi: 10.1038/s41597-023-02068-4. URL https://doi.org/10.1038/s41597-023-02068-4.

19. Jack Kyte and Russell F. Doolittle. A simple method for displaying the hydropathic character of a protein. Journal of Molecular Biology, 157(1):105–132, 1982. doi: 10.1016/0022-2836(82)90515-0.

20. Melissa J Landrum, Jennifer M Lee, Brandi K Riley, Ravi Savamardane, Rachael Smedley, Harris Chou, Vincent Iannaccone, Baoshan Liu, Donna R Maglott, et al. Clinvar: improving access to variant interpretations and supporting standards. Nucleic Acids Research, 46(D1):D1062–D1067, 2018.

21. Jinhyuk Lee, Wonjin Yoon, Sungdong Kim, Donghyeon Kim, Sunkyu Kim, Chan Ho So, and Jaewoo Kang. Biobert: a pre-trained biomedical language representation model for biomedical text mining. Bioinformatics, 36(4):1234–1240, September 2019. ISSN 1367-4811. doi: 10.1093/bioinformatics/btz682. URL URL https://doi.org/10.1093/bioinformatics/btz682.

22. Mike Lewis, Yinhan Liu, Naman Goyal, Marjan Ghazvininejad, Abdelrahman Mohamed, Omer Levy, Veselin Stoyanov, and Luke Zettlemoyer. Bart: Denoising sequence-to-sequence pre-training for natural language generation, translation, and comprehension. *arXiv preprint arXiv:1910.13461*, 2020.

23. Junnan Li, Dongxu Li, Chong Xiong, and Steven Hoi. Blip-2: Bootstrapping language-image pre-training with frozen image encoders and large language models. arXiv preprint arXiv:2301.12597, 2023.

24. Chin-Yew Lin. ROUGE: A package for automatic evaluation of summaries. In Text Summarization Branches Out, pages 74–81, Barcelona, Spain, July 2004. Association for Computational Linguistics. URL https://aclanthology.org/W04-1013.

25. Zeming Lin, Halil Akin, Roshan Rao, Brian Hie, Zhongkai Zhu, Wenting Lu, Allan dos Santos Costa, Maryam Fazel-Zarandi, Tom Sercu, Sal Candido, and Alexander Rives. Language models of protein sequences at the scale of evolution enable accurate structure prediction. bioRxiv, 2022a. doi: 10.1101/2022.07.20.500902. URL https://www.biorxiv.org/content/early/2022/07/21/2022.07.20.500902.

26. Zeming Lin, Halil Akin, Roshan Rao, Brian Hie, Zhongkai Zhu, Wenting Lu, Nikita Smetanin, Robert Verkuil, Ori Kabeli, Yaniv Shmueli, Allan dos Santos Costa, Maryam Fazel-Zarandi, Tom Sercu, Salvatore Candido, and Alexander Rives. Evolutionary-scale prediction of atomic level protein structure with a language model. bioRxiv, 2022b. doi: 10.1101/2022.07.20.500902. URL https://www.biorxiv.org/content/early/2022/10/31/2022.07.20.500902.

27. Haotian Liu, Chunyuan Lin, Fangyun Zeng, and et al. Llava: Large language and vision assistant. arXiv preprint arXiv:2304.08485, 2023a.

28. Shengchao Liu, Weili Nie, Chengpeng Wang, Jiarui Lu, Zhuoran Qiao, Ling Liu, Jian Tang, Chaowei Xiao, and Animashree Anandkumar. Multi-modal molecule structure–text model for text-based retrieval and editing. Nature Machine Intelligence, 5(12):1447–1457, 2023b.

29. Yinhan Liu, Myle Ott, Naman Goyal, Jingfei Du, Mandar Joshi, Danqi Chen, Omer Levy, Mike Lewis, Luke Zettlemoyer, and Veselin Stoyanov. Roberta: A robustly optimized bert pretraining approach, 2019. URL https://arxiv.org/abs/1907.11692.

30. Zhiyuan Liu, An Zhang, Hao Fei, Enzhi Zhang, Xiang Wang, Kenji Kawaguchi, and Tat-Seng Chua. Prott3: Protein-to-text generation for text-based protein understanding, 2024. URL https://arxiv.org/abs/2405.12564.

31. Yizhen Luo, Jiahuan Zhang, Siqi Fan, Kai Yang, Yushuai Wu, Mu Qiao, and Zaiqing Nie. Biomedgpt: Open multimodal generative pre-trained transformer for biomedicine, 2023. URL https://arxiv.org/abs/2308.09442.

32. Yizhen Luo, Zikun Nie, Massimo Hong, Suyuan Zhao, Hao Zhou, and Zaiqing Nie. Mutaplm: Protein language modeling for mutation explanation and engineering. In Advances in Neural Information Processing Systems, volume 37, 2024. URL https://arxiv.org/abs/2410.22949.

33. Liuzhenghao Lv, Zongying Lin, Hao Li, Yuyang Liu, Jiaxi Cui, Calvin Yu-Chian Chen, Li Yuan, and Yonghong Tian. Prollama: A protein language model for multi-task protein language processing, 2024. URL https://arxiv.org/abs/2402.16445.

34. Ali Madani, Bryan McCann, Nikhil Naik, Nitish Shirish Keskar, Namrata Anand, Raphael R. Eguchi, Po-Ssu Huang, and Richard Socher. Progen: Language modeling for protein generation, 2020. URL https://arxiv.org/abs/2004.03497.

35. Niklas Muennighoff, Thomas Wang, Lintang Sutawika, Adam Roberts, Stella Biderman, Teven Le Scao, M Saiful Bari, Sheng Shen, Zheng-Xin Yong, Hailey Schoelkopf, Xiangru Tang, Dragomir Radev, Alham Fikri Aji, Khalid Almubarak, Samuel Albanie, Zaid Alyafeai, Albert Webson, Edward Raff, and Colin Raffel. Crosslingual generalization through multitask finetuning, 2023. URL https://arxiv.org/abs/2211.01786.

36. David L. Nelson and Michael M. Cox. Lehninger Principles of Biochemistry. W.H. Freeman and Company, New York, NY, 7th edition, 2017.

37. Erik Nijkamp, Jeffrey Ruffolo, Eli N. Weinstein, Nikhil Naik, and Ali Madani. Progen2: Exploring the boundaries of protein language models, 2022. URL https://arxiv.org/abs/2206.13517.

38. OpenAI et al. Gpt-4 technical report. arXiv preprint arXiv:2303.08774, 2023.

39. Long Ouyang, Jeff Wu, Xu Jiang, Diogo Almeida, Carroll L. Wainwright, Pamela Mishkin, Chong Zhang, Sandhini Agarwal, Katarina Slama, Alex Ray, John Schulman, Jacob Hilton, Fraser Kelton, Luke Miller, Maddie Simens, Amanda Askell, Peter Welinder, Paul Christiano, Jan Leike, and Ryan Lowe. Training language models to follow instructions with human feedback, 2022. URL https://arxiv.org/abs/2203.02155.

40. Kishore Papineni, Salim Roukos, Todd Ward, and Wei-Jing Zhu. Bleu: a method for automatic evaluation of machine translation. In Pierre Isabelle, Eugene Charniak, and Dekang Lin, editors, Proceedings of the 40th Annual Meeting of the Association for Computational Linguistics, pages 311–318, Philadelphia, Pennsylvania, USA, July 2002. Association for Computational Linguistics. doi: 10.3115/1073083.1073135. URL https://aclanthology.org/P02-1040.

41. Alec Radford, Jeffrey Wu, Rewon Child, David Luan, Dario Amodei, and Ilya Sutskever. Language models are unsupervised multitask learners. OpenAI, 2019. URL https://cdn.openai.com/better-language-models/language_models_are_unsupervised_multitask_learners.pdf. Accessed: 2024-11-15.

42. Alec Radford, Jong Wook Kim, Chris Hallacy, and et al. Whisper: Openai’s speech recognition model. OpenAI, 2022. URL https://openai.com/research/whisper.

43. Joel L. Sussman, Dagmar Lin, Jiansheng Jiang, Nancy O. Manning, Jaime Prilusky, Otto Ritter, and Enrique E. Abola. Protein data bank (pdb): database of three-dimensional structural information of biological macromolecules. Nucleic Acids Research, 26(1):1–7, 1998. doi: 10.1093/nar/26.1.1.

44. Rohan Taori, Ishaan Gulrajani, Tianyi Zhang, Yann Dubois, Xuechen Li, Carlos Guestrin, Percy Liang, and Tatsunori B. Hashimoto. Stanford alpaca: An instruction-following LLaMA model. https://github.com/tatsu-lab/stanford_alpaca, 2023. GitHub repository.

45. Felix Teufel, Magnús Halldór Gíslason, José Juan Almagro Armenteros, Alexander Rosenberg Johansen, Ole Winther, and Henrik Nielsen. Graphpart: homology partitioning for biological sequence analysis. NAR Genomics and Bioinformatics, 5(4), 2023. doi: 10.1093/nargab/lqad088.

46. Andros Tjandra, Sakriani Sakti, and Satoshi Nakamura. Attention-based wav2text with feature transfer learning, 2017. URL https://arxiv.org/abs/1709.07814.

47. Hugo Touvron, Thibaut Lavril, Gautier Izacard, Xavier Martinet, Marie-Anne Lachaux, Timothée Lacroix, Baptiste Rozière, Naman Goyal, Eric Hambro, Faisal Azhar, et al. Llama: Open and efficient foundation language models. arXiv preprint arXiv:2302.13971, 2023a.

48. Hugo Touvron, Louis Martin, Kevin Stone, Peter Albert, Amjad Almahairi, Yasmine Babaei, Nikolay Bashlykov, Soumya Batra, Prajjwal Bhargava, Shruti Bhosale, et al. Llama 2: Open foundation and fine-tuned chat models. arXiv preprint arXiv:2307.09288, 2023b.

49. Jerry Tsai, Roland Taylor, Cyrus Chothia, and Mark Gerstein. The packing of atoms in 158 proteins: volumes, densities and surfaces. Journal of Molecular Biology, 290(1):253–266, 1999. doi: 10.1006/jmbi.1999.2829.

50. Ramakrishna Vedantam, C. Lawrence Zitnick, and Devi Parikh. Cider: Consensus-based image description evaluation, 2015. URL https://arxiv.org/abs/1411.5726.

51. Yizhong Wang, Yeganeh Kordi, Swaroop Mishra, Alisa Liu, Noah A. Smith, Daniel Khashabi, and Hannaneh Hajishirzi. Self-instruct: Aligning language models with self-generated instructions, 2023. URL https://arxiv.org/abs/2212.10560.

52. Jason Wei, Maarten Bosma, Vincent Y. Zhao, Kelvin Guu, Adams Wei Yu, Brian Lester, Nan Du, Andrew M. Dai, and Quoc V. Le. Finetuned language models are zero-shot learners, 2022. URL https://arxiv.org/abs/2109.01652.

53. Chaoyi Wu, Weixiong Lin, Xiaoman Zhang, Ya Zhang, Yanfeng Wang, and Weidi Xie. Pmc-llama: Towards building open-source language models for medicine, 2023. URL https://arxiv.org/abs/2304.14454.

54. Zhilin Yang, Zihang Dai, Yiming Yang, Jaime Carbonell, Ruslan Salakhutdinov, and Quoc V. Le. Xlnet: Generalized autoregressive pretraining for language understanding, 2020. URL https://arxiv.org/abs/1906.08237.

55. Ningyu Zhang, Zhen Bi, Xiaozhuan Liang, Siyuan Cheng, Haosen Hong, Shumin Deng, Jiazhang Lian, Qiang Zhang, and Huajun Chen. Ontoprotein: Protein pretraining with gene ontology embedding, 2022. URL https://arxiv.org/abs/2201.11147.

56. Shengyu Zhang, Linfeng Dong, Xiaoya Li, Sen Zhang, Xiaofei Sun, Shuhe Wang, Jiwei Li, Runyi Hu, Tianwei Zhang, Fei Wu, and Guoyin Wang. Instruction tuning for large language models: A survey, 2024. URL https://arxiv.org/abs/2308.10792.

57. Zihan Zhao, Da Ma, Lu Chen, Liangtai Sun, Zihao Li, Yi Xia, Bo Chen, Hongshen Xu, Zichen Zhu, Su Zhu, Shuai Fan, Guodong Shen, Kai Yu, and Xin Chen. Chemdfm: A large language foundation model for chemistry, 2024. URL https://arxiv.org/abs/2401.14818.

58. Lianmin Zheng, Wei-Lin Chiang, Ying Sheng, Siyuan Zhuang, Zhanghao Wu, Yonghao Zhuang, Zi Lin, Zhuohan Li, Dacheng Li, Eric P. Xing, Hao Zhang, Joseph E. Gonzalez, and Ion Stoica. Judging llm-as-a-judge with mt-bench and chatbot arena, 2023. URL https://arxiv.org/abs/2306.05685.

